# Programmable RNA writing with trans-splicing

**DOI:** 10.1101/2024.01.31.578223

**Authors:** Cian Schmitt-Ulms, Alisan Kayabolen, Marcos Manero-Carranza, Nathan Zhou, Keira Donnelly, Sabrina Pia Nuccio, Kazuki Kato, Hiroshi Nishimasu, Jonathan S. Gootenberg, Omar O. Abudayyeh

## Abstract

RNA editing offers the opportunity to introduce either stable or transient modifications to nucleic acid sequence without permanent off-target effects, but installation of arbitrary edits into the transcriptome is currently infeasible. Here, we describe Programmable RNA Editing & Cleavage for Insertion, Substitution, and Erasure (PRECISE), a versatile RNA editing method for writing RNA of arbitrary length and sequence into existing pre-mRNAs via 5′ or 3′ trans-splicing. In trans-splicing, an exogenous template is introduced to compete with the endogenous pre-mRNA, allowing for replacement of upstream or downstream exon sequence. Using Cas7-11 cleavage of pre-mRNAs to bias towards editing outcomes, we boost the efficiency of RNA trans-splicing by 10–100 fold, achieving editing rates between 5–50% and 85% on endogenous and reporter transcripts, respectively, while maintaining high-fidelity. We demonstrate PRECISE editing across 11 distinct endogenous transcripts of widely varying expression levels, showcasing more than 50 types of edits, including all 12 possible transversions and transitions, insertions ranging from 1 to 1,863 nucleotides, and deletions. We show high efficiency replacement of exon 4 of MECP2, addressing most mutations that drive the Rett Syndrome; editing of SHANK3 transcripts, a gene involved in Autism; and replacement of exon 1 of HTT, removing the hallmark repeat expansions of Huntington′s disease. Whole transcriptome sequencing reveals the high precision of PRECISE editing and lack of off-target trans-splicing activity. Furthermore, we combine payload engineering and ribozymes for protein-free, high-efficiency trans-splicing, with demonstrated efficiency in editing HTT exon 1 via AAV delivery. We show that the high activity of PRECISE editing enables editing in non-dividing neurons and patient-derived Huntington’s disease fibroblasts. PRECISE editing markedly broadens the scope of genetic editing, is straightforward to deliver over existing gene editing tools like prime editing, lacks permanent off-targets, and can enable any type of genetic edit large or small, including edits not otherwise possible with existing RNA base editors, widening the spectrum of addressable diseases.

## Main Text

The programmable manipulation of genetic information in living cells and organisms has massive implications in both basic research and therapeutics. Gene editing tools for programmable enzymatic modification of DNA, including base editors^1–3^, prime editing^4^, and insertion tools^5–7^, have made significant progress, but these tools are large and bulky, cannot install arbitrary edits, leaving many of the 141,342 known pathogenic mutations unaddressed, and have the risk of permanent off-targets^8^ and bystander edits^9^, potentially limiting clinical utility. In contrast, RNA editors are typically simpler and easier to deliver to diverse tissues and have no risk of permanent off-targets^10–12^. However, mature RNA editing technologies are currently limited, and can only effect two base transitions, (A to I or C to U)^13–22^, leaving the ten other possible base transitions and transversions, as well as small or large insertions or deletions, completely unaddressed. Furthermore, both DNA and RNA editing approaches rely on large protein cargos, often precluding the use of common delivery vectors, such as adeno associated viruses (AAVs). Alternatively, trans-splicing based RNA editing approaches can replace exons for flexible edits, but have been traditionally hampered by low efficiencies^23–30^. As transversions, insertions, and deletions account for a majority of pathogenic mutations^31^, having the means to efficiently install these changes across any transcript in any cell type with viral or non-viral delivery is critical.

Here, we develop an RNA writing technology that enables programmable editing of RNA for all possible single edit base transitions and transversions, any sized deletion, or any sized insertion. Our approach, Programmable RNA Editing & Cleavage for Insertion, Substitution, and Erasure (PRECISE), is capable of both 5′ trans-splicing and 3′ trans-splicing, enabling the replacement of one to more exons of a target transcript efficiently and with high specificity. For 3′ trans-splicing, PRECISE editing employs a programmable RNase to separate cis exons from the pre-mRNA, promoting trans-splicing of an engineered trans-template. For 5′ trans-splicing, cleavage of the poly(A) tail of the trans-template by either programmable RNases or engineered ribozymes increases splicing efficiency; when using ribozymes, only the RNA trans-template is needed, with no exogenous proteins required offering the simplest approach for programmable RNA writing. As PRECISE editing is smaller and less complex than prime editors, base editors, and PASTE, 5′ PRECISE systems can be packaged as a single-vector AAV or lentivirus, achieving up to 15% editing efficiency in HEK293FT cells. Evaluating PRECISE editing in primary cells, we find 5% editing in primary human iPSC-derived neurons and 4% editing in HD patient derived fibroblasts. With the capacity to install any type of single-base edit and any sized deletion or insertion into RNA, PRECISE editing has the potential to advance the study and treatment of a majority of pathogenic genetic variants.

## Results

### PRECISE editing writes sequences into pre-mRNAs with Cas7-11 assisted trans-splicing

To overcome inefficiencies with previously trans-splicing-based RNA editing approaches ^24–30^, we reasoned that precise cleavage of pre-mRNAs could separate the downstream cis exons, biasing competition towards the trans-splicing template and increasing trans-splicing efficiency. We elected to precisely cleave pre-mRNAs using the recently described CRISPR-Cas7-11 nuclease ^32–36^, which generates specific cuts in target RNAs, resulting in the complete PRECISE system (**Fig. 1a**). Since cleavage with PRECISE occurs in intronic regions, overall transcript levels should be unperturbed, as total splicing would still be maintained between cleaved pre-mRNA and the endogenous cis exons or the provided trans-template. We tested the feasibility of PRECISE editing on a luciferase reporter from a Gaussia luciferase (Gluc), containing an open-reading frame split across two exons with the second exon interrupted with a pre-termination stop codon. 3′ trans-splicing via PRECISE introduces a repaired exon without this stop codon, restoring luciferase activity. We designed sequences to hybridize the template with the pre-mRNA, referred to as the template guide RNA (tgRNA), and a CRISPR RNA (crRNA) to direct *Desulfonema ishimotonii* Cas7-11 (*Dis*Cas7-11) to cleave the intervening sequence. In the trans-template cargo, we introduced sequences essential to splicing: the branching point (BP), splicing signal (SS), and polypyrimidine tract (PPT). Delivering a targeting crRNA, *Dis*Cas7-11, and trans-template cargo to HEK293FT cells rescued Gluc production and produced trans-edited mRNA, as measured by next generation sequencing (NGS) **(Fig. 1b-c)**. Critically, editing required Cas7-11 targeting, as substitution of a non-targeting crRNA reduced protein production by 20-fold and editing by 40-fold. Furthermore, editing required BP, SS, and PPT sequences, and omission of any of these sequences drastically reduced editing **(Fig. 1b-c)**. To further explore the dependence of editing of crRNA and tgRNA selection, we tested a panel of crRNA and tgRNA locations on our reporter transcripts. We found that crRNA 1 with cargo tgRNA 2 resulted in the highest levels of correction, boosting Gluc correction by 7.2-fold over the previous best combination (**Extended Data Fig. 1a-d**). As seen before, efficient 3′ trans-splicing with PRECISE editing with these new *Dis*Cas7-11 crRNA and cargo tgRNA combinations was only possible when all three of the SS, BP, and PP signals were present on the trans-template (**Extended Data Fig. 1a-d**).

**Figure 1.**
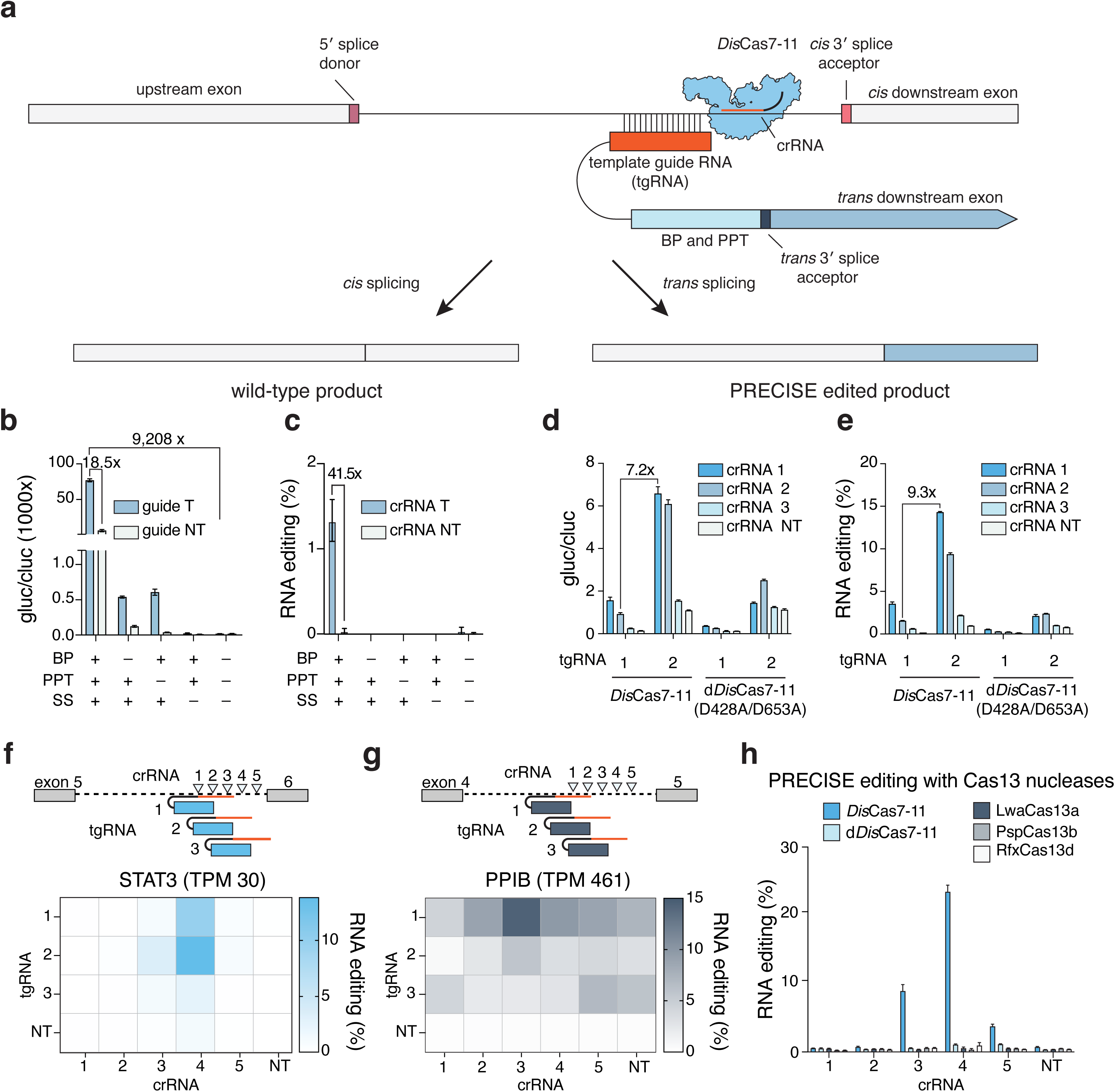
Development of PRECISE RNA editing. **(a)** Schematic showing 3′ PRECISE editing with a trans-splicing cargo and Cas7-11 crRNA targeted to a 3′ region of the intron. BP, branch point; PPT, polypyrimidine tract. **(b)** Splicing assay employing a split-luciferase reporter reconstituting a *Gaussia* luciferase transcript through PRECISE RNA editing, contrasting 3′ cargos with and without a branching point (BP), polypyrimidine tract (PPT), or splicing signal (SS). A *Dis*Cas7-11 targeting guide is compared to a non-targeting guide. *Gaussia* luciferase values normalized to a constitutively co-expressed *Cypridina* luciferase control. **(c)** RNA editing analysis of the splicing assay, using a barcoded cargo and sequencing primers spanning the splice junction. **(d)** Splicing assay employing a split-luciferase reporter reconstituting a *Gaussia* luciferase transcript through PRECISE RNA editing demonstrating three different *Dis*Cas7-11 targeting guides compared against a non-targeting guide and two different cargo hybridization guides (1 and 2). *Gaussia* luciferase values normalized to constitutively co-expressed *Cypridina* luciferase control. Wt denotes wild-type *Dis*Cas7-11. Dead denotes catalytically inactive *Dis*Cas7-11. **(e)** RNA editing analysis for the splicing assay, using a barcoded cargo and sequencing primers spanning the splice junction. **(f)** Endogenous PRECISE editing for a set of crossed guides and cargos targeting the STAT3 pre-mRNA. Editing rates expressed as % of reads containing “barcoding” mutations / total reads of “barcoded” and wild-type transcripts. NT guide, scrambled crRNA; NT cargo, scrambled cargo guide. **(g)** Endogenous PRECISE editing for a set of crossed guides and cargos targeting the PPIB pre-mRNA. Editing rates expressed as % of reads containing “barcoding” mutations / total reads of “barcoded” and wildtype transcripts. NT guide, scrambled crRNA; NT cargo, scrambled hybridization. **(h)** Comparison of splicing rates for a set of cargos targeting STAT3 with either Cas7-11 or Cas13 crRNAs targeting the same STAT3 pre-mRNA intron. *Psp, Prevotella sp.* Cas13; *Lwa*, *Leptotrichia wadei* Cas13; *Ruminococcus flavifaciens* Cas13. All constructs are cloned with 5′ and 3′ flanking SV40 bipartite NLS sequences.

### PRECISE editing requires *Dis*Cas7-11 catalytic activity and is functional on and endogenous transcripts

After initial demonstration of PRECISE editing on the Gluc reporter, we evaluated the dependency of editing on *Dis*Cas7-11 cleavage **(Fig. 1d**). To confirm that the RNA editing rate was similarly improved as with the Gluc protein correction, we sequenced transcripts from cells edited with these constructs and found that *Dis*Cas7-11 crRNA 1 in combination with tgRNA 2 resulted in a 9.3-fold improvement in editing over the previous best combination with ~15% RNA editing, and that this editing was required known catalytic sites of *Dis*Cas7-11 **(Fig. 1e).** RNA editing rates also nearly perfectly correlated with the level of Gluc luciferase activity observed, as expected **(Fig. 1d-e)**.

We next evaluated PRECISE editing on two endogenous transcripts, STAT3 and PPIB. We designed a panel of three cargo hybridization regions and five *Dis*Cas7-11 crRNAs targeting locations around the branch point of intron 4 of PPIB or intron 4 of STAT3 pre-mRNA transcripts. We sequenced the mRNA transcripts and found that editing efficiency was heavily dependent on the positions of both the cargo tgRNA and *Dis*Cas7-11 crRNAs. With optimal positioning and cargo optimization, editing efficiencies as high as 13.8% and 14.4% could be achieved for STAT3 and PPIB, respectively (**Fig. 1f-g and Extended Data Fig. 2–3**). As observed with Gluc, the *Dis*Cas7-11 crRNAs closest to the branching point resulted in the highest level of trans-splicing, suggesting that interfering with this splicing regulatory site is critical for efficient RNA writing. In agreement with this, catalytically inactive d*Dis*Cas7-11-based trans-splicing activity was higher than without d*Dis*Cas7-11, likely due to blocking of the branch point, and trans-splicing was highest when combined with an active *Dis*Cas7-11 nuclease to allow for removal of cis exons (**Extended Data Fig. 3a-b**). We also explored different cargo tgRNA lengths, and found that hybridization lengths of 150 nt yielded the highest trans-splicing activity with PRECISE editing (**Extended Data Fig. 2a-d**). To further optimize PRECISE, we placed linkers of various lengths between the tgRNA and the regulatory signals (BP, SS, and PPT). Similar to other cargo elements, we found that the most efficient linker length varied between the targets tested, potentially due to differences in intron sizes or RNA secondary structures. While this may represent another orthogonal angle for optimizing a specific target, we ultimately selected a 14-nt linker as a consistent option for cargo designs moving forward **(Extended Data Fig. 2e-f)**.

**Figure 2.**
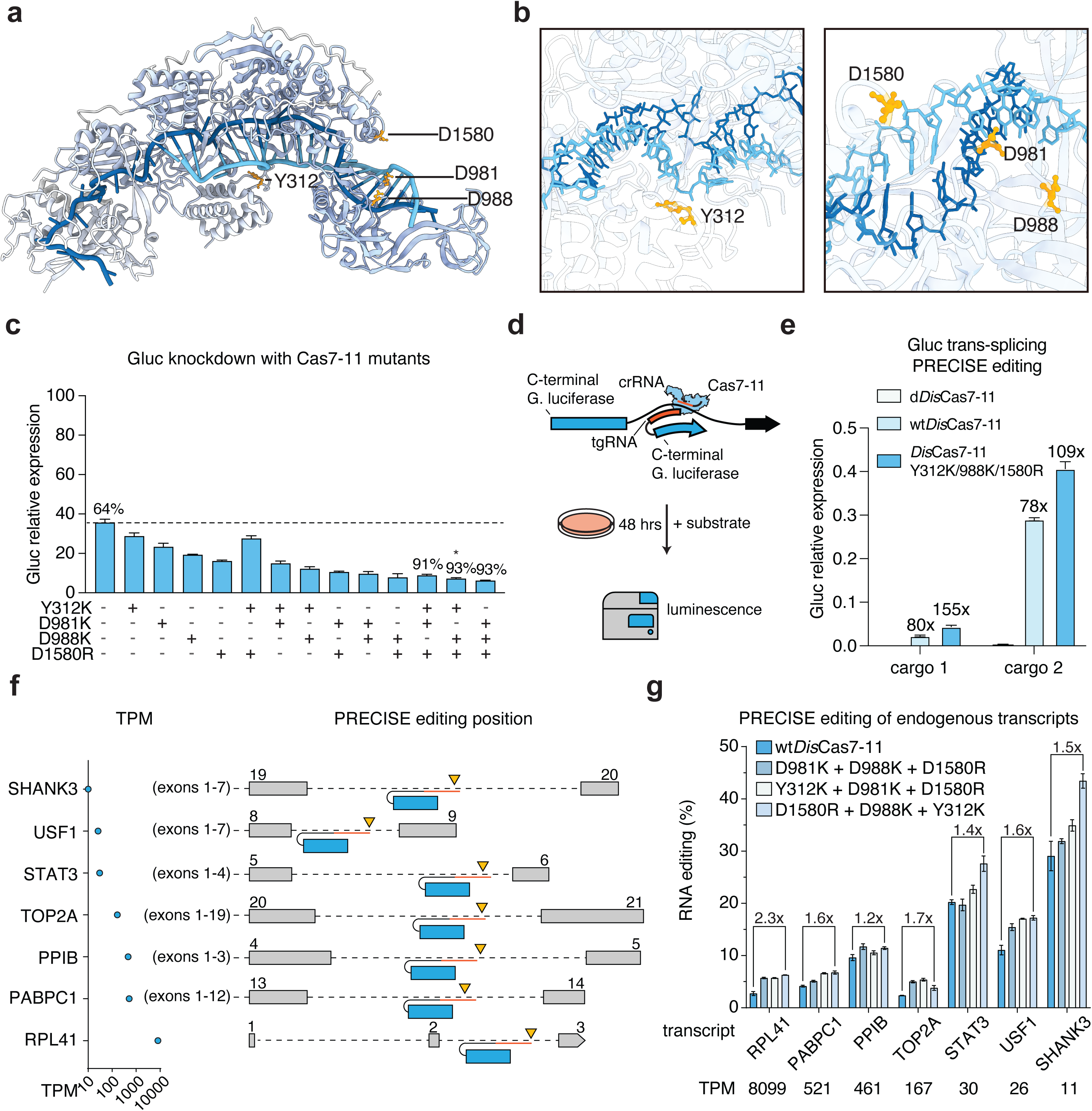
Rational engineering of *Dis*Cas7-11 catalytic activity to improve PRECISE editing. **(a)** Structure of *Dis*Cas7-11 complexed with a crRNA (light blue) and tgRNA (dark blue) showing the positions of residues targeted for mutagenesis for improving RNase activity. Residues with mutations causing a large increase in cleavage efficiency are highlighted (yellow). **(b)** Zooms showing the positions of residues selected for mutagenesis to generate enhanced Cas7-11 catalytic variants. **(c)** Knockdown efficiency of a panel of single, double, and triple mutants of *Dis*Cas7-11 expressed as a ratio between *Gaussia* (targeted) and *Cypridina* luciferase (constitutive) expression, normalized to the ratio for a non-targeting guide condition. The dotted line indicates *Gaussia* luciferase expression levels with knockdown by wild-type *Dis*Cas7-11. The knockdown efficiency as a percentage is denoted above select conditions. **(d)** Schematic showing the construction of a splicing reporter for PRECISE editing, showing the N-terminal domain of *Gaussia* luciferase (blue) in exon 1 with the intron 46 from *COL7A1* gene. The Gluc pre-mRNA is expressed and the PRECISE cargo (blue) is shown bound to the intron, carrying the remaining C-terminal sequence of *Gaussia* luciferase in another exon. **(e)** Splicing assay employing a split-luciferase reporter reconstituting a *Gaussia* luciferase transcript through PRECISE, using two cargos with different cargo hybridization guides and either catalytically inactive (d*Dis*Cas7-11), wild-type *Dis*Cas7-11 (wt*Dis*Cas7-11), or a high-performing triple mutant of Cas7-11 (*Dis*Cas7-11^Y312K/D988K/D1580R^). *Gaussia* luciferaserelative expression is quantified as *Gaussia* luciferase expression normalized to *Cypridina* luciferase expression. Fold changes are relative to rates with d*Dis*Cas7-11. **(f)** Schematic showing the exons and introns targeted for endogenous PRECISE editing for each endogenous transcript evaluated. Only flanking exons are shown with position indicated by number. Flanking exon and targeted intron sizes and cargo and guide positions are to scale. Transcripts per million expression levels for the targeted genes in HEK293FT cells are plotted. **(g)** PRECISE RNA editing quantified by next-generation sequencing for a panel of endogenous targets at the positions indicated in (f) with different *Dis*Cas7-11 mutants. Percent editing represents the ratio between junction reads containing barcoded silent mutations to total reads of the targeted exon-exon boundary.

**Figure 3.**
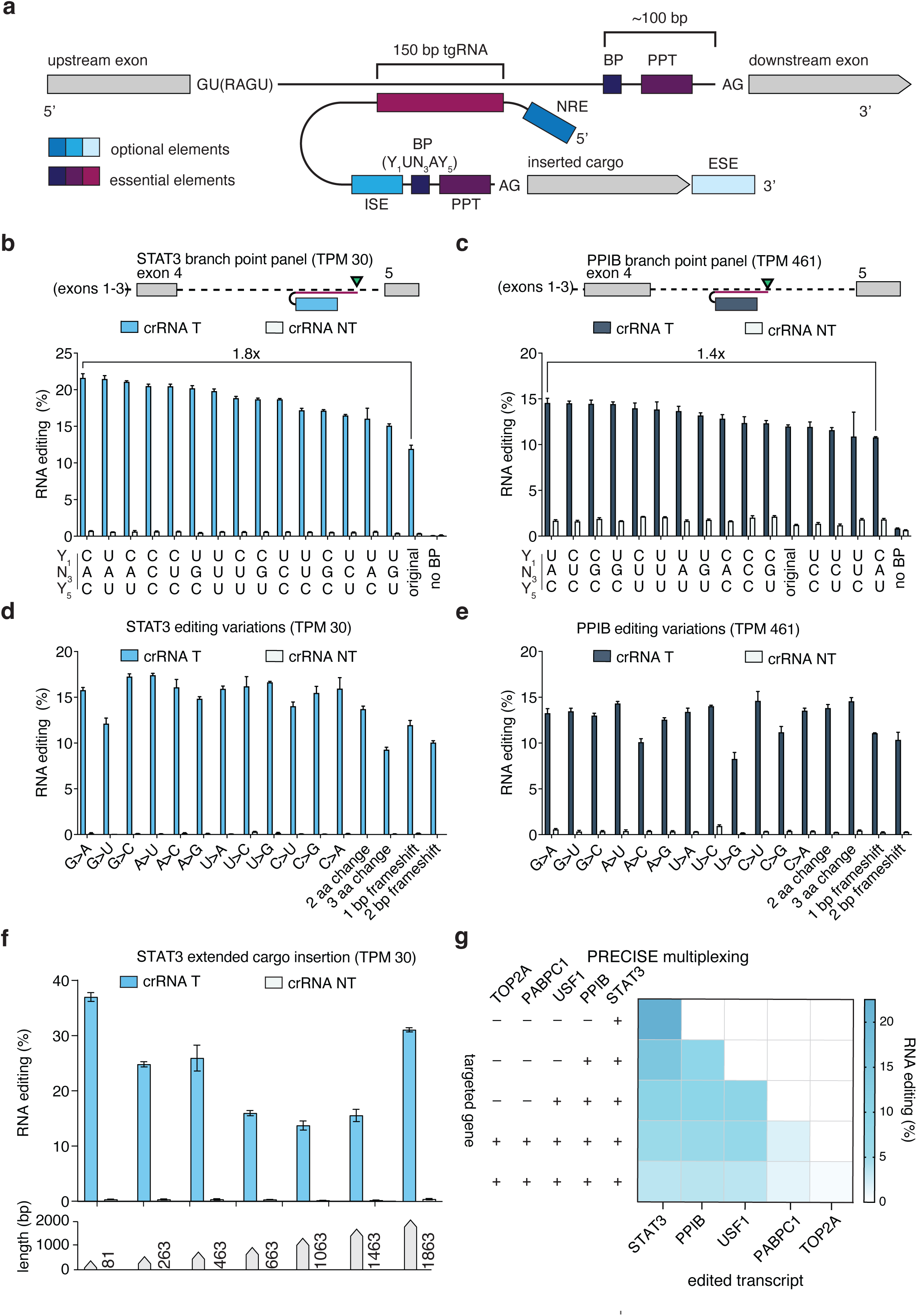
Optimization and evaluation of PRECISE editing capabilities, including small and large edits, insertions, and multiplexing. **(a)** Schematic showing the 3′ trans-splicing approach for optimization of PRECISE template branch points, intronic splicing enhancers, exonic splicing enhancers, and nuclear retention elements. **(b)** Evaluation of a panel of different branch point sequences for PRECISE 3′ trans-splicing editing of the endogenous STAT3 transcript. **(c)** Evaluation of a panel of different branch point sequences for PRECISE 3′ trans-splicing editing of the endogenous PPIB transcript. **(d)** Installation of all possible single point mutations and other changes at the STAT3 endogenous transcript exon 6 with PRECISE editing. Editing is compared with targeting and non-targeting *Dis*Cas7-11 guides. **(e)** Installation of all possible single point mutations and other changes at the PPIB endogenous transcript exon 5 with PRECISE editing. Editing is compared with targeting and non-targeting *Dis*Cas7-11 guides. **(f)** Installation of varying sized inserts at the STAT3 endogenous transcript exon 6 with PRECISE editing. Editing is compared with targeting and non-targeting *Dis*Cas7-11 guides. **(g)** Multiplexed PRECISE editing of 81–120 bp sized inserts at five different endogenous transcripts: *STAT3* (S), *PPIB* (P), *USF1* (U), *PABPC1* (P), and *TOP2A* (T). Editing is measured via next-generation sequencing.

Lastly, we were curious if the specific cleavage activity of *Dis*Cas7-11 was a strict requirement for the high-efficiency trans-splicing we observed. We compared PRECISE editing with *Dis*Cas7-11 versus *Lwa*Cas13a, *Psp*Cas13b, and *Rfx*Cas13d cleavage. We hypothesized that the collateral activity of Cas13 orthologs^37^ would non-specifically degrade transcripts, leading to inefficient trans-splicing. Upon RNA sequencing, we observed that all three Cas13 orthologs had substantially reduced trans-splicing activity, suggesting that precise cleavage in the pre-mRNA was necessary for efficient RNA editing (**Fig. 1h**). To mitigate this effect, we tested a high-fidelity version of *Rfx*Cas13d (hfCas13d) engineered for substantially reduced collateral activity ^38^. Using a panel of crRNAs, we found that low levels of boosted trans-splicing on PPIB could be observed by RNA sequencing, but they were ~10-fold lower than PRECISE editing with *Dis*Cas7-11 **(Extended Data Fig. 3c)**, suggesting that either collateral activity is still an issue or the precise cuts of *Dis*Cas7-11 are important for the observed boost in trans-splicing activity.

### Diverse Cas7-11 orthologs are compatible with PRECISE editing

To explore potential improvements for the PRECISE system, we turned to Cas7-11 diversity as a source for new trans-splicing enhancing nucleases. We selected a panel of diverse Cas7-11 orthologs from different bacterial genomes and metagenomic assemblies, and evaluated them for RNA knockdown. As orthologs may be cross-compatible with different direct repeats (DR) and crRNA scaffolds, leading to improved activity ^39^, we tested orthologs for Gluc knockdown with not only their own DR, but predicted DRs from all other orthologs, both in the sense and antisense direction. We found significant cross-compatibility between orthologs and DRs and multiple efficient nucleases, including Cas7-11 orthologs from Hydrothermal sediment (HsmCas7-11) and Hydrothermal vent sediment (HvsCas7-11) metagenome sequences, which had improved performance with the *Canditatus Scalindua brodae (Csb)* Cas7-11 DR (**Extended Data Fig. 4a)**. Applying these combinations for PRECISE, we found that, while they effectively boosted trans-splicing efficiency relative to a non-targeting control, they were not superior to *Dis*Cas7-11 (**Extended Data Fig. 4b-c**).

### Engineering Cas7-11 with rational mutagenesis improves RNA cleavage efficiency

In an effort to further enhance PRECISE editing rates, we generated a panel of rational variants of the wild-type *Dis*Cas7-11, building off our previously solved structure^33^. Focusing on residues in proximity to the crRNA and target RNA, we replaced polar or negatively charged residues with positively charged amino acids, either lysine (K) and arginine (R) (**Fig. 2a-b**). We first evaluated the cleavage efficiency of the Cas7-11 mutants on a luciferase reporter assay, and found that a variety of substitutions improved transcript knockdown (**Fig. 2c** and **Extended Data Fig. 5**). In particular, a subset of mutations (Y312K, D988K, D1580R, and D981K) each offered a moderate enhancement with up to 33% higher knockdown activity over the wild-type Cas7-11 (**Fig. 2c**), probably due to enhanced binding of the Cas7-11 mutants to their crRNAs or targeted transcripts (**Fig. 2b**). Effective single mutations of *Dis*Cas7-11 with improved cleavage activity had additive effects as double and triple mutation *Dis*Cas7-11 variants, ultimately resulting in the selection of an engineered triple Y312K/D988K/D1580R mutant, termed enhanced *Dis*Cas7-11 or “e*Dis*Cas7-11.” e*Dis*Cas7-11 had significantly more knockdown activity, peaking at 93% activity (**Fig. 2c**).

### Enhanced *Dis*Cas7-11 improves PRECISE efficiency

We next tested e*Dis*Cas7-11 in the luciferase splicing assay described earlier to evaluate the impact of higher catalytic activity on overall trans-splicing rates **(Fig. 2d)**. The replacement of wild-type *Dis*Cas7-11 with e*Dis*Cas7-11 increased reporter splicing rates by ~1.4-2.0 fold (**Fig. 2e**), indicating that 3′ transcript cleavage efficiency has a major impact on trans-splicing.

Using the new e*Dis*Cas7-11 construct, we next applied PRECISE editing to a wide range of endogenous transcripts. Applying the same crRNA and tgRNA screening strategy used to edit STAT3 and PPIB, we identified designs that allow for efficient endogenous PRECISE editing for the RPL41, PABPC1, TOP2A, STAT3, and SHANK3 pre-mRNAs, which represent a large range of expression levels ( >8,000 TPM, RPL41) to very low (< 15 TPM, SHANK3) (**Fig. 2f**). For all 7 endogenous transcripts tested for PRECISE editing, e*Dis*Cas7-11 enabled improved PRECISE editing over using wild-type *Dis*Cas7-11 (**Fig. 2g**).

### Optimized branch points and splicing factor recruitment improve PRECISE efficiency

The mechanism of PRECISE means that features of the cargo RNA could have substantial impacts on efficiency, and we reasoned that modifications to the PRECISE cargos could further increase trans-splicing rates. Previous work on endogenous trans-splicing has shown the importance of specific sequence elements for incorporation of the cargo, such as the presence of polypyrimidine tracts (PPTs) and branch points (BPs) in the cargo (**Fig. 3a**), which have a significant impact on splicing rates^40^.

Our original cargo designs used a linker sequence from previous natural trans-splicing experiments ^40^ with a GAGGAA branch point sequence. These cargos inserted with moderate efficiency in our initial tests, potentially due to their branch point differing from the yUnAy^41^ human consensus motif. We hypothesized that alternate branch points matching the human consensus motif would promote recognition of our cargo for trans-splicing in human cells. Endogenous PRECISE cargos for STAT3 and PPIB were generated to screen each of the 16 potential motif-matching branch points (C/U-U-A/U/C/G-A-C/U) and these cargos were tested for trans-splicing. For STAT3, nearly all new branch points tested had higher splicing efficiency than base design, with the cUnAy branch point sequences performing best (**Fig. 3b**). For PPIB, more than half of the new consensus-motif branchpoint cargos were more efficient than the original cargo design (**Fig. 3c**). We also tested constructs containing a number of other highly conserved branchpoints, including the yeast branchpoint, and versions of the human or yeast consensus branch points with additional intervening linker sequences (**Extended Data Fig. 6a-c**), but these new sequences did not improve PRECISE editing over best performing human motif sequences. We ultimately selected the cUuAc branch point for our enhanced cargo design, due to its consistently high trans-splicing rates on both the endogenous STAT3 and PPIB transcripts. To further optimize trans-template cargos, we tested a series of intronic splicing enhancer (ISE) sequences, exonic splicing enhancer (ESE) sequences, and nuclear retention elements (NRE) from a variety of different contexts, finding that these regulatory sequences could also boost PRECISE editing (**Extended Data Fig. 6d-h, Supp. Table 2**). These features represent a variety of different orthogonal approaches for increasing efficiency at a specific target. However, rate increases for one gene (*e.g.*, STAT3) may not be universally optimal for a new target due to differences in exon size, intron size, positioning of splicing features (*e.g.*, natural branch points or enhancer sequences) and therefore some amount of screening may be useful to fully optimize a particular target.

### Improving trans-splicing rates with splicing factor fusions to Cas7-11

Transcript splicing inside cells is normally controlled by the spliceosome, a large complex consisting of over 100 proteins and nucleoproteins^42^. The spliceosome is responsible for recognizing and binding sequence features on an mRNA transcript (such as the branch point and poly-pyrimidine tract) and depends on the recruitment of a variety of RNA binding proteins and subunits ^43,44^. We hypothesized that by fusing some early spliceosome members to Cas7-11, we could induce the formation of the spliceosome at our transcript of interest, potentially enhancing the overall efficiency of PRECISE editing. We explored both N- and C-terminal fusions of several splicing factors to *Dis*Cas7-11, including RBM17, SF3B6, and both U2AF1 and U2AF2, and found that across four different endogenous transcripts, STAT3, PPIB, TOP2A, and PABPC1, we could enhance trans-splicing efficiency, especially with C-terminal fusions **(Extended Data Fig. 7a-d, Supp. Table 3)**. Overall, we observed between 1.6–4.1-fold improvement in splicing due to splicing factor recruitment with U2AF1 generally providing the largest boost in efficiency across the 4 transcripts tested (**Extended Data Fig. 7e**). An added benefit of splicing factor recruitment with PRECISE (PRECISE-sf) is that most of the tested splicing factor proteins are small in size, ranging from 51 to 240 amino acids, making linked constructs theoretically still small enough to be encapsulated in smaller delivery vectors such as AAVs.

### PRECISE editing can install single base, kilobase-scale, and multiplexed edits

In order to show that PRECISE editing is capable of inserting diverse sequences, we generated a set of cargos installing each possible single-base transition and transversion for two endogenous targets, STAT3 and PPIB (**Supp. Table 2)**. Each of these cargos inserted with similar rates (**Fig. 3d-e**), showing that editing was not affected by cargo sequence. We further generated cargos that install multiple-nucleotide changes in the inserted exon to demonstrate that PRECISE editing can be used to make larger changes, such as those resulting in single or multiple amino-acid edits (**Extended Data Fig. 7f-h**). We also generated a set of constructs with 1, 2, 3, 12, 24, or 48 bp insertions, and another set with a range of deletions, to demonstrate that both frameshift changes and larger modifications to the spliced sequence are possible (**Extended Data Fig. 7f-h**). To show that PRECISE could flexibly insert cargos of arbitrary length, we next generated another set of constructs to measure insertion of larger sequences. Starting with the STAT3 cargo used previously, we generated progressively larger inserts ranging from 81 to 1,863 nt with high efficiency (**Fig. 3f**). Encouragingly, there was only a moderate decrease in efficiency, from ~37% to ~34%, when comparing efficiency of insertion for the shortest and longest sequences, suggesting that even longer insertions may be possible without a major reduction in efficiency. Lastly, we hypothesized that because PRECISE editing was efficient on single transcripts and does not generate double stranded breaks (DSB) on DNA, multiplexed editing should be efficient for multiple transcripts with minimal effects to the cell. To evaluate this, we tested multiplexing on the 5 efficient targets, STAT3, PPIB, USF1, PABPC1, and TOP2A, and found that multiplexing of 2–5 transcripts simultaneously was possible with high efficiency (**Fig. 3g**).

### PRECISE editing generates detectable protein products

We next determined how PRECISE-mediated RNA editing translated to protein-level changes. To analyze protein edits resulting from PRECISE editing, we overexpressed a USF1 transcript and used PRECISE editing to introduce a C-terminal flag tag to the USF1 protein, such that the size of the protein increases relative to its wild-type size, allowing visualization by western blotting. We recorded ~85% trans-splicing with *Dis*Cas7-11 targeting (PRECISE) and 16% with cargo-only trans-splicing (**Fig. 4a**). By Western blotting, we observed corresponding protein editing as evidenced by the expected trans-spliced protein size shift. Semi-quantitative analysis revealed that the corresponding protein editing was ~45% and 19% for PRECISE editing and natural trans-splicing, respectively (**Fig. 4a**). We next validatee protein level edits on two endogenous transcripts previously edited with PRECISE, HDAC1 and PPIB. We found that RNA editing mediated via PRECISE at ~15% for HDAC1 and 18% for PPIB translated to the protein level by Western blot analysis (**Fig. 4b-c**). Correspondingly, semi-quantitative analysis revealed that the protein editing matched the overall level of RNA editing (**Fig. 4b-c**).

**Figure 4.**
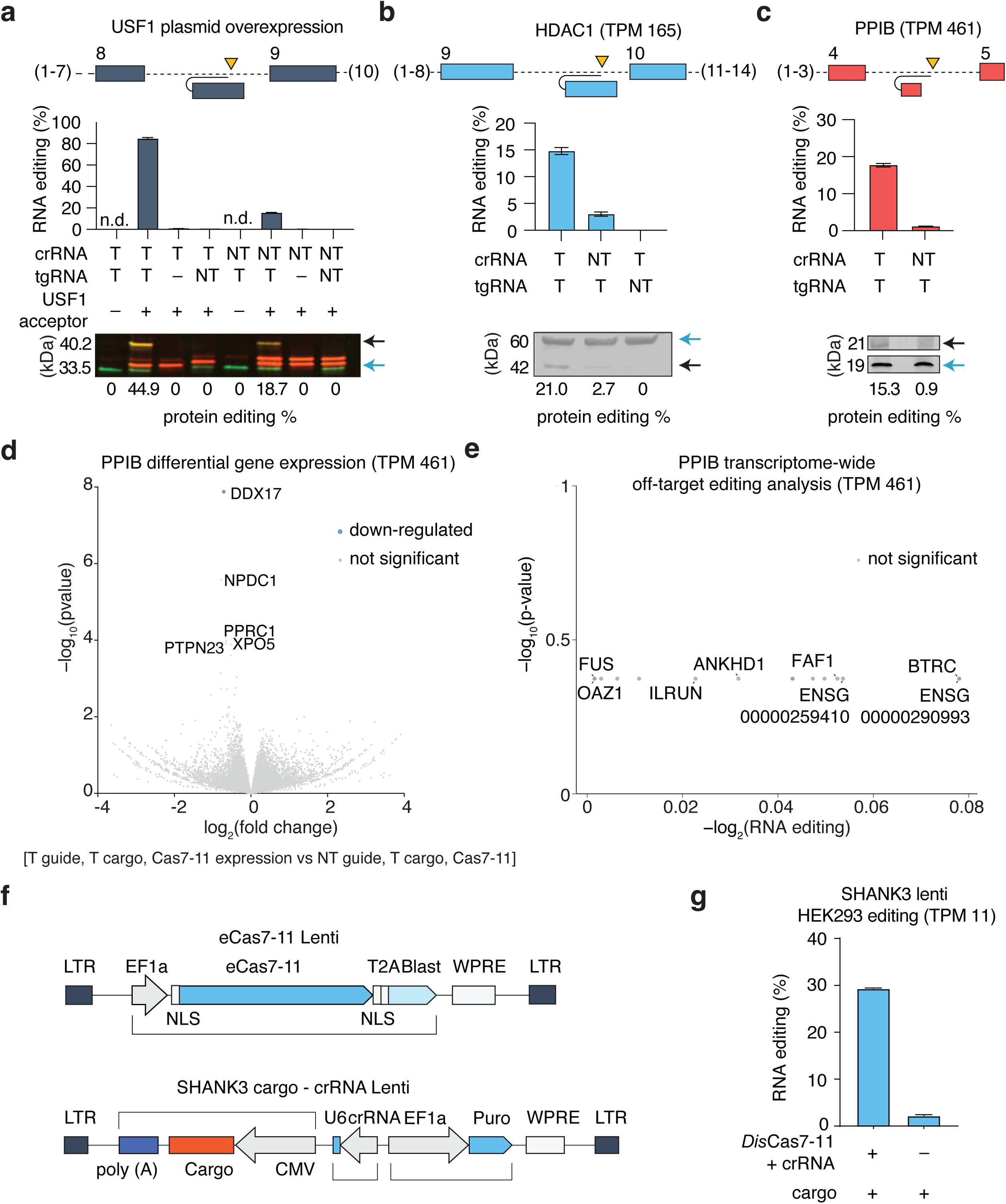
Evaluation of protein level edits, transcriptome-wide specificity, and delivery by lentivirus. **(a)** Evaluation of USF1 protein editing due to PRECISE-based RNA writing by Western Blotting. USF1 is overexpressed and is targeted with either targeting or non-targeting *Dis*Cas7-11 guides and targeting or non-targeting cargo hybridization guides. Shown is the RNA editing rate (top) and Western blot gel with semi-quantitative protein editing rate (bottom). Arrows indicate either the unedited protein (blue) or the edited trans-spliced protein (black). (**b)** Evaluation of endogenous HDAC1 protein editing due to PRECISE-based RNA editing by Western Blotting. The HDAC1 transcript is targeted with either targeting or non-targeting *Dis*Cas7-11 guides and targeting or non-targeting cargo hybridization guides. Shown is the RNA editing rate (top) and Western blot gel with semi-quantitative protein editing rate (bottom). Arrows indicate either the unedited protein (blue) or the edited trans-spliced protein (black). **(c)** Evaluation of endogenous PPIB protein editing due to PRECISE-based RNA writing by Western Blotting. The PPIB transcript is targeted with either targeting or non-targeting *Dis*Cas7-11 guides and a targeting cargo hybridization guide. Shown is the RNA editing rate (top) and Western blot gel with semi-quantitative protein editing rate (bottom). Arrows indicate either the unedited protein (blue) or the edited trans-spliced protein (black). **(d)** Volcano plots of differentially expressed genes as measured by transcriptome-wide sequencing of cells with PPIB PRECISE editing. **(e)** Number of significant trans-spliced off-targets identified by transcriptome-wide sequencing of cells following PRECISE editing of PPIB. **(f)** Schematic of PRECISE RNA writing for 3′ trans-splicing via lentiviral delivery. Shown are the construct designs for a two-vector system. **(g)** Demonstration of PRECISE editing of the SHANK3 transcript with lentiviral delivery of *Dis*Cas7-11 expression, guides, and RNA trans-template in HEK293FT cells. RNA writing of a 108 bp sequence into the endogenous SHANK3 transcript with targeting and non-targeting *Dis*Cas7-11 guides is compared.

### PRECISE editing minimally perturbs both target transcript levels and the general transcriptome

To explore the effect of PRECISE editing on the cell at the transcriptomic level, both in terms of target transcript levels and general off-targets, we quantified the expression levels of transcripts targeted by PRECISE editing to understand if cleavage or trans-splicing results in altered expression levels. Using quantitative-PCR assays that measure the total expression of unedited and edited transcripts, we observed no significant changes in expression levels, suggesting that PRECISE editing does not contribute towards knockdown or increased expression of targeted transcripts **(Extended Data Fig. 8a)**. We also analyzed RNA-sequencing for differentially regulated gene expression in response to the PRECISE editing and found only a single gene, DDX17, to be differentially expressed between the targeting and non-targeting crRNA PRECISE editing conditions, which correspond to PRECISE editing rates of ~15 and ~1% respectively (**Fig. 4d and Extended Data Fig. 8b-c**). DDX17 is an RNA helicase, which has been implicated in alternative RNA splicing and ribosome biogenesis^45^. We also analyzed the whole-transcriptome mRNA sequencing for off-target trans-splicing and found detection of our PPIB trans-spliced transcripts, but failed to detect any off-target spliced transcripts that were significantly occurring **(Fig. 4e and Extended Data Fig. 8d)**. In addition, comparing a targeting crRNA, targeting tgRNA, and active *Dis*Cas7-11 condition to a non-targeting crRNA, targeting tgRNA, and dead *Dis*Cas7-11 condition, we found 2 significantly down-regulated genes and 5 significantly up-regulated genes (**Extended Data Fig. 8b**). This differential expression likely indicates that the presence of cleaved transcripts as expected in the active, targeting Cas7-11 could cause moderate perturbations in some cell types.

### PRECISE editing with lentiviral delivery enables correction of pathogenic mutations

We extended the use of 3′ PRECISE trans-splicing to primary cell types, focusing on neurons as an initial demonstration. To deliver to neurons, we designed a lentiviral delivery system with *Dis*Cas7-11 in one vector and the crRNA and trans-template cargo in a separate vector (**Fig. 4f**) to avoid cleavage without concurrent splicing. We packaged the SHANK3 targeting PRECISE constructs in lentivirus and found that up to 30% RNA editing could be achieved in HEK293FT cells, validating that PRECISE editing is efficient with viral delivery (**Fig. 4g**).

### PRECISE editing can replace 5′ exons on mRNA

While 3′ PRECISE editing can install a majority of desirable edit types in mRNAs, situations where the first exon or portions of only the 5′ ends of genes need to be edited demand alternative approaches. To edit the upstream sequence of a mRNA instead, PRECISE can be adapted for 5′ trans-splicing cargos: 5′ trans-splicing constructs have the opposite arrangement of cargo elements and *Dis*Cas7-11 crRNAs targeting upstream of the cargo tgRNA rather than downstream. We designed a 5′ trans-splicing cargo with 5′ replacement exons expressed with a trailing tgRNA on the 3′ end and an intervening linker that includes splicing sequence motifs normally denoting a 5′ exon boundary (**Fig. 5a**). Similar to the 3′ end of the intron, which normally contains the BP, PPT, and SS sequences, the 5′ intron end contains a splice donor contained in the human consensus splicing sequence (CSS), with splicing further facilitated by intronic splicing enhancers (ISEs) (**Fig. 5a, Supp. Table 2**). Using these designs, we piloted 5′ trans-splicing on the *HTT* exon 1, where triplet expansions cause Huntington′s Disease pathology, replacing the endogenous exon 1 with a new copy of the exon carrying no CAG repeats and being only 68 nt in length. In our initial experiment, we found that low levels of trans-splicing could be observed in the ~2–3% editing range **(Fig. 5b)**. Because of this low editing, we hypothesized editing could be boosted by removing the poly(A) signal on the cargo via cleavage of the cargo at the 3′ end with *Dis*Cas7-11, promoting cargo retention in the nucleus, by cleaving. This approach improved 5′ PRECISE substantially, producing ~35–38% trans-splicing efficiencies across multiple tgRNAs tested **(Fig. 5b)**. We next sought to optimize our cargo design, evaluating a panel of ISEs, BPs, and CSS sequences, and evaluated the efficiency of these different cargoes using NGS. We found that the addition of ISE, BP, and optimized CSS sequences resulted in higher efficiencies **(Fig. 5c)**. This trans-splicing was *Dis*Cas7-11 dependent, as a non-targeting crRNA eliminated the observed trans-splicing (**Fig. 5c**).

**Figure 5.**
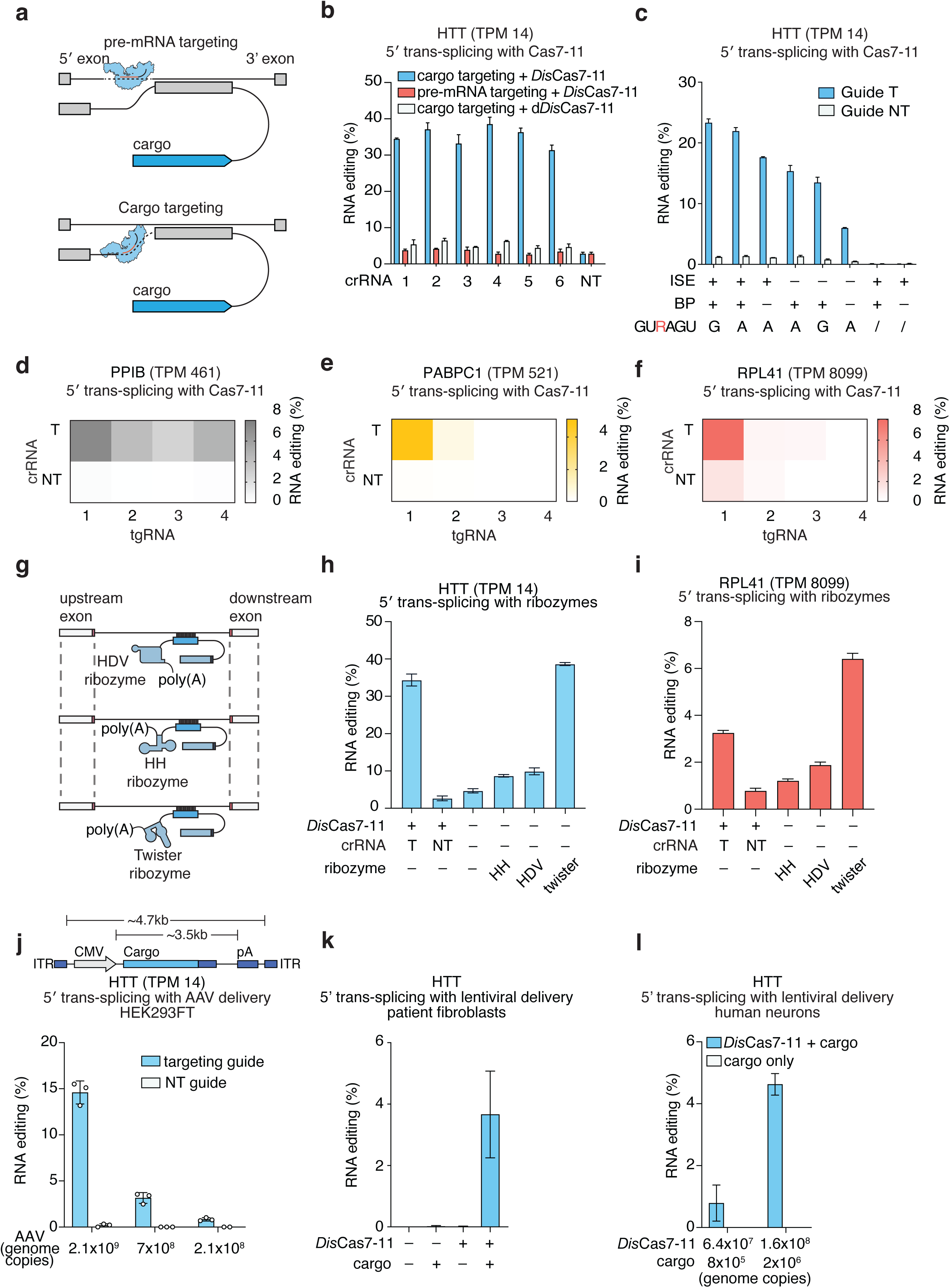
Development of PRECISE editing for efficient 5′ trans-splicing without proteins and only using the RNA trans-splicing template. **(a)** Schematic showing 5′ PRECISE with two different approaches: 1) Pre-mRNA targeting is shown with a trans-splicing cargo and *Dis*Cas7-11 crRNA targeted to a 3′ region of the intron area targeted by the cargo hybridization guide; and 2) Cargo targeting is shown where the *Dis*Cas7-11 guide cleaves the 3′ end of the trans-template, separating the poly(A) tail from the cargo. **(b)** RNA editing rates for using PRECISE editing to replace HTT exon 1 with a short synthetic exon using *Dis*Cas7-11 constructs that either target the pre-mRNA intron or the cargo template. Active *Dis*Cas7-11 is compared with d*Dis*Cas7-11. **(c)** RNA editing rates using PRECISE to replace HTT exon 1 with a short synthetic first exon. Rates are compared for constructs with different combinations of regulatory sequences, including intronic splicing enhancers (ISEs), branch points (BPs), and splicing signals. Cargos with either no 5′ consensus splicing signal, indicated as (/), or a GUGAGU or GUAAGU sequence are compared. **(d)** RNA editing rates for 5′ trans-splicing PRECISE editing on the endogenous PPIB transcript using a set of cargos with hybridization guides targeting different regions of PPIB intron 4. A *Dis*Cas7-11 crRNA targeting the 3′ end of the cargo payload is compared with a non-targeting *Dis*Cas7-11 guide. **(e)** RNA editing rates for 5′ trans-splicing PRECISE editing on the endogenous PABPC1 transcript using a set of cargos with hybridization guides targeting different regions of PABPC1 intron 13. A targeting *Dis*Cas7-11 crRNA targeting the 3′ end of the cargo payload is compared with a non-targeting *Dis*Cas7-11 guide **(f)** RNA editing rates for 5′ trans-splicing PRECISE editing on the endogenous RPL41 transcript using a set of cargos with hybridization guides targeting different regions of RPL41 intron 2. A targeting *Dis*Cas7-11 crRNA targeting the 3′ end of the cargo payload is compared with a non-targeting *Dis*Cas7-11 guide. **(g)** Schematic of protein-free PRECISE editing using ribozymes to liberate trans-templates of the poly(A) tail and enable trans-splicing based RNA writing. **(h)** RNA editing rate of 5′ trans-splicing PRECISE editing on the endogenous HTT transcript with trans-templates cut by either *Dis*Cas7-11 or one of several ribozymes. *Dis*Cas7-11 targeting is compared to a non-targeting guide. **(i)** RNA editing rate of 5′ trans-splicing PRECISE editing on the endogenous RPL41 transcript with trans-templates cut by either *Dis*Cas7-11 or one of several ribozymes. *Dis*Cas7-11 targeting is compared to a non-targeting guide. **(j)** RNA editing rate of 5’ PRECISE editing on the endogenous HTT transcript in HEK-293FT cells by AAV delivery. Both targeting cargo guides and non-targeting cargo guides are compared using AAV delivery. **(k)** RNA editing rate of 5′ trans-splicing PRECISE editing on the endogenous HTT transcript in expanded-repeat patient-derived fibroblasts via lentiviral delivery. Trans-templates are cut by *Dis*Cas7-11 using a two-vector system at different titer amounts. **(l)** RNA editing rate of 5′ trans-splicing PRECISE editing on the endogenous HTT transcript in iPSC-derived human neurons via lentiviral delivery. Trans-templates are cut by *Dis*Cas7-11 using a two-vector system at different titer amounts.

After demonstrating the principle of *Dis*Cas7-11-facilitated 5′ PRECISE editing on *HTT*, we next tested a panel of more endogenous targets for 5′ exon replacement. Using the optimal cargo design on the *PPIB*, *PABPC1*, and *RPL41* pre-mRNA transcripts, we found that *Dis*Cas7-11 targeting of the cargo could enhance 5′ trans-splicing at all three genes, producing a range of 5– 8% RNA editing (**Fig. 5d-f**). As 5′ trans-splicing PRECISE efficiency was cargo dependent, with most efficient cargos for each endogenous transcript having targeting cargo guides closer to the 3′ splicing acceptor. The 5′ trans-splicing efficiencies support the notion that PRECISE editing is capable of both 5′ or 3′ trans-splicing editing.

Lastly, using a reporter system, we sought to show whether dual 5′ and 3′ trans-splicing could be achieved to replace internal exons. We designed a luciferase reporter split across three exons with the middle exon containing a set of pre-termination stop codons **(Extended Data Fig. 9)**. We then designed a template cargo with two guide regions targeting the first and second intron and flanking the middle luciferase exon. In addition, a panel of *Dis*Cas7-11 crRNAs were designed as pairs in intron 1 and intron 2 to liberate the existing stop codon exon from the pre-mRNA. We evaluated all possible pairs of crRNAs and tgRNA designs, finding that certain combinations enabled up to ~80% replacement of the second luciferase exon and that this was dependent on an active *Dis*Cas7-11 being used for RNA cleavage **(Extended Data Fig. 9)**. Moreover, non-targeting crRNAs or tgRNAs had substantially reduced activity, verifying that PRECISE editing of internal exons depends on guide RNA.

### PRECISE editing with ribozymes improve 5′ trans-splicing efficiency

As *Dis*Cas7-11 induced removal of the poly(A) tail on the trans-template cargo was so effective at promoting trans-spliced editing, we hypothesized whether simpler mechanisms of poly(A) removal would also succeed. Ribozymes are catalytically active RNA molecules capable of specific ribonucleolytic self-cleavage, which we thought could substitute for *Dis*Cas7-11 trans-template cleavage. To evaluate this hypothesis, we designed a set of trans-templates using a panel of ribozymes, including hepatitis delta virus (HDV), hammerhead (HH), or twister ribozymes, to remove the 3′ poly(A) tail (**Fig. 5g**). We found that overall, the twister and HDV ribozymes performed the best across RPL41 and HTT and that 5′ trans-splicing was improved ~10-fold when using the twister ribozyme, indicating that separation of poly(A) tail is indeed critical for efficient 5′ trans-splicing (**Fig. 5h-i**). Moreover, the twister ribozyme based trans-templates are more active than the *Dis*Cas7-11 cargo targeting designs and have the added benefit of being a single-component system, requiring only the trans-template.

### AAV and lentiviral delivery of PRECISE editing constructs in cell lines and neurons

We next explored whether the single-component simplicity of ribozyme-based 5′ trans-splicing was suitable for efficient HTT exon 1 PRECISE editing via AAV or lentiviral delivery. We first sought to evaluate AAV delivery with the HTT trans-splicing twister ribozyme design and packaged AAV8 vectors with this payload (**Fig. 5j**). We transduced HEK293 cells with these AAVs and found that ~15% RNA editing could be achieved in a dose-dependent fashion, validating that this editing approach was compatible with AAV delivery (**Fig. 5j**). In human iPSC-derived neurons, we found that the PRECISE editing AAVs could achieve ~0.5% RNA editing efficiency **(Extended Data Fig. 10a)**. Lastly, we showed that truncated small *Dis*Cas7-11 variants previously developed^33^ could also achieve high levels of PRECISE editing in HEK293FT cells, especially when engineered with mutations that increase catalytic efficiency **(Extended Data Fig. 10b-c)**. Smaller versions of Cas7-11 PRECISE editors are useful for further increasing the packaging capacity of the AAV vectors.

We next designed and packaged a lentivirus system to evaluate HTT transcript correction in a HD patient derived fibroblast line containing 66x CAG repeats. In HEK293FT cells, the lentiviral system could achieve ~18% RNA editing levels on the HTT transcripts **(Extended Data Fig. 10d)**. We transduced this fibroblast line with the lentivirus and observed PRECISE editing that peaked at ~4% correction of the HTT transcript (**Fig. 5k**). We next tested induced human cortical neurons with the lentiviral delivered PRECISE system, and similarly found that ~4.5% editing could be achieved, further validating the generalizability of PRECISE editing for different cell types, including non-dividing primary cells (**Fig. 5l**).

## Discussion

Programmable replacement of any length of sequence in mRNAs is an especially powerful feature of PRECISE editing, allows for versatile correction of small or large edits, insertions, or deletions. We showcase PRECISE editing on nine different endogenous transcripts with efficiencies ranging from 5% to 50% and up to 85% on reporter transcripts, demonstrate large-scale exon replacement up to 1,863 bp, show varying sized deletions, and illustrate several therapeutically relevant applications, including SHANK3 exon replacement in neurons (autism spectrum disorder) and HTT exon 1 replacement via single-vector AAV delivery (Huntington’s disease). Because exon replacement with PRECISE editing allows targeting of any sized edit, insertion, or deletion, 140,267 pathogenic variants in ClinVar are addressable, allowing targeting of ~96% of known disease-causing mutations, which represents variants that arise within multi-exon transcripts.

PRECISE editing offers the ability for both efficient 5′ and 3′ trans-splicing. For 3′ trans-splicing, we found that *Dis*Cas7-11 targeting of intron enabled efficient cleavage by releasing cis-exons and facilitating the trans-template to dominate the splicing reaction. For 5′ trans-splicing, we found that simply liberating the poly(A) tail from the trans-template via ribozymes allowed for efficient trans-splicing, presumably through nuclear retention of the trans-template and efficiency of the trans-template alone to serve as the 5′ donor without pre-mRNA cleavage. An advantage of PRECISE editing is that there are numerous parameters that can be optimized, including *Dis*Cas7-11 guide choice, template hybridization sequence length, linker length, splicing signals, and branching points, to enable maximal possible editing efficiency, which allows for more optimization than other DNA and RNA editing approaches. A major advancement with PRECISE editing for 5′ trans-splicing is that it is protein-free, involving only a single RNA component, and so is simple enough to allow up to ~3.7 kb trans-template payload to be delivered in a single AAV, allowing for facile translation to diverse tissues using the plethora of tissue specific AAV capsids that have now been developed ^46,47^.

Further work is needed to characterize and optimize PRECISE editing in diverse cells and organisms. As splicing machinery is universally expressed in all cell types, it is expected that PRECISE editing will be broadly applicable, although more will be needed to understand off-target trans-splicing, strategies to mitigate non-specific trans-splicing, and the effects off-targets might have on the cell. As PRECISE edits at the RNA level, more work will be needed to understand the kinetics of editing, whether transient editing via mRNA delivery is possible for temporary editing, and how permanent AAV-based editing is for correction of genetic disease. There are more than 7,000 genetic diseases with hundreds of thousands of genetic variants and PRECISE editing via large exon replacement allows for the treatment of many variants for a given gene at once safely and reversibly, promoting the potential development of gene therapies that address many patient variants with a single treatment.

## Acknowledgments

We would like to thank C. Fell for his assistance with the Galaxy RNA sequencing analysis workflow; K. Jiang for support during the initial RNA sequencing sample preparation; A. Bubnys for generously providing us with neurons; and M. Wienisch for supplying the HTT fibroblasts, which were originally sourced from Coriell. We thank the members of the Abudayyeh-Gootenberg labs for support and advice.

## Author contributions

O.O.A. and J.S.G conceived the study and participated in the design, execution, and analysis of experiments. C.S. and A.K. designed and performed all the experiments in this study and analyzed the data. M.M.C. and S.P.N. designed and assisted with characterizing the technology. N.Z. developed and tested initial luciferase and endogenous transcript data. K.D. assisted with all experiments in the study. K.K. and H.N. determined residues for Cas7-11 mutagenesis and engineering to improve activity and helped with analysis. C.S., A.K., O.O.A., and J.S.G wrote the manuscript with help from all authors.

## Data and materials availability

Sequencing data will be available at Sequence Read Archive. Expression plasmids are available from Addgene under UBMTA; support information and computational tools are available at https://www.abugootlab.org/.

## Figure Legends

**Extended Data Figure 1.**
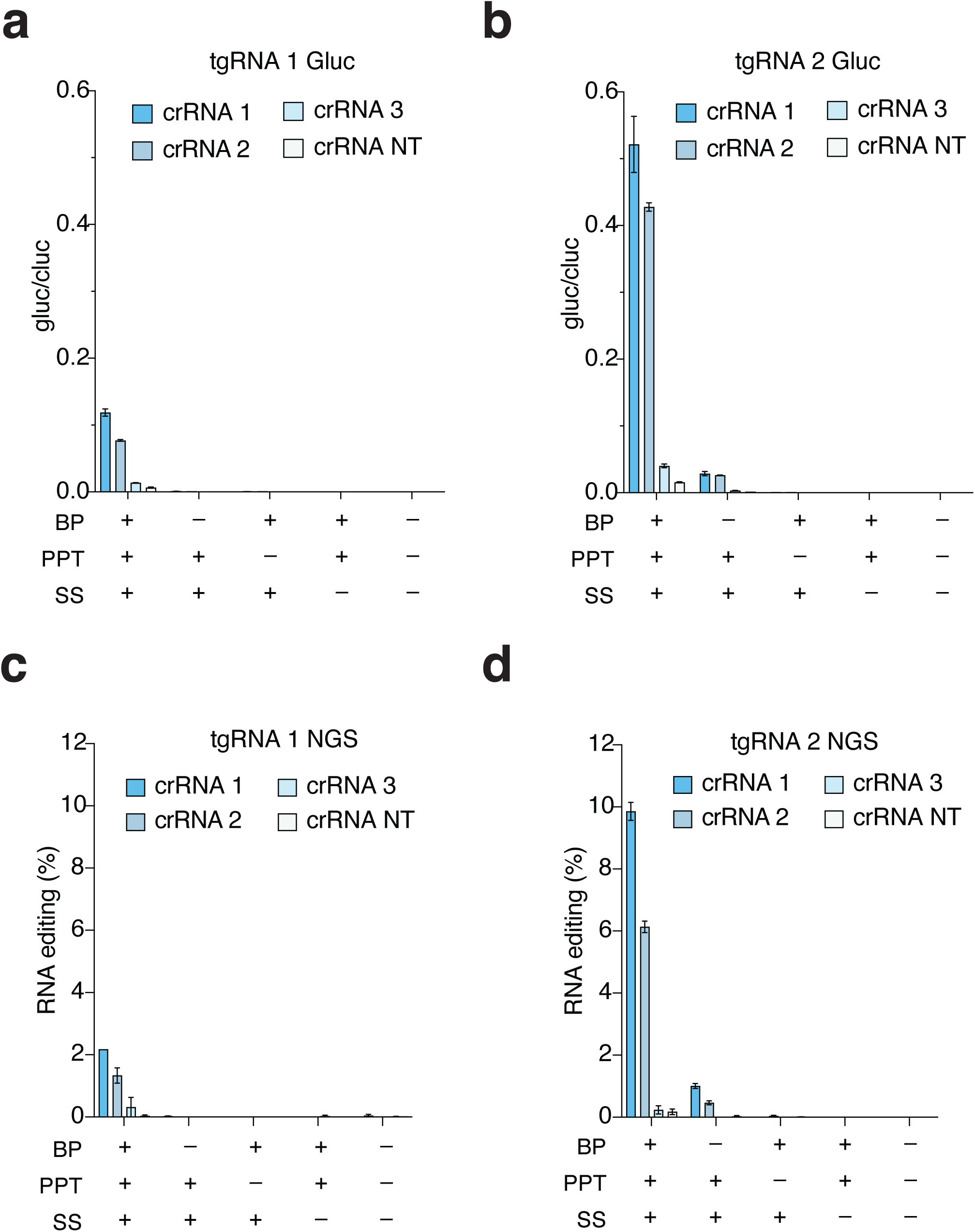
Optimization of PRECISE editing on Gluc reporter transcripts. **(a)** Splicing assay employing a split-luciferase reporter reconstituting a *Gaussia* luciferase transcript through PRECISE RNA editing, contrasting 3′ cargos with and without a branching point (BP), polypyrimidine tract (PPT), or splicing signal (SS). Trans-template contains C-terminal fragments of *Gaussia* luciferase and a hybridization region. Three *Dis*Cas7-11 targeting guides are compared to a non-targeting guide. *Gaussia* luciferase values normalized to constitutively co-expressed *Cypridina* luciferase control. **(b)** Splicing assay employing a split-luciferase reporter reconstituting a *Gaussia* luciferase transcript through PRECISE RNA editing, contrasting 3′ cargos with and without a branching point (BP), polypyrimidine tract (PPT), or splicing signal (SS). Trans-template contains C-terminal fragments of G-luciferase and cargo hybridization region 2. Three *Dis*Cas7-11 targeting guides are compared to a non-targeting guide. *Gaussia* luciferase values normalized to constitutively co-expressed *Cypridina* luciferase control. **(c)** RNA editing rates of the PRECISE editing conditions in part (a) exploring different *Dis*Cas7-11 targeting guides and effect of different regulatory signals on the trans-template. **(d)** RNA editing rates of the PRECISE editing conditions in part (b) exploring different *Dis*Cas7-11 targeting guides and effect of different regulatory signals on the trans-template.

**Extended Data Figure 2.**
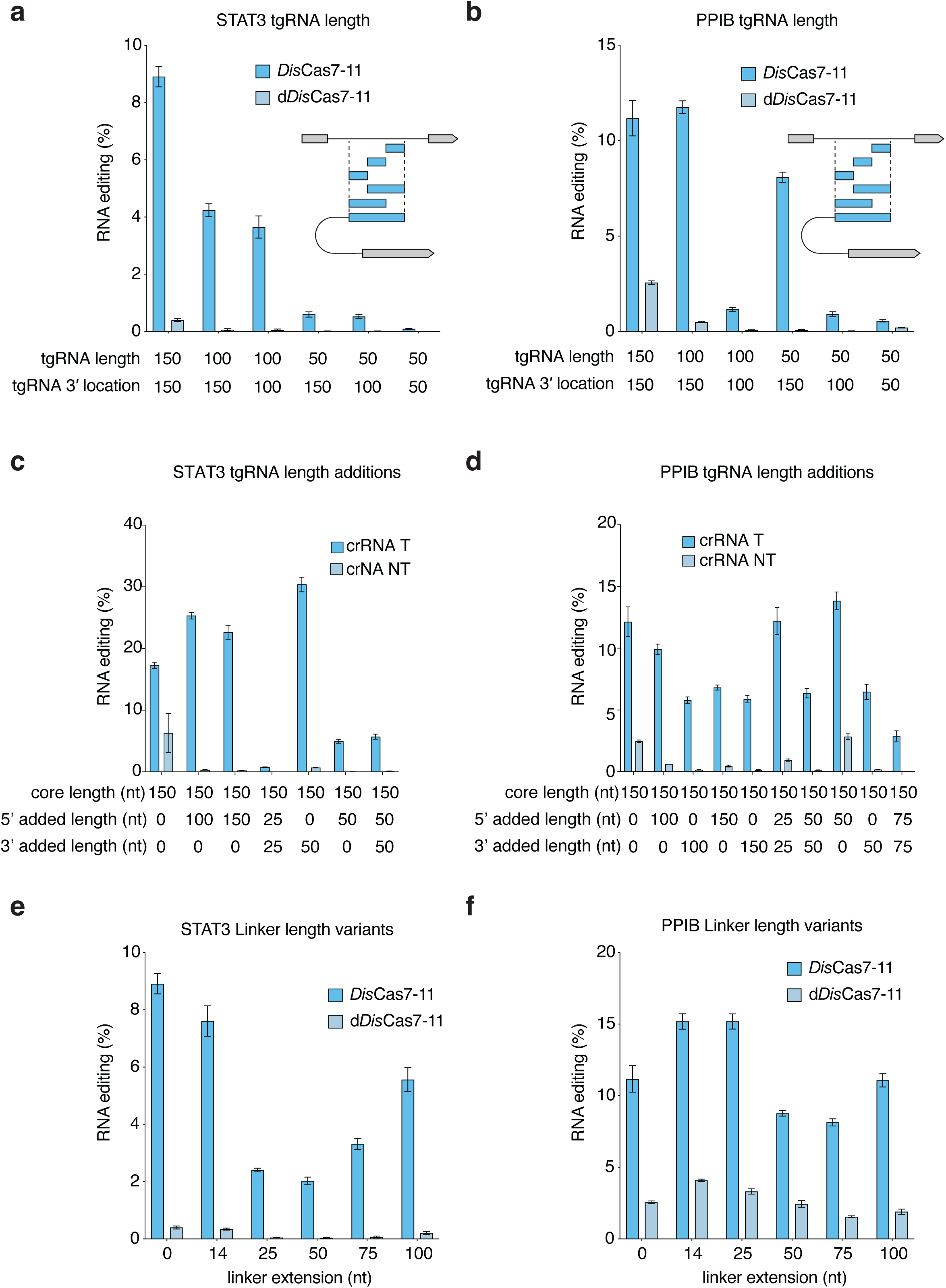
Optimization of PRECISE editing trans-templates on endogenous transcripts. **(a)** RNA editing rate of PRECISE on the endogenous STAT3 transcript using trans-templates with different cargo guide lengths and positions and either active *Dis*Cas7-11 or d*Dis*Cas7-11. **(b)** RNA editing rate of PRECISE on the endogenous PPIB transcript using trans-templates with different cargo guide lengths and positions and either active *Dis*Cas7-11 or d*Dis*Cas7-11. **(c)** RNA editing rate of PRECISE on the endogenous STAT3 transcript using trans-templates with different hybridization lengths and extensions. **(d)** RNA editing rate of PRECISE on the endogenous PPIB transcript using trans-templates with different hybridization lengths and extensions. **(e)** RNA editing rate of PRECISE on the endogenous STAT3 transcript using trans-templates with different linker lengths between the cargo guide and the splicing sequence. **(f)** RNA editing rate of PRECISE on the endogenous PPIB transcript using trans-templates with different linker lengths between the cargo guide and the splicing sequence.

**Extended Data Figure 3.**
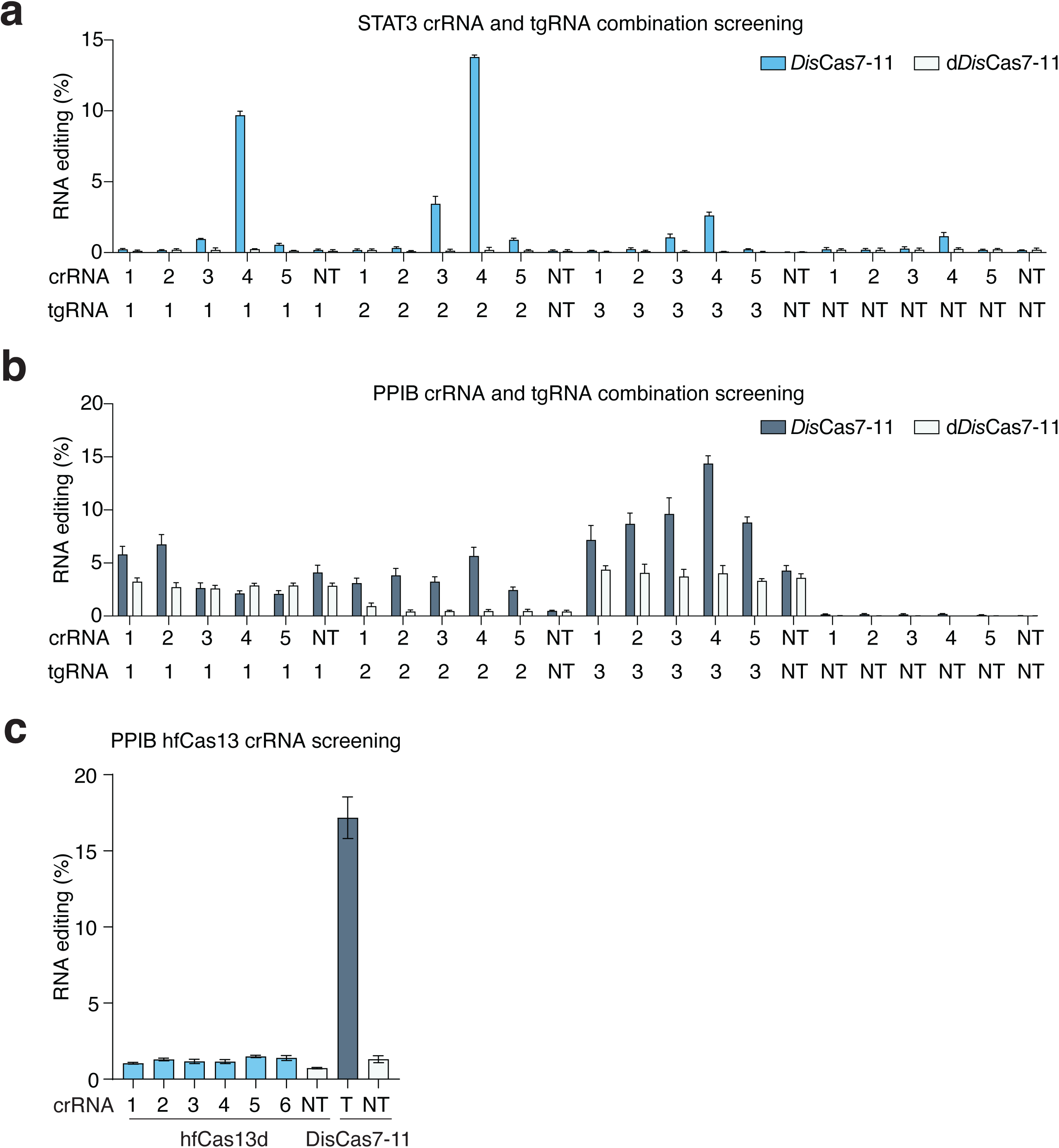
Comprehensive *Dis*Cas7-11 guide and cargo guide tiling for optimizing PRECISE editing efficiency. **(a)** RNA editing rate of PRECISE on the endogenous STAT3 transcript using trans-templates with different *Dis*Cas7-11 guides paired with different cargo guides and either active *Dis*Cas7-11 or d*Dis*Cas7-11. **(b)** RNA editing rate of PRECISE on the endogenous PPIB transcript using trans-templates with different *Dis*Cas7-11 guides paired with different cargo guides and either active *Dis*Cas7-11 or d*Dis*Cas7-11. **(c)** Evaluation of high-fidelity Cas13d (hfCas13d) for trans-splicing at the PPIB transcript compared to PRECISE editing with *Dis*Cas7-11. A panel of targeting guides is compared to a non-targeting guide for each of the nucleases, as indicated.

**Extended Data Figure 4.**
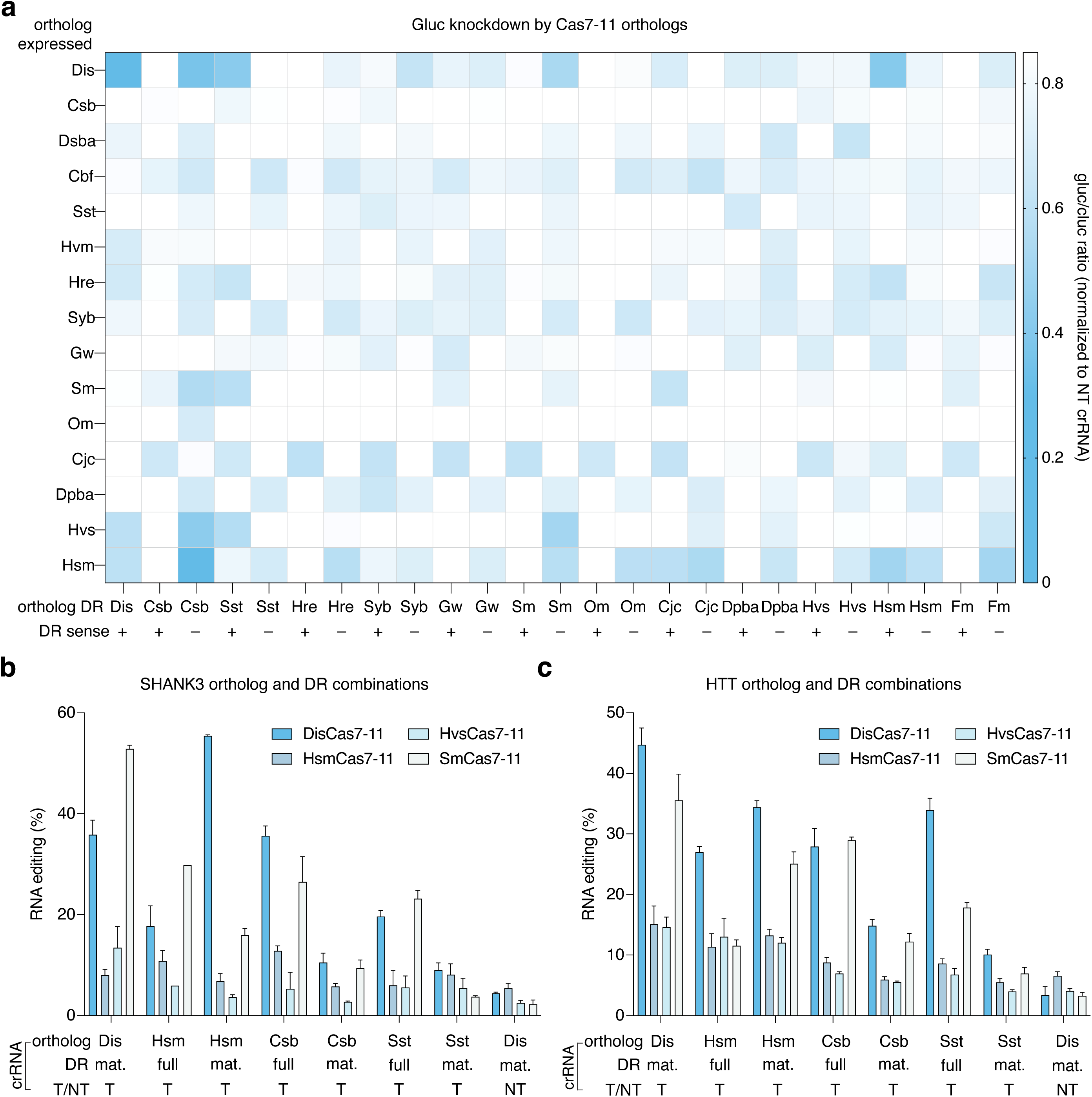
Exploration of Cas7-11 orthologs and their application to PRECISE editing. **(a)** A panel of Cas7-11 orthologs is tested with every known Cas7-11 ortholog crRNA in either the sense or antisense direction for knockdown of G-luciferase activity in HEK293FT cells. **(b)** Top four Cas7-11 orthologs are evaluated for their ability to stimulate 3′ PRECISE editing on the endogenous SHANK3 transcript. Each ortholog is evaluated with DRs from other orthologs that maximally enabled knockdown in part (a). Orthologs are tested with either the full length DRs or predicted mature DRs. **(c)** Top four Cas7-11 orthologs are evaluated for their ability to stimulate 5′ PRECISE editing on the endogenous HTT transcript. Each ortholog is evaluated with DRs from other orthologs that maximally enabled knockdown in part (a). Orthologs are tested with either the full length DRs or predicted mature DRs. Hsm, “Hydrothermal sediment microbial communities from Guaymas Basin, California, USA 4484” Cas7-11; Hvs, “Hydrothermal vent sediment bacterial communities from Southern Trench, Guaymas Basin, Mexico-4870-07-3-4_MG” Cas7-11.

**Extended Data Figure 5.**
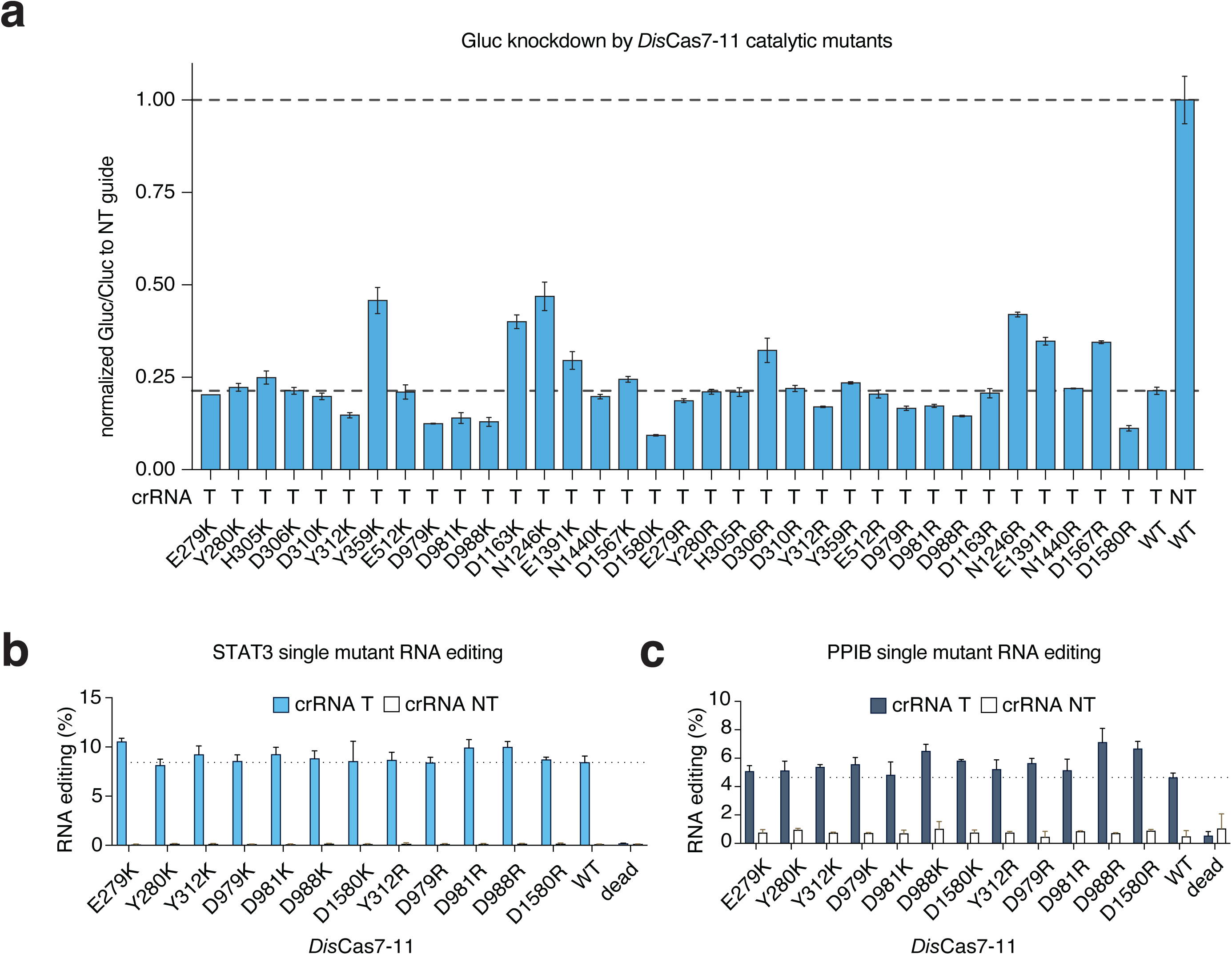
Mutagenesis of *Dis*Cas7-11 for improved catalytic activity and higher trans-splicing rates via PRECISE editing. **(a)** Panel of *Dis*Cas7-11 single protein mutants evaluated for G-luciferase knockdown. G-luciferase activity is normalized to a C-luciferase control and knockdown activity is calculated relative to a non-targeting guide. **(b)** Panel of *Dis*Cas7-11 single protein mutants evaluated for endogenous STAT3 3′ PRECISE editing. A targeting guide is compared to a non-targeting guide. **(c)** Panel of *Dis*Cas7-11 single protein mutants evaluated for endogenous PPIB 3′ PRECISE editing. A targeting guide is compared to a non-targeting guide.

**Extended Data Figure 6.**
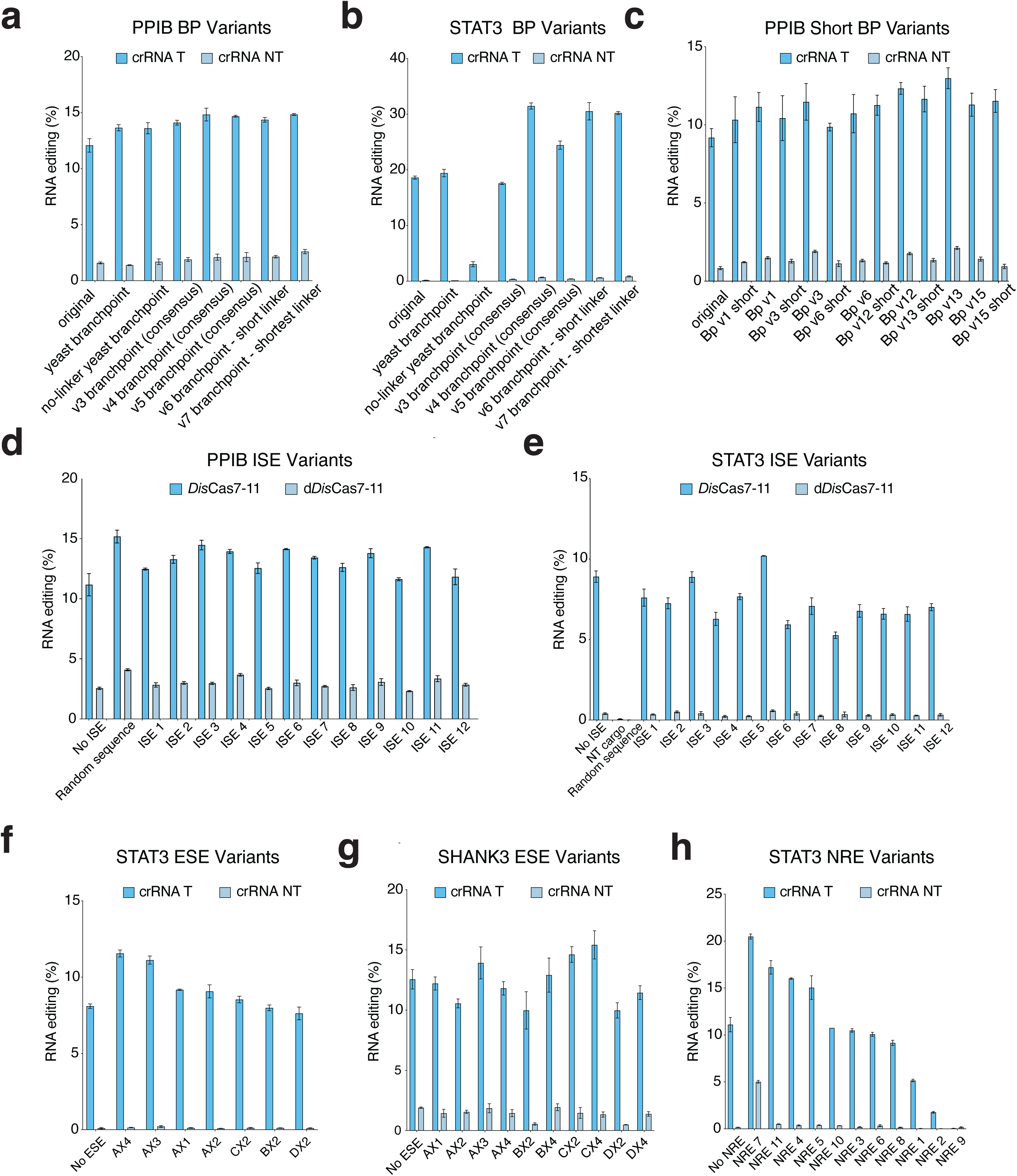
Engineering of trans-template cargos for efficient PRECISE editing. **(a)** Evaluation of 3′ trans-splicing PRECISE editing on the endogenous PPIB transcript with trans-template cargos carrying different branch point sequences. Editing is compared between targeting and non-targeting crRNAs. **(b)** Evaluation of 3′ trans-splicing PRECISE editing on the endogenous STAT3 transcript with trans-template cargos carrying different branch point sequences. Editing is compared between targeting and non-targeting crRNAs. **(c)** Evaluation of 3′ trans-splicing PRECISE editing on the endogenous PPIB transcript with trans-template cargos carrying shorter branch point sequence variants. Editing is compared between targeting and non-targeting crRNAs. **(d)** Evaluation of 3′ trans-splicing PRECISE editing on the endogenous PPIB transcript with trans-template cargos carrying different intron splicing enhancer (ISE) sequences. Editing is compared between e*Dis*Cas7-11 and d*Dis*Cas7-11. **(e)** Evaluation of 3′ trans-splicing PRECISE editing on the endogenous STAT3 transcript with trans-template cargos carrying different ISE sequences. Editing is compared between e*Dis*Cas7-11 and d*Dis*Cas7-11. **(f)** Evaluation of 3′ trans-splicing PRECISE editing on the endogenous STAT3 transcript with trans-template cargos carrying different exon splicing enhancer (ESE) sequences. Editing is compared between targeting and non-targeting guides. **(g)** Evaluation of 3′ trans-splicing PRECISE editing on the endogenous SHANK3 transcript with trans-template cargos carrying different exon splicing enhancer (ESE) sequences. Editing is compared between targeting and non-targeting guides. **(h)** Evaluation of 3′ trans-splicing PRECISE editing on the endogenous STAT3 transcript with trans-template cargos carrying different nuclear retention element (NRE) sequences. Editing is compared between targeting and non-targeting guides.

**Extended Data Figure 7.**
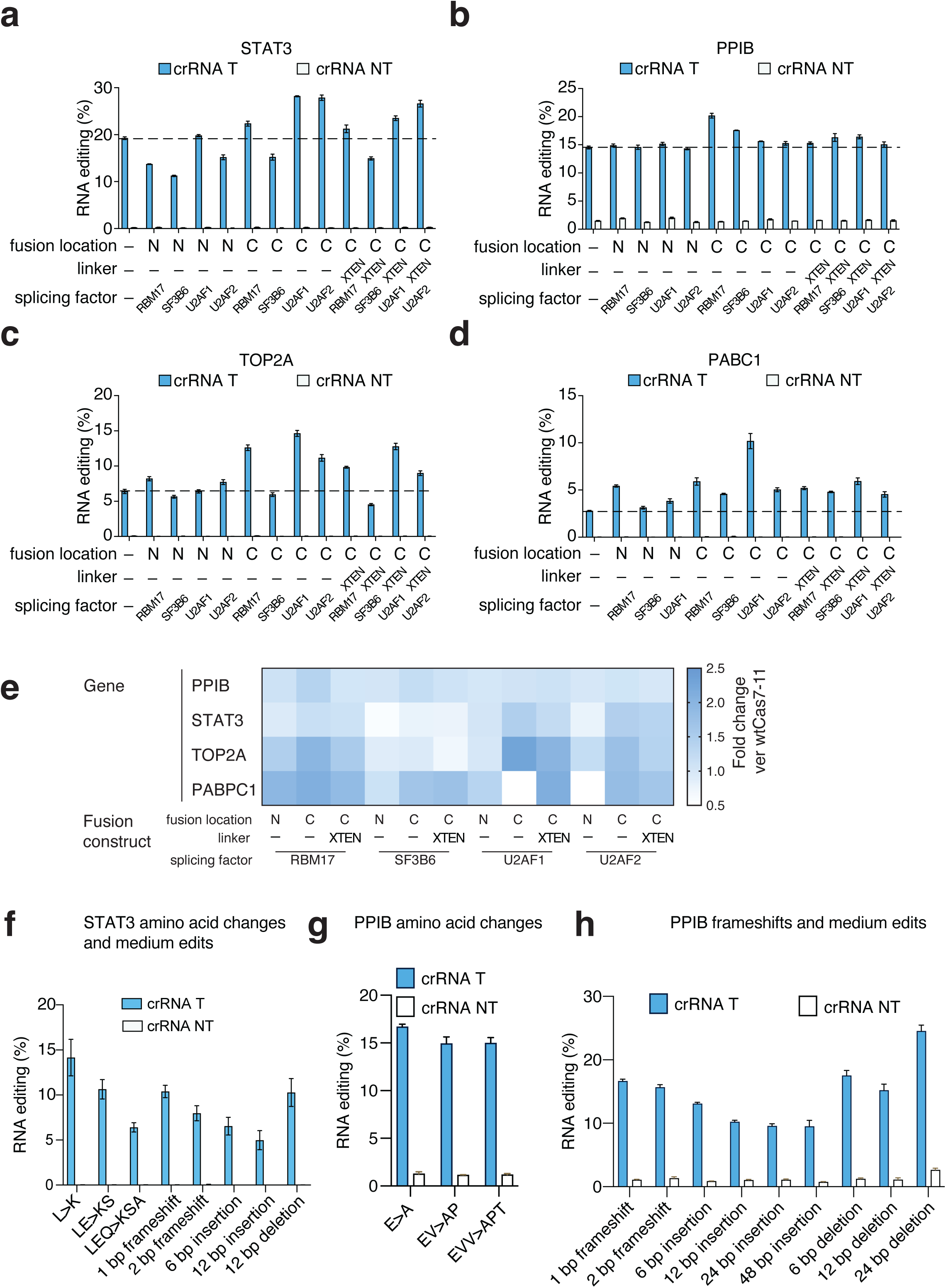
Enhanced trans-splicing efficiencies with splicing factor recruitment and further characterization of edit types possible with PRECISE editing. **(a)** Evaluation of 3′ trans-splicing PRECISE editing on endogenous STAT3 transcript with *Dis*Cas7-11 fusions to different splicing factors. Trans-splicing with a targeting *Dis*Cas7-11 guide is compared to a non-targeting guide. Fusions are on either the N- or C-terminus and either an XTEN linker or no linker is used. **(b)** Evaluation of 3′ trans-splicing PRECISE editing on endogenous PPIB transcript with *Dis*Cas7-11 fusions to different splicing factors. Trans-splicing with a targeting *Dis*Cas7-11 guide is compared to a non-targeting guide. Fusions are on either the N- or C-terminus and either an XTEN linker or no linker is used. **(c)** Evaluation of 3′ trans-splicing PRECISE editing on endogenous TOP2A transcript with *Dis*Cas7-11 fusions to different splicing factors. Trans-splicing with a targeting *Dis*Cas7-11 guide is compared to a non-targeting guide. Fusions are on either the N- or C-terminus and either an XTEN linker or no linker is used. **(d)** Evaluation of 3′ trans-splicing PRECISE editing on endogenous PABPC1 transcript with *Dis*Cas7-11 fusions to different splicing factors. Trans-splicing with a targeting *Dis*Cas7-11 guide is compared to a non-targeting guide. Fusions are on either the N- or C-terminus and either an XTEN linker or no linker is used. **(e)** Heatmap of 3’ PRECISE editing efficiency on 4 endogenous targets, PPIB, STAT3, TOP2A, and PABPC1. Fusions are on either the N- or C-terminus and either an XTEN linker or no linker is used. Rates are expressed as fold changes relative to *Dis*Cas7-11 with no fusion. **(f)** Evaluation of 3′ trans-splicing PRECISE editing on endogenous STAT3 transcript for trans-templates carrying different types of edits, including amino acid mutations, frameshifts, and nucleotide insertions. **(g)** Evaluation of 3′ trans-splicing PRECISE editing on endogenous PPIB transcript for trans-templates carrying different amino acid mutation edits. **(h)** Evaluation of 3′ trans-splicing PRECISE editing on endogenous PPIB transcript for trans-templates carrying different edits, including frameshifts, nucleotide insertions, and nucleotide deletions.

**Extended Data Figure 8.**
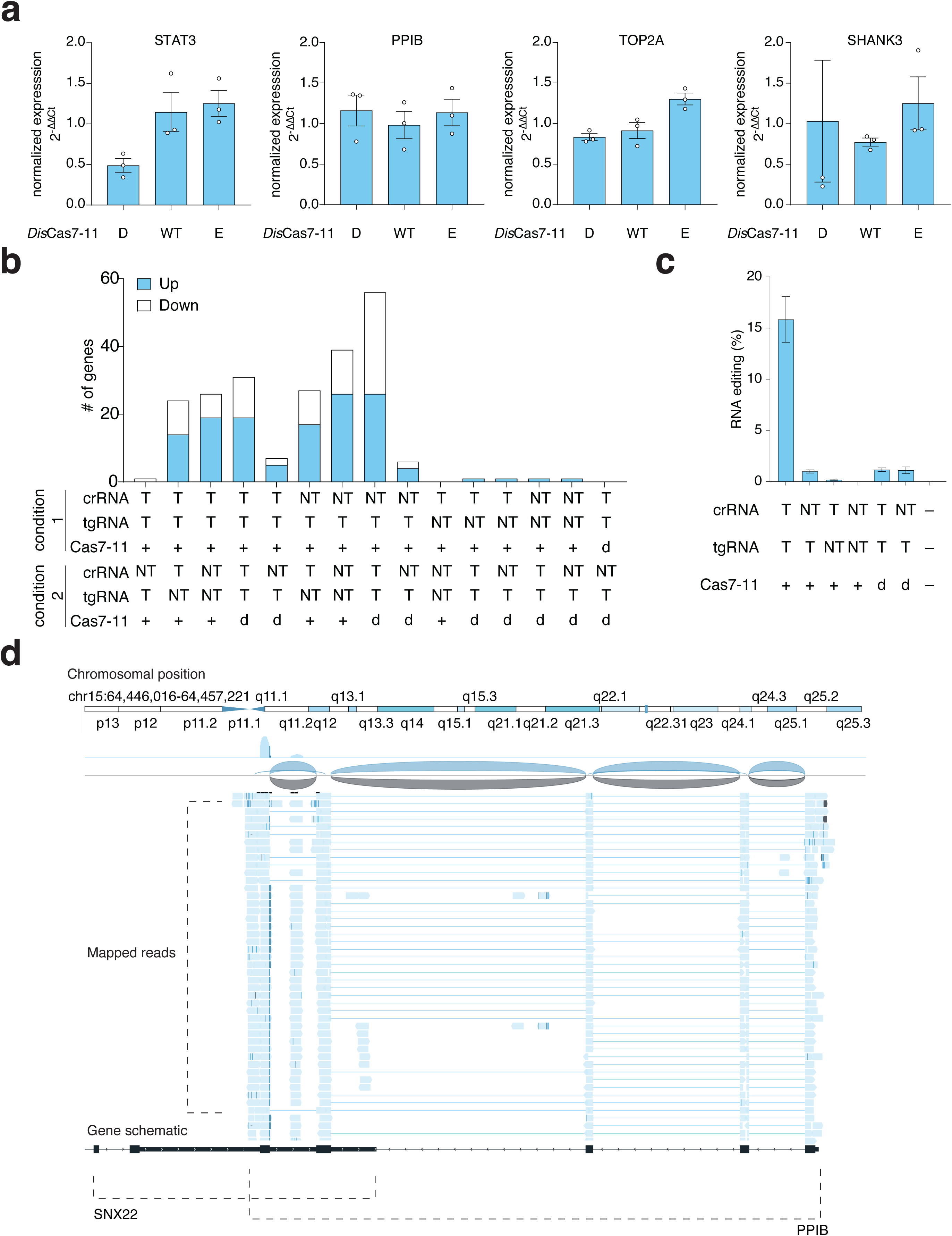
Exploration of transcript perturbations and off-targets due to PRECISE editing. **(a)** Relative expression of four different endogenous transcripts targeted with *Dis*Cas7-11 PRECISE editing, as measured by qPCR. For each replicate, transcript expression is normalized to GAPDH expression and then adjusted transcript expression is normalized to a *Dis*Cas7-11 non-targeting guide control. Conditions shown include PRECISE editing with d*Dis*Cas7-11 (D), active *Dis*Cas7-11 (WT), and e*Dis*Cas7-11 (E). **(b)** Differentially expressed genes for a set of pairwise comparisons from whole-transcriptome RNASeq performed on PPIB-targeting PRECISE edited HEK293FT cells. Numbers of upregulated genes, downregulated genes, and total counts for each comparison are shown. T and NT indicate targeting or non-targeting guides, respectively. (+) and (d) indicate wild-type *Dis*Cas7-11 and d*Dis*Cas7-11, respectively. **(c)** Evaluation of 3′ PRECISE editing on the endogenous PPIB transcript for samples used for whole transcriptome RNA sequencing. T and NT indicate targeting or non-targeting guides, respectively. (+) and (d) indicate wild-type *Dis*Cas7-11 and d*Dis*Cas7-11, respectively. **(d)** Adaptation of Integrative Genomics Viewer^48^ plot of the endogenous PPIB transcript subjected to PRECISE editing, mapping whole-transcriptome RNAseq reads.

**Extended Data Figure 9.**
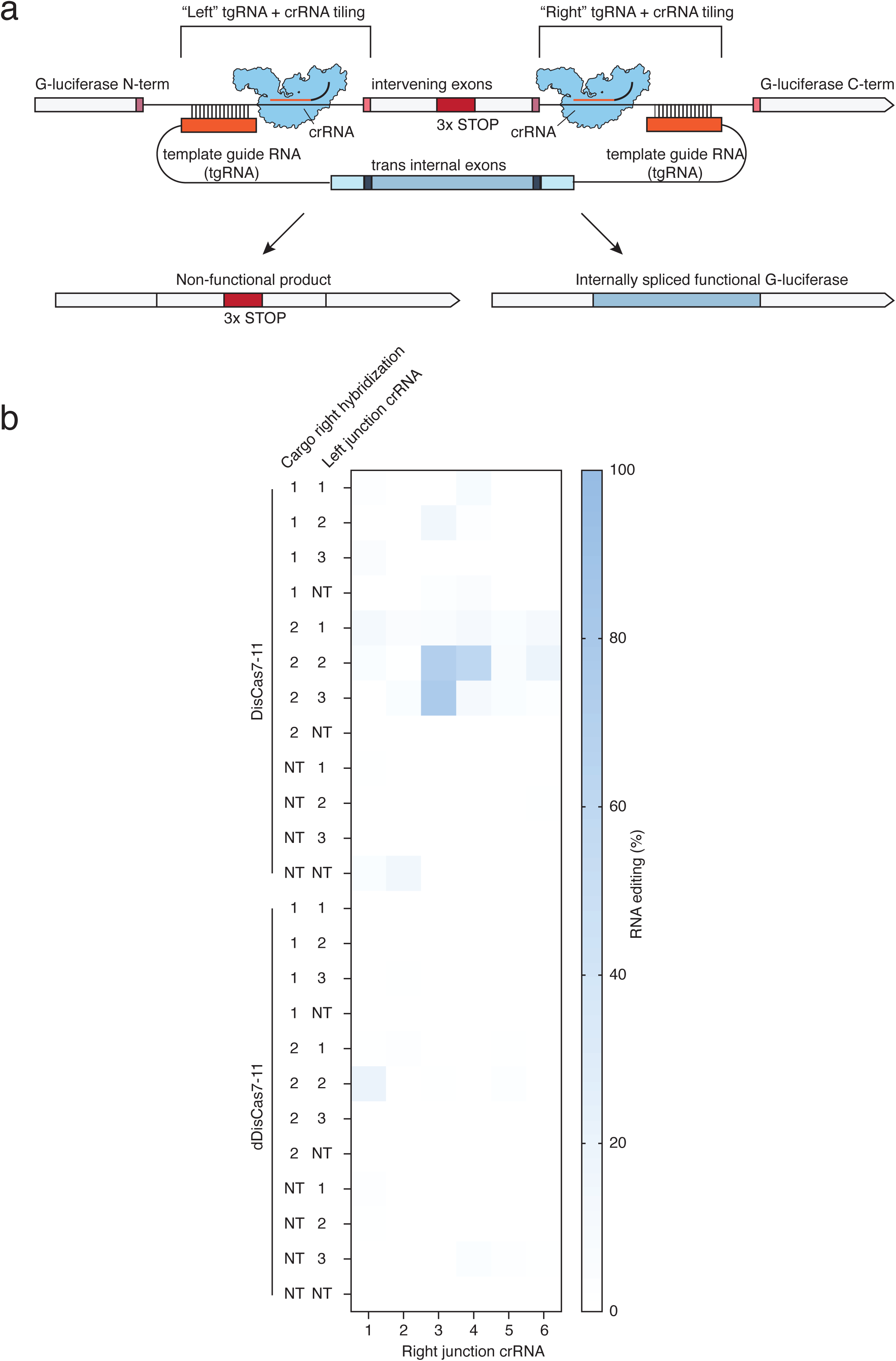
Internal trans-splicing by PRECISE using split-luciferase reporter. **(a)** Schematic of internal exon PRECISE editing showing replacement of G-luciferase exons in a reporter to restore luciferase activity. Approximate positions are shown for both 5′ and 3′ tiling positions of crRNAs and tgRNAs. **(b)** Evaluation of insertion-spanning internal PRECISE editing on overexpressed luciferase splicing reporter reconstituting functional G-luciferase, measured by NGS. Cargos with tgRNAs targeted 5′ and 3′ to the inserted exons or scrambled guide sequences are compared with either active or d*Dis*Cas7-11 with different pairs of crRNAs. Different combinations of 5′ and 3′ intron targeting PRECISE guides are shown. NT (cargo tgRNAs) denotes scrambled tgRNA guide sequence. NT (crRNA) denotes a non-targeting guide for *Dis*Cas7-11.

**Extended Data Figure 10.**
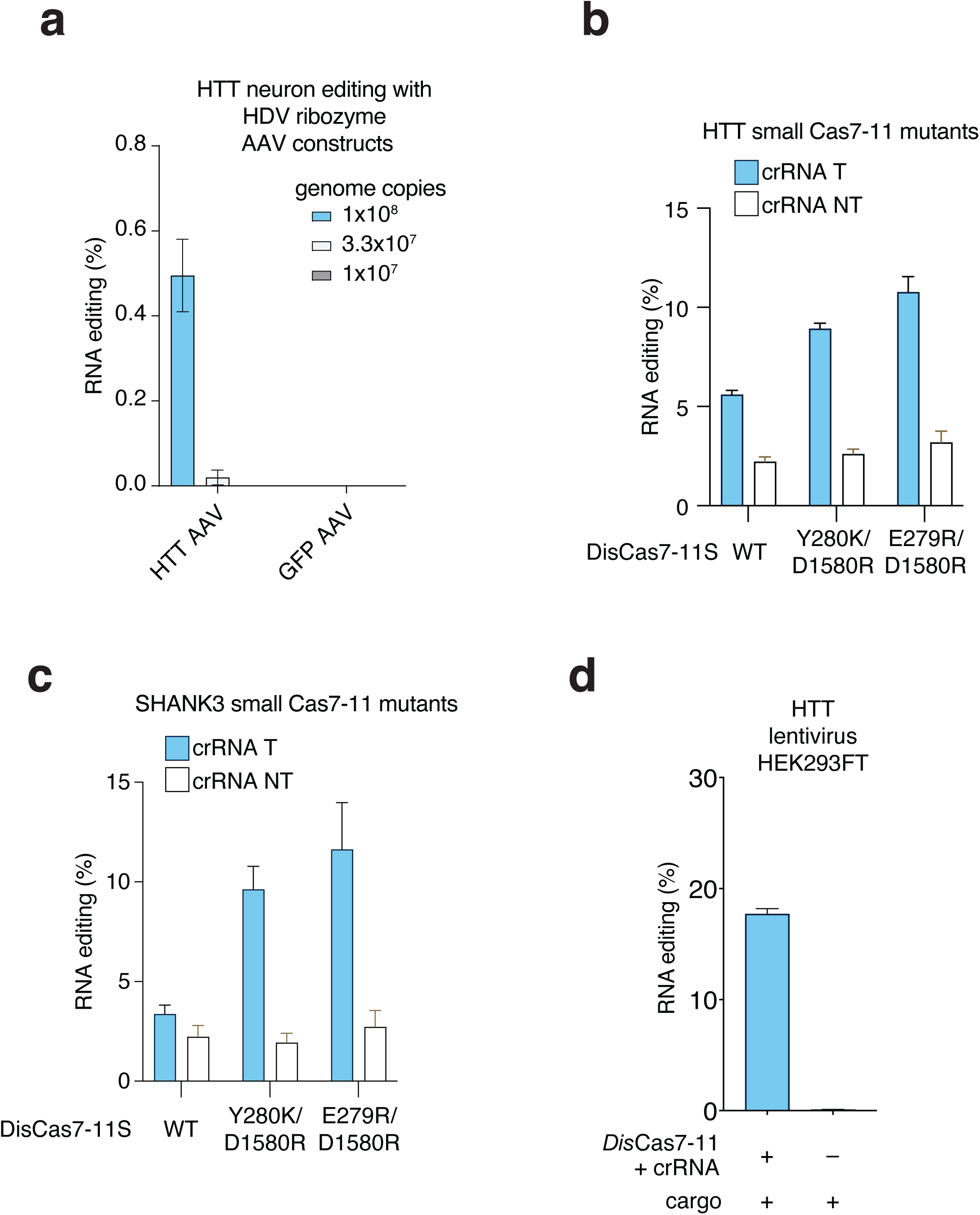
Additional characterization of 5′ trans-splicing with PRECISE editing. **(a)** Evaluation of 5′ PRECISE editing on endogenous HTT transcript in iPSC-derived human neurons by AAV delivery of protein-free cargo with HDV ribozyme. 3 different doses of AAV for the delivery of a targeting cargo or a control GFP vector are compared. **(b)** Evaluation of 5′ PRECISE editing on endogenous HTT transcripts with wild-type small *Dis*Cas7-11 and mutant versions. A targeting guide is compared to a non-targeting guide. **(c)** Evaluation of 3′ PRECISE editing on endogenous SHANK3 transcripts with wild-type small *Dis*Cas7-11 and mutant versions. A targeting guide is compared to a non-targeting guide. **(d)** Demonstration of 5′ PRECISE editing of the HTT transcript with lentiviral delivery of *Dis*Cas7-11, guide, and RNA trans-template in HEK293FT cells. RNA writing into the endogenous HTT transcript with targeting and non-targeting *Dis*Cas7-11 guides is compared.

## Methods

### Cloning of crRNA guides for Cas7-11 and Cas13

Guides for both Cas7-11 and Cas13 were cloned using Golden Gate Assembly. Forward and reverse-complementary sequences with overhangs were ordered as 50 μM single-stranded oligos (QuintaraBio) and annealed via a reaction containing 2 μL of each oligo, 1x Ligation buffer, and 1 μL of T4 PNK (New England Biolabs,M0201S) in Ultrapure water (Gibco, 10977015). Oligo anneals were incubated at 37℃ for 30 minutes, then 95℃ for 5 minutes. Annealed oligos were diluted 1:10 in UltraPure water, then assembled in a reaction containing 0.8 μL of annealed and diluted dsDNA oligo, 25 ng of vector (encoding either Cas13 or Cas7-11 DRs), 2 μL of Rapid Ligation Buffer (Thermo Fisher Scientific, K1422), 0.1 μL of T7 Ligase (New England Biolabs, M0318S), 0.4 μL of Fermentas FD Eco31I (Thermo Fisher Scientific, FD0293), and 0.1 μL of 20 mg/mL BSA (Thermo Fisher Scientific, B14) in UltraPure water, for a total of 20 μL reactions. Reactions were incubated first at 37 °C, then 20 °C for 5 min each for a total of 15 cycles. Post incubation, assembled products were diluted 1:1 with UltraPure water. 2 μL of product was transformed into One Shot™ Stbl3™ Chemically Competent E. coli. Cells, then plated on Agarose plates with 100 μg/mL Ampicillin for overnight outgrowth at 37℃. Single clones were picked from plated into 1 mL of TB media containing 100 μg/mL Ampicillin in 2 mL 96-well plates (VWR, 75870-796) and grown overnight in a 37 °C rotating shaker. Plasmid DNA was purified from cells using a Qiagen 96 Plus Miniprep Kit and Econospin Miniprep Filter plates (Epoch Life Science, 2020-001). Purified plasmids were prepared for sequencing using a Tn5 transposase and tagmentation approach and sequenced using an Illumina MiSeq. Correct clones were verified using Geneious Prime (sequences in Supplementary Table 1).

### Cloning of cargo constructs

Cargo constructs were ordered as eBlock Gene Fragments from IDT DNA and cloned by Gibson Assembly. A pcDNA3.1 - mCardinal cloning backbone (Addgene #51311) was digested using Fermentas FD *Bam*HI and Fermentas FD *Eco*RI (Thermo Fisher Scientific,FD0054, FD0274) with 10x FastDigest Buffer, for a reaction containing 2–5 μg of backbone, 1X FastDigest buffer, and 1 μL of each enzyme in a total reaction size of 20 μL in UltraPure water. Digestions were incubated for 1 hour at 37 ℃ and then diluted 1:5 with UltraPure water before loading on E-gel EX 2% Agarose gels (Invitrogen, G401002). Backbones were purified using the Monarch Gel Extraction Kit (Monarch), then assembled directly with the eBlock cargo constructs at a 1:3 molar ratio of backbone:insert, using 50 ng of backbone and 2.5 μL of HiFi DNA Assembly Mix (New England Bioscience) in a total 5 μL reaction size. Assembly mixes were incubated for 1 hour at 50 °C. Post incubation, assembled products were diluted 1:1 with UltraPure water. 2 μL of product was transformed into One Shot™ Stbl3™ Chemically Competent E. coli. Cells, then plated on Agarose plates with 100 μg/mL Ampicillin for overnight outgrowth at 37 ℃. Single clones were picked from plated into 1 mL of TB media containing 100 μg/mL Ampicillin in 2 mL 96-well plates (VWR) and grown overnight in a 37 °C rotating shaker. Plasmid DNA was purified from cells using a Qiagen 96 Plus Miniprep Kit and Econospin Miniprep Filter plates (Epoch Life Science). Purified plasmids were prepared for sequencing using a Tn5 transposase and tagmentation approach and sequenced using an Illumina MiSeq. Correct clones were verified using Geneious Prime (sequences in Supplementary Table 2).

### Cloning of Cas7-11 effector constructs

Cas7-11 effector constructs were cloned by Gibson Assembly using both PCRs and eBlock Gene Fragments where applicable. *Dis*Cas7-11 mutants were generated as two-insert Gibsons, inserting point mutations in primer overhangs. Oligos for mutagenesis were ordered from QuintaraBio. Cas7-11 orthologs amino acid sequences were reverse-translated to match preferred human codons in SnapGene. Sequences were split into 1–1.5 kilobase fragments and ordered as GenParts from GenScript. Primers were ordered from QuintaraBio to amplify from the ends of each GenPart. Fragments were amplified by PCR in a reaction consisting of 12.5 μL KAPA HiFi HotStart DNA Polymerase (Roche), using 5uM forward and reverse primer pairs, and 12.5 ng of template in a total reaction size of 25 μL. hfRfxCas13 constructs were subcloned from Addgene #190034, while *Lwa*, *Psp*, and *Rfx* Cas13 sequences were ordered as eBlocks before cloning into the expression backbone. Spliceosome proteins for fusion experiments were ordered as eBlocks from IDT DNA. An expression backbone originally containing wild-type Cas7-11 (pDF0506) was digested with Fermentas FD0596 NotI and Fermentas FD1464 AgeI (Thermo Fisher Scientific) with 10x FastDigest Buffer, for a reaction containing 2–5 μg of backbone, 1X FastDigest buffer, and 1 μL of each enzyme in a total reaction size of 20 μL in UltraPure water. Digestions were incubated for 1 hour at 37 ℃ and then diluted 1:5 with UltraPure water before loading on E-gel EX 2% Agarose gels (Invitrogen). Backbones were purified using the Monarch Gel Extraction Kit (Monarch). Both PCR and direct eBlock inserts for Cas7-11, Cas13, and spliceosome constructs were assembled using a 1:3 molar ratio of backbone:insert, using 50 ng of backbone and 2.5 μL of HiFi DNA Assembly Mix (New England Bioscience) in a total 5 μL reaction size. Assembly mixes were incubated for 1 hour at 50 °C. Post incubation, assembled products were diluted 1:1 with UltraPure water. 2 μL of product was transformed into One Shot™ Stbl3™ Chemically Competent E. coli. Cells, then plated on Agarose plates with 100 μg/mL Ampicillin for overnight outgrowth at 37℃. Single clones were picked from plated into 1 mL of TB media containing 100 μg/mL Ampicillin in 2 mL 96-well plates (VWR) and grown overnight in a 37 °C rotating shaker. Plasmid DNA was purified from cells using a Qiagen 96 Plus Miniprep Kit and Econospin Miniprep Filter plates (Epoch Life Science). Purified plasmids were prepared for sequencing using a Tn5 transposase and tagmentation approach and sequenced using an Illumina MiSeq. Correct clones were verified using Geneious Prime (sequences in Supplementary Table 3).

### Mammalian cell culture

#### HEK cells

HEK293FT cells (Invitrogen, R70007) were cultured in Dulbecco’s modified Eagle medium with high glucose, sodium pyruvate and GlutaMAX (Thermo Fisher Scientific, 35050079) and supplemented with 10% (vol/vol) fetal bovine serum (FBS) and 1× penicillin–streptomycin (Thermo Fisher Scientific, 35050079). Cells were maintained at 37 ℃ and 5% CO_2_ throughout all experiments.

#### Neurons

iPSC-derived neurons were generated according to the approach outlined in Tian et al. 2019 “CRISPR Interference-Based Platform for Multimodal Genetic Screens in Human iPSC-Derived Neurons:“. Neurons were plated on black-well 96-well plates for all experiments and maintained in Differentiation media consisting of Dulbecco’s modified Eagle medium supplemented with 0.5X Neurobasal-A, 1X NEAA, 0.5X GlutaMAX, 0.5X B27-VA, 10 ng/mL NT-3, 10 ng/mL BDNF, 1 μg/mL mouse laminin, and 2 μg/mL doxycycline.

#### Fibroblasts

Huntington’s disease-afflicted patient fibroblasts (GM21756)were purchased from the Coriell Institute. They were cultured in Dermal Fibroblast Growth Media (DFGM) supplemented with 10% (v/v) FBS and 1% Penicillin/Streptomycin. Fibroblasts were plated as 5×10^3^ cells/well on 96-well plates for transduction experiments.

### Transfection

HEK293FT cells were plated in the day before in Dulbecco’s modified Eagle medium with high glucose, sodium pyruvate and GlutaMAX (Thermo Fisher Scientific, 35050079) and supplemented with 10% (vol/vol) fetal bovine serum (FBS) and 1× penicillin–streptomycin (Thermo Fisher Scientific, 15140122) at densities from 10–20 k/ well in cell-culture coated 96 well plates (Corning Costar, CLS3628). For the majority of experiments evaluating standard guide and cargo constructs, 10 ng of guide expressing plasmid, 10 ng of cargo expressing plasmid, and 40 ng of Cas7-11 expressing plasmid were cotransfected using Lipofectamine 3000 (Thermo Fisher Scientific, L3000001) according to manufacturer standards.

### Luciferase assays for knockdown and splicing

Culture media containing secreted luciferase was collected 48 h post transfected. *Gaussia luciferase* and *Cypridina luciferase* luminescence was read using the Gaussia Luciferase Assay reagent (Targeting Systems, GAR-2B) and Cypridina Luciferase Assay reagent (Targeting Systems, VLAR-2) respectively, according to the manufacturer’s instructions. Luminescence was measured on a Biotek Synergy Neo 2 reader. *G*. *luciferase* values were normalized against *C. luciferase* readings as a control for transfection variability.

### RNA isolation and cDNA prep

For HEK cell transfections, RNA was harvested from the 96 well plates 72 hours post transfection. Media was removed from the wells, which were then washed with 100 μL of 1X PBS pH 7.4 (Gibco, 10010023). The PBS was then removed and 50 μL of a lysis buffer added, consisting of 9.6 mM Tris–HCl (pH 7.8), 0.5 mM MgCl_2_, 0.44 mM CaCl_2_, 10 μM DTT, 0.1% (wt/vol) Triton X-114, and 3 U/m proteinase K in UltraPure water, pH ~7.8, supplemented with additional DNAse I (Sigma-Aldrich, D2821-50KU) and Proteinase K (New England Biolabs, P8107S). Wells were scraped in the process of lysis buffer addition, and then incubated for 8 minutes exactly before quenching with a stop solution consisting of 1 mM Proteinase K inhibitor, 90 mM EGTA, and 113 μM DTT in UltraPure water. For experiments in fibroblasts or neurons, RNeasy mini kits (Qiagen, 74104) were used. Cells either lysed directly, or lysed after pelleting. RNA was collected from 96-wells and processed according to the manufacturer’s protocol, with 25 μL elution volumes. RNA from all experiments was reverse-transcribed to cDNA using the RevertAid cDNA prep kit (K1621, Thermo Fisher Scientific). For qPCR assays and higher abundance genes, random hexamer and Oligo-dT priming was used. For low abundance genes, specific reverse transcription primers (Supplementary Table 4) binding to the downstream exon or cargo sequence were used. Reverse transcription was performed according to manufacturer protocols.

### Amplicon sequencing for trans-splicing editing

NGS amplicons were designed to span splicing junctions of interest, with 4 staggered forward primers and a single reverse primer per assay (Supplementary Table 4). Sequencing primers were designed with 65 ℃ binding temperatures. Sequencing was performed in two PCR steps, with the first step amplifying the region of interest and the second barcoding the amplicons using Illumina standard adaptors. For some lower TPM genes, additional first-round cycles were used to increase yield. Barcoded wells from the 2nd-round NGS product were pooled by experiment and loaded onto E-gel EX 2% Agarose gels (Invitrogen, G401002) for visualization. Amplicons were purified using the Monarch Gel Extraction Kit (New England Biolabs, T1030) before sequencing using an Illumina MiSeq. Following sequencing, raw reads were analyzed using a custom R script, calculating a percentage by searching for counts of the wildtype and trans-product splicing junctions.

### AAV production and transduction

AAV constructs were produced in HEK293FT cells cultured in T225 flasks by transfection of 1:1:1 molar ratios of helper plasmid, capsid, and transfer plasmids (per construct), totalling 90 μg of DNA. PEI was used for all transfections, and media was changed on transfected cells 4–6 hours post transfection. 48 hours post transfection, the supernatant from transfected flasks was collected and briefly centrifuged at 1000 x g for 5 minutes to pellet cell debris. Clarified supernatant was then passed through a 0.45 μm syringe filter before centrifuging through 100 kDa MWCO Amicon filters to concentrate (Millipore Sigma, UFC910024). Two washes with PBS were also performed in an Amicon filter. The concentrated AAV was collected and a small fraction used to estimate viral genome titers by qPCR as follows: first, a DNAse digestion was performed using 1 μL of concentrated AAV, 2 μL of DNAse I buffer, and 0.5 μL of DNase I (New England Biolabs, M0303S) in UltraPure water for a total reaction volume of 20 μL. The DNAse digestion was incubated at 37 °C for 1 hour, then 75 °C for 15 minutes. 5 μL of the DNAse digestion product was then treated with 1 μL of Proteinase K (New England Biolabs, P8107S) in UltraPure water for a total reaction volume of 20 μL. The Proteinase K treatment was incubated at 50 °C for 30 minutes, then 98 °C for 10 minutes. qPCR was performed using custom primers binding within the intra-ITR CMV promoter of all constructs, using Fast SYBR Green Master Mix (Applied Biosystems: 4385612). 0.2 μL each of 50 μM forward and reverse primers were combined with 4 μL of the treated sample, along with 10 μL of 2x SYBR Green mix in UltraPure water for a total reaction volume of 20 μL. A serial dilution was prepared for each transfer plasmid from 1:10 to 1:10^11^ and SYBR reactions were set up for each dilution. Reactions were run on the CFX384 Touch Real-Time PCR System (BioRad). Copy numbers were calculated according to the standard curves generated by the serial dilution reactions.

### Lentivirus production and transduction

Lentiviruses were produced in HEK293FT cells cultured in T225 flasks by cotransfection of 30μg of packaging plasmid (psPAX2, Addgene #12260), 30 μg of envelope plasmid (VSV-G, Addgene #8454), and 30 μg of transfer plasmid using 270 μl of polyethylene imine (PEI). Media was changed 6–8 hours post transfection with fresh D10. Media containing lentiviruses were harvested after 48 h of transfection, briefly centrifuged at 1000 x g for 5 minutes to pellet cell debris, and filtered through 0.45 μm vacuum filters. Then ultracentrifuged for 2 h at 120,000 x g, and resuspended in PBS to a total volume of 200–300 μL. HEK293FT cells, hHD fibroblasts, and iPSC-derived neurons were plated on 96-well plates as 2×10^4^, 5×10^3^, 1×10^6^ cells/well, respectively. They were infected with concentrated lentiviruses in DMEM 10% FBS, in the presence of polybrene with a final concentration of 10 μg/mL.

### Protein quantification by western blot

All cargo constructs include an epitope tag or tandem tag (FLAG or 3xFLAG) for assay by anti-FLAG primary antibodies (Cell Signaling Technology, 2368). HEK293FT cells were transfected at 24 well, 6 well, (using Lipofectamine 3000) or T25 scales (using PEI) for western blot experiments using manufacturer suggested DNA amounts. 72 hours post transfection, media was removed from plates or flasks, washed once with 1X PBS (Gibco, 10010023) and replaced with 100 μL (for 24 wells), 250 μL (for 6 wells), or 1mL (for T25 experiments) of protein lysis buffer consisting of 1% Triton X-100, 50 mM Tris pH 8, 150 mM NaCl, and 1X Protease Inhibitor Cocktail (Thermo Fisher Scientific, 78429). Cells were lysed on ice for approximately 10 minutes, then collected in Eppendorfs and centrifuged at 16,000 x g for 15 minutes to pellet cell debris. The supernatant was collected and concentration determined by Pierce BCA assay (Thermo Fisher Scientific 23227). Concentrations were normalized and equal amounts from each sample run on Bio-Rad 4–20% Mini-PROTEAN gels (4561094) using a BioRad Tetra at 150-200V. Gels were transferred to nitrocellulose membranes via an iBlot-2 transfer device (Thermo Fisher Scientific), blocked for 1h at RT using Intercept PBS LI-COR Blocking Buffer (LI-COR, 927-70000), and incubated overnight with primary antibody at 4 °C. Membranes were washed 4 x 5 minutes in 0.1% Tween 20 in PBS on a rotating shaker before incubation with secondary antibodies. Fluorescent dye-labeled antibodies (IRDye® 680RD Goat anti-Mouse IgG Secondary Antibody, IRDye® 800CW Goat anti-Mouse IgG Secondary Antibody, LI-COR, 926-68071, 926-32210) were used for as secondaries, incubating at room temperature for 1 hour. Membranes were washed 4 x 5 minutes in 0.1% Tween 20 in PBS after incubation with secondary antibody, then imaged using a LI-COR Odyssey Scanner.

### qPCR quantification for knockdown

HEK cells were transfected in a 96-well plate, RNA was harvested 72 h post-transfection and cDNA was synthesized as previously described in the RNA isolation cDNA prep section. Reactions were set up using Fast SYBR Green Master Mix (Applied Biosystems: 4385612) and qRT-PCR gene-specific primers (Supplementary Table 4.) according to manufacturing protocols. Reactions were run on the CFX384 Touch Real-Time PCR System (BioRad). Ct values and melt curves of the cDNA samples were obtained after 40 cycles of amplification. The Ct values for genes of interest were normalized to ACTB, and expressions of genes are represented as 2-[ΔΔCt] for fold change over the control condition.

### RNAseq perturbation analysis

An RNeasy kit (Qiagen) was used to purify RNA for RNAseq perturbation and off-target analysis. Samples were normalized to an input of 150 ng before preparation with a NEBNext® Ultra™ II RNA Library Prep (Magnetic Bead) kit (New England Biolabs). The polyA mRNA protocol was used for all preps. Samples were barcoded using Illumina i5 and i7 adaptors from NEBNext® Multiplex Oligos for Illumina (Dual Index Primers Set 1) kit (New England Biolabs) and sequenced using a 150-cycle NextSeq 500/550 High Output Kit v2.5 with 75 forward and 75 reverse reads. The output samples were processed using Galaxy (version 23.1.rc1. FASTQ trimming was performed by Cutadapt, using an R1 minimum length cutoff of 20 and quality cutoff of 20. Trimmed reads were mapped to the human genome release 19 (GRCh37.p13) using STAR with GeneCounts. featureCounts was then used to generate reads per gene, specifying the input as unstranded, using the same GFF file for hg19 used by STAR, GFF feature type filter as “exon”, GFF gene identifier as “gene_id”, and input as “Paired end, count as single fragment”. Read filtering minimum mapping quality per read was set to 10. Differentially expressed genes were calculated using DeSeq2 on the count data. Volcano plots were generated using Galaxy and colored based on a 0.01 FDR.

### Off-target analysis for splicing

Transcriptome reads processed with the arriba workflow^49^. After alignment of reads to the hg38 cDNA reference with STAR aligner^50^, the resulting aligned reads were processed using arriba with a mismatch p-value cutoff of 0.05. Off-target fusion mapping with a 3′ PPIB mapping were compiled, and the significance was calculated by a Student’s t-test across all replicates.

