## Supplementary Information for "Programmable RNA writing with trans-splicing"

[2] Department of Chemistry and Biotechnology, Graduate School of Engineering,
The University of Tokyo
7-3-1 Hongo, Bunkyo-ku, Tokyo 113-8656, Japan

[3] Structural Biology Division,
Research Center for Advanced Science and Technology,
The University of Tokyo
4-6-1 Komaba, Meguro-ku, Tokyo 153-8904, Japan

[4] Department of Biological Sciences, Graduate School of Science,
The University of Tokyo
7-3-1 Hongo, Bunkyo-ku, Tokyo 113-0033, Japan

[5] Inamori Research Institute for Science
620 Suiginya-cho, Shimogyo-ku, Kyoto 600-8411, Japan

[6] Japan Science and Technology Agency, Core Research for Evolutional Science and Technology
4-1-8 Honcho, Kawaguchi, Saitama, 332-0012, Japan

‡ These authors jointly supervised the work.

*These authors contributed equally.

### Supplementary Tables

#### Supplementary Table 1: Guide sequences.

| | | |
|---|---|---|
| 1 | *Dis*Cas7-11 USF1 3' targeting guide | GTTGATGTCACGGAACccaaaagtaggttcacactttggacctcatt |
| 2 | *Dis*Cas7-11 RPL41 3' targeting guide | GTTGATGTCACGGAACtccctcccacattaaatcaaacgtccacata |
| 3 | *Dis*Cas7-11 PAPBC1 3' targeting guide | GTTGATGTCACGGAACgtttcttccctcaaatgaaagtataaattgt |
| 4 | *Dis*Cas7-11 PPIB 3' targeting guide 4 | GTTGATGTCACGGAACgccaagggtgaggaggaggaagagggtgacc |
| 5 | *Dis*Cas7-11 TOP2A 3' targeting guide | GTTGATGTCACGGAACtttaacaatatttattgagcacttgctatgt |
| 6 | *Dis*Cas7-11 SHANK3 3' targeting guide | GTTGATGTCACGGAACaggcgccgggttggcaagtgggcagggaaca |
| 7 | *Dis*Cas7-11 STAT3 3' targeting guide 4 | GTTGATGTCACGGAACagaaaatataaagtttctgaggagaattcaa |
| 8 | *Dis*Cas7-11 NT guide | GTTGATGTCACGGAACggtaatgcctggcttgtcgacgcatagtctg |
| 9 | *Dis*Cas7-11 HTT 5' targeting guide | GTTGATGTCACGGAACtgacagccattggtgaactcgtgccctgtgc |
| 10 | *Dis*Cas7-11 PABPC1 5' guide 1 | GTTGATGTCACGGAACagtgtgtgatacttgaaaggtctagccatct |
| 11 | *Dis*Cas7-11 PABPC1 5' guide 2 | GTTGATGTCACGGAACctttatacaacttaggtcccacactagtgtg |
| 12 | *Dis*Cas7-11 PABPC1 5' guide 3 | GTTGATGTCACGGAACcgggggcttctggtatttgtctttgctttat |
| 13 | *Dis*Cas7-11 PABPC1 5' guide 4 | GTTGATGTCACGGAACgaattcttttatatgtgagaaatttcggggg |
| 14 | *Dis*Cas7-11 PPIB 5' guide 1 | GTTGATGTCACGGAACctcataggatttttaccgtcaccaaaatcag |
| 15 | *Dis*Cas7-11 PPIB 5' guide 2 | GTTGATGTCACGGAACgaaaagggtctggagctttcattagattctc |
| 16 | *Dis*Cas7-11 PPIB 5' guide 3 | GTTGATGTCACGGAACgggtatagataagcatgttttccaagaaaag |
| 17 | *Dis*Cas7-11 PPIB 5' guide 4 | GTTGATGTCACGGAACatattgttatcctgtagtccaaggagggtat |
| 18 | *Dis*Cas7-11 RPL41 5' guide 1 | GTTGATGTCACGGAACgaaaaacgtttgagtgttttctccctggagc |
| 19 | *Dis*Cas7-11 RPL41 5' guide 2 | GTTGATGTCACGGAACgaaatcttaaaagatctttaggagaaaaacg |
| 20 | *Dis*Cas7-11 RPL41 5' guide 3 | GTTGATGTCACGGAACcaaaacttaggagaaacatttggtttggaaa |
| 21 | *Dis*Cas7-11 RPL41 5' guide 4 | GTTGATGTCACGGAACcccaggaggagggaagttccttggacaaaac |
| 22 | *Lwa*Cas13 Guide 1 | ATTTAGACTACCCCAAAAACGAAGGGGACTtggctcacgcctgtaatgccagcactttgag |
| 23 | *Lwa*Cas13 Guide 2 | ATTTAGACTACCCCAAAAACGAAGGGGACTgtatccccaagagaaggctccctgttggcca |
| 24 | *Lwa*Cas13 Guide 3 | ATTTAGACTACCCCAAAAACGAAGGGGACTcaaaacctcaaaaaagatacatgcaggacct |
| 25 | *Lwa*Cas13 Guide 4 | ATTTAGACTACCCCAAAAACGAAGGGGACTagaaaatataaagtttctgaggagaattcaa |
| 26 | *Lwa*Cas13 Guide 5 | ATTTAGACTACCCCAAAAACGAAGGGGACTacaaaaaaacagaagtaaagaaagatttcct |
| 27 | *Lwa*Cas13 Guide NT | ATTTAGACTACCCCAAAAACGAAGGGGACTggtccgctgccgttcgcttgggacatcctgt |
| 28 | *Psp*Cas13 Guide 1 | GTTGTGGAAGGTCCAGTTTTGAGGGGCTATtggctcacgcctgtaatgccagcactttgag |
| 29 | *Psp*Cas13 Guide 2 | GTTGTGGAAGGTCCAGTTTTGAGGGGCTATgtatccccaagagaaggctccctgttggcca |
| 30 | *Psp*Cas13 Guide 3 | GTTGTGGAAGGTCCAGTTTTGAGGGGCTATcaaaacctcaaaaaagatacatgcaggacct |
| 31 | *Psp*Cas13 Guide 4 | GTTGTGGAAGGTCCAGTTTTGAGGGGCTATagaaaatataaagtttctgaggagaattcaa |
| 32 | *Psp*Cas13 Guide 5 | GTTGTGGAAGGTCCAGTTTTGAGGGGCTATacaaaaaaacagaagtaaagaaagatttcct |
| 33 | *Psp*Cas13 Guide NT | GTTGTGGAAGGTCCAGTTTTGAGGGGCTATggtccgctgccgttcgcttgggacatcctgt |

| | | |
|---|---|---|
| 34 | *Rfx*Cas13 Guide 1 | AACCCCTACCAACTGGTCGGGGTTTGAAACtggctcacgcctgtaatgccagcactttgag |
| 35 | *Rfx*Cas13 Guide 2 | AACCCCTACCAACTGGTCGGGGTTTGAAACgtatccccaagagaaggctccctgttggcca |
| 36 | *Rfx*Cas13 Guide 3 | AACCCCTACCAACTGGTCGGGGTTTGAAACcaaaacctcaaaaaagatacatgcaggacct |
| 37 | *Rfx*Cas13 Guide 4 | AACCCCTACCAACTGGTCGGGGTTTGAAACagaaaatataaagtttctgaggagaattcaa |
| 38 | *Rfx*Cas13 Guide 5 | AACCCCTACCAACTGGTCGGGGTTTGAAACacaaaaaaacagaagtaaagaaagatttcct |
| 39 | *Rfx*Cas13 Guide NT | AACCCCTACCAACTGGTCGGGGTTTGAAACggtccgctgccgttcgcttgggacatcctgt |

#### Supplementary Table 2: Cargo Sequences.

| | | |
|---|---|---|
| 1 | STAT3 variation substitute A to T | gaacagaaaatataaagtttctgaggagaattcaaatgaagccaaaacctcaaaaaagatacatgcaggacctgcaggcagtatccccaagagaaggctccctgttggccaggtgcagtggctcacgcctgtaatgccagcactttgagaacgaggaattctcttctttttttttctgcaggTtctagaacagaaaatgaaagtggtagagaatctccaggatgactttgatttcaactataaaaccctcaagagtcaaggaGACTACAAAGACGATGACGACAAGTAA |
| 2 | STAT3 variation substitute A to C | gaacagaaaatataaagtttctgaggagaattcaaatgaagccaaaacctcaaaaaagatacatgcaggacctgcaggcagtatccccaagagaaggctccctgttggccaggtgcagtggctcacgcctgtaatgccagcactttgagaacgaggaattctcttctttttttttctgcaggCtctagaacagaaaatgaaagtggtagagaatctccaggatgactttgatttcaactataaaaccctcaagagtcaaggaGACTACAAAGACGATGACGACAAGTAA |
| 3 | STAT3 variation substitute A to G | gaacagaaaatataaagtttctgaggagaattcaaatgaagccaaaacctcaaaaaagatacatgcaggacctgcaggcagtatccccaagagaaggctccctgttggccaggtgcagtggctcacgcctgtaatgccagcactttgagaacgaggaattctcttctttttttttctgcaggGtctagaacagaaaatgaaagtggtagagaatctccaggatgactttgatttcaactataaaaccctcaagagtcaaggaGACTACAAAGACGATGACGACAAGTAA |
| 4 | STAT3 variation substitute T to A | gaacagaaaatataaagtttctgaggagaattcaaatgaagccaaaacctcaaaaaagatacatgcaggacctgcaggcagtatccccaagagaaggctccctgttggccaggtgcagtggctcacgcctgtaatgccagcactttgagaacgaggaattctcttctttttttttctgcaggaActagaacagaaaatgaaagtggtagagaatctccaggatgactttgatttcaactataaaaccctcaagagtcaaggaGACTACAAAGACGATGACGACAAGTAA |
| 5 | STAT3 variation substitute T to C | gaacagaaaatataaagtttctgaggagaattcaaatgaagccaaaacctcaaaaaagatacatgcaggacctgcaggcagtatccccaagagaaggctccctgttggccaggtgcagtggctcacgcctgtaatgccagcactttgagaacgaggaattctcttctttttttttctgcaggaCctagaacagaaaatgaaagtggtagagaatctccaggatgactttgatttcaactataaaaccctcaagagtcaaggaGACTACAAAGACGATGACGACAAGTAA |
| 6 | STAT3 variation substitute T to G | gaacagaaaatataaagtttctgaggagaattcaaatgaagccaaaacctcaaaaaagatacatgcaggacctgcaggcagtatccccaagagaaggctccctgttggccaggtgcagtggctcacgcctgtaatgccagcactttgagaacgaggaattctcttctttttttttctgcaggaGctagaacagaaaatgaaagtggtagagaatctccaggatgactttgatttcaactataaaaccctcaagagtcaaggaGACTACAAAGACGATGACGACAAGTAA |
| 7 | STAT3 variation substitute C to T | gaacagaaaatataaagtttctgaggagaattcaaatgaagccaaaacctcaaaaaagatacatgcaggacctgcaggcagtatccccaagagaaggctccctgttggccaggtgcagtggctcacgcctgtaatgccagcactttgagaacgaggaattctcttctttttttttctgcaggatTtagaacagaaaatgaaagtggtagagaatctccaggatgactttgatttcaactataaaaccctcaagagtcaaggaGACTACAAAGACGATGACGACAAGTAA |
| 8 | STAT3 variation substitute C to G | gaacagaaaatataaagtttctgaggagaattcaaatgaagccaaaacctcaaaaaagatacatgcaggacctgcaggcagtatccccaagagaaggctccctgttggccaggtgcagtggctcacgcctgtaatgccagcactttgagaacgaggaattctcttctttttttttctgcaggatGtagaacagaaaatgaaagtggtagagaatctccaggatgactttgatttcaactataaaaccctcaagagtcaaggaGACTACAAAGACGATGACGACAAGTAA |
| 9 | STAT3 variation substitute C to A | gaacagaaaatataaagtttctgaggagaattcaaatgaagccaaaacctcaaaaaagatacatgcaggacctgcaggcagtatccccaagagaaggctccctgttggccaggtgcagtggctcacgcctgtaatgccagcactttgagaacgaggaattctcttctttttttttctgcaggatAtagaacagaaaatgaaagtggtagagaatctccaggatgactttgatttcaactataaaaccctcaagagtcaaggaGACTACAAAGACGATGACGACAAGTAA |
| 10 | STAT3 variation substitute G to A | gaacagaaaatataaagtttctgaggagaattcaaatgaagccaaaacctcaaaaaagatacatgcaggacctgcaggcagtatccccaagagaaggctccctgttggccaggtgcagtggctcacgcctgtaatgccagcactttgagaacgaggaattctcttctttttttttctgcagAatctagaacagaaaatgaaagtggtagagaatctccaggatgactttgatttcaactataaaaccctcaagagtcaaggaGACTACAAAGACGATGACGACAAGTAA |
| 11 | STAT3 variation substitute G to T | gaacagaaaatataaagtttctgaggagaattcaaatgaagccaaaacctcaaaaaagatacatgcaggacctgcaggcagtatccccaagagaaggctccctgttggccaggtgcagtggctcacgcctgtaatgccagcactttgagaacgaggaattctcttctttttttttctgcagTatctagaacagaaaatgaaagtggtagagaatctccaggatgactttgatttcaactataaaaccctcaagagtcaaggaGACTACAAAGACGATGACGACAAGTAA |
| 12 | STAT3 variation substitute G to C | gaacagaaaatataaagtttctgaggagaattcaaatgaagccaaaacctcaaaaaagatacatgcaggacctgcaggcagtatccccaagagaaggctccctgttggccaggtgcagtggctcacgcctgtaatgccagcactttgagaacgaggaattctcttctttttttttctgcagCatctagaacagaaaatgaaagtggtagagaatctccaggatgactttgatttcaactataaaaccctcaagagtcaaggaGACTACAAAGACGATGACGACAAGTAA |
| 13 | STAT3 variation 1aa change | gaacagaaaatataaagtttctgaggagaattcaaatgaagccaaaacctcaaaaaagatacatgcaggacctgcaggcagtatccccaagagaaggctccctgttggccaggtgcagtggctcacgcctgtaatgccagcactttgagaacgaggaattctcttctttttttttctgcaggatAAGgaacagaaaatgaaagtggtagagaatctccaggatgactttgatttcaactataaaaccctcaagagtcaaggaGACTACAAAGACGATGACGACAAGTAA |
| 14 | STAT3 variation 2aa change | gaacagaaaatataaagtttctgaggagaattcaaatgaagccaaaacctcaaaaaagatacatgcaggacctgcaggcagtatccccaagagaaggctccctgttggccaggtgcagtggctcacgcctgtaatgccagcactttgagaacgaggaattctcttctttttttttctgcaggatAAGTCAcagaaaatgaaagtggtagagaatctccaggatgactttgatttcaactataaaaccctcaagagtcaaggaGACTACAAAGACGATGACGACAAGTAA |
| 15 | STAT3 variation 3aa change | gaacagaaaatataaagtttctgaggagaattcaaatgaagccaaaacctcaaaaaagatacatgcaggacctgcaggcagtatccccaagagaaggctccctgttggccaggtgcagtggctcacgcctgtaatgccagcactttgagaacgaggaattctcttctttttttttctgcaggatAAGTCAGCAaaaatgaaagtggtagagaatctccaggatgactttgatttcaactataaaaccctcaagagtcaaggaGACTACAAAGACGATGACGACAAGTAA |
| 16 | STAT3 variation frameshift correction 1bp | gaacagaaaatataaagtttctgaggagaattcaaatgaagccaaaacctcaaaaaagatacatgcaggacctgcaggcagtatccccaagagaaggctccctgttggccaggtgcagtggctcacgcctgtaatgccagcactttgagaacgaggaattctcttctttttttttctgcaggaAtctagaacagaaaatgaaagtggtagagaatctccaggatgactttgatttcaactataaaaccctcaagagtcaaggaGACTACAAAGACGATGACGACAAGTAA |
| 17 | STAT3 variation frameshift correction 2bp | gaacagaaaatataaagtttctgaggagaattcaaatgaagccaaaacctcaaaaaagatacatgcaggacctgcaggcagtatccccaagagaaggctccctgttggccaggtgcagtggctcacgcctgtaatgccagcactttgagaacgaggaattctcttctttttttttctgcaggaAGtctagaacagaaaatgaaagtggtagagaatctccaggatgactttgatttcaactataaaaccctcaagagtcaaggaGACTACAAAGACGATGACGACAAGTAA |
| 18 | STAT3 variation 6 bp insertion | gaacagaaaatataaagtttctgaggagaattcaaatgaagccaaaacctcaaaaaagatacatgcaggacctgcaggcagtatccccaagagaaggctccctgttggccaggtgcagtggctcacgcctgtaatgccagcactttgagaacgaggaattctcttctttttttttctgcaggatTTGCACctagaacagaaaatgaaagtggtagagaatctccaggatgactttgatttcaactataaaaccctcaagagtcaaggaGACTACAAAGACGATGACGACAAGTAA |
| 19 | STAT3 variation 12 bp insertion | gaacagaaaatataaagtttctgaggagaattcaaatgaagccaaaacctcaaaaaagatacatgcaggacctgcaggcagtatccccaagagaaggctccctgttggccaggtgcagtggctcacgcctgtaatgccagcactttgagaacgaggaattctcttctttttttttctgcaggatTTGCACATTGCGctagaacagaaaatgaaagtggtagagaatctccaggatgactttgatttcaactataaaaccctcaagagtcaaggaGACTACAAAGACGATGACGACAAGTAA |

| | | |
|---|---|---|
| 20 | STAT3 variation 24 bp insertion | gaacagaaaatataaagtttctgaggagaattcaaatgaagccaaaacctcaaaaaagatacatgcaggacctgcaggcagtatccccaagagaaggctccctgttggccaggtgcagtggctcacgcctgtaatgccagcactttgagaacgaggaattctcttcttttttttctgcaggatTTGCACATTGCGTCAACTCATAAGctagaacagaaaatgaaagtggtagagaatctccaggatgactttgatttcaactataaaaccctcaagagtcaaggaGACTACAAAGACGATGACGACAAGTAA |
| 21 | STAT3 variation 48 bp insertion | gaacagaaaatataaagtttctgaggagaattcaaatgaagccaaaacctcaaaaaagatacatgcaggacctgcaggcagtatccccaagagaaggctccctgttggccaggtgcagtggctcacgcctgtaatgccagcactttgagaacgaggaattctcttcttttttttctgcaggatTTGCACATTGCGTCAACTCATAAGATGTCTCAACGGCATGCGCAACTTctagaacagaaaatgaaagtggtagagaatctccaggatgactttgatttcaactataaaaccctcaagagtcaaggaGACTACAAAGACGATGACGACAAGTAA |
| 22 | STAT3 variation 96 bp insertion | gaacagaaaatataaagtttctgaggagaattcaaatgaagccaaaacctcaaaaaagatacatgcaggacctgcaggcagtatccccaagagaaggctccctgttggccaggtgcagtggctcacgcctgtaatgccagcactttgagaacgaggaattctcttcttttttttctgcaggatTTGCACATTGCGTCAACTCATAAGATGTCTCAACGGCATGCGCAACTTGTGAAGTGTCTACTATCCTTAAACGCATATCTCGCACAGTATCTCCCGctagaacagaaaatgaaagtggtagagaatctccaggatgactttgatttcaactataaaaccctcaagagtcaaggaGACTACAAAGACGATGACGACAAGTAA |
| 23 | STAT3 variation 12 bp deletion | gaacagaaaatataaagtttctgaggagaattcaaatgaagccaaaacctcaaaaaagatacatgcaggacctgcaggcagtatccccaagagaaggctccctgttggccaggtgcagtggctcacgcctgtaatgccagcactttgagaacgaggaattctcttcttttttttctgcagaaaatgaaagtggtagagaatctccaggatgactttgatttcaactataaaaccctcaagagtcaaggaGACTACAAAGACGATGACGACAAGTAA |
| 24 | PPIB variation substitute A to T | atgtggcttctcagggacattgcgttcagctgcactctgtatacctcaggggtgggaccagcacgtcactgagtgaaggaggggagggaggctctggcagttgtgcagccttcctggctgggctctgagggggctggaagaatttagaacaacgaggaattctcttcttttttttctgcaggTggtggtgcggaaggtggagagcaccaagacagacagccgggataaacccctgaaggatgtgatcatcgcagactgcggcaagatcgaggtggagaagccctttgccatcgccaaggagGACTACAAAGACGATGACGACAAGTAA |
| 25 | PPIB variation substitute A to C | atgtggcttctcagggacattgcgttcagctgcactctgtatacctcaggggtgggaccagcacgtcactgagtgaaggaggggagggaggctctggcagttgtgcagccttcctggctgggctctgagggggctggaagaatttagaacaacgaggaattctcttcttttttttctgcaggCggtggtgcggaaggtggagagcaccaagacagacagccgggataaacccctgaaggatgtgatcatcgcagactgcggcaagatcgaggtggagaagccctttgccatcgccaaggagGACTACAAAGACGATGACGACAAGTAA |
| 26 | PPIB variation substitute A to G | atgtggcttctcagggacattgcgttcagctgcactctgtatacctcaggggtgggaccagcacgtcactgagtgaaggaggggagggaggctctggcagttgtgcagccttcctggctgggctctgagggggctggaagaatttagaacaacgaggaattctcttcttttttttctgcaggGggtggtgcggaaggtggagagcaccaagacagacagccgggataaacccctgaaggatgtgatcatcgcagactgcggcaagatcgaggtggagaagccctttgccatcgccaaggagGACTACAAAGACGATGACGACAAGTAA |
| 27 | PPIB variation substitute T to A | atgtggcttctcagggacattgcgttcagctgcactctgtatacctcaggggtgggaccagcacgtcactgagtgaaggaggggagggaggctctggcagttgtgcagccttcctggctgggctctgagggggctggaagaatttagaacaacgaggaattctcttcttttttttctgcaggaggAggtgcggaaggtggagagcaccaagacagacagccgggataaacccctgaaggatgtgatcatcgcagactgcggcaagatcgaggtggagaagccctttgccatcgccaaggagGACTACAAAGACGATGACGACAAGTAA |
| 28 | PPIB variation substitute T to C | atgtggcttctcagggacattgcgttcagctgcactctgtatacctcaggggtgggaccagcacgtcactgagtgaaggaggggagggaggctctggcagttgtgcagccttcctggctgggctctgagggggctggaagaatttagaacaacgaggaattctcttcttttttttctgcaggaggCggtgcggaaggtggagagcaccaagacagacagccgggataaacccctgaaggatgtgatcatcgcagactgcggcaagatcgaggtggagaagccctttgccatcgccaaggagGACTACAAAGACGATGACGACAAGTAA |
| 29 | PPIB variation substitute T to G | atgtggcttctcagggacattgcgttcagctgcactctgtatacctcaggggtgggaccagcacgtcactgagtgaaggaggggagggaggctctggcagttgtgcagccttcctggctgggctctgagggggctggaagaatttagaacaacgaggaattctcttcttttttttctgcaggaggGggtgcggaaggtggagagcaccaagacagacagccgggataaacccctgaaggatgtgatcatcgcagactgcggcaagatcgaggtggagaagccctttgccatcgccaaggagGACTACAAAGACGATGACGACAAGTAA |
| 30 | PPIB variation substitute C to T | atgtggcttctcagggacattgcgttcagctgcactctgtatacctcaggggtgggaccagcacgtcactgagtgaaggaggggagggaggctctggcagttgtgcagccttcctggctgggctctgagggggctggaagaatttagaacaacgaggaattctcttcttttttttctgcaggaggtggtgTggaaggtggagagcaccaagacagacagccgggataaacccctgaaggatgtgatcatcgcagactgcggcaagatcgaggtggagaagccctttgccatcgccaaggagGACTACAAAGACGATGACGACAAGTAA |
| 31 | PPIB variation substitute C to G | atgtggcttctcagggacattgcgttcagctgcactctgtatacctcaggggtgggaccagcacgtcactgagtgaaggaggggagggaggctctggcagttgtgcagccttcctggctgggctctgagggggctggaagaatttagaacaacgaggaattctcttcttttttttctgcaggaggtggtgGggaaggtggagagcaccaagacagacagccgggataaacccctgaaggatgtgatcatcgcagactgcggcaagatcgaggtggagaagccctttgccatcgccaaggagGACTACAAAGACGATGACGACAAGTAA |
| 32 | PPIB variation substitute C to A | atgtggcttctcagggacattgcgttcagctgcactctgtatacctcaggggtgggaccagcacgtcactgagtgaaggaggggagggaggctctggcagttgtgcagccttcctggctgggctctgagggggctggaagaatttagaacaacgaggaattctcttcttttttttctgcaggaggtggtgAggaaggtggagagcaccaagacagacagccgggataaacccctgaaggatgtgatcatcgcagactgcggcaagatcgaggtggagaagccctttgccatcgccaaggagGACTACAAAGACGATGACGACAAGTAA |
| 33 | PPIB variation substitute G to A | atgtggcttctcagggacattgcgttcagctgcactctgtatacctcaggggtgggaccagcacgtcactgagtgaaggaggggagggaggctctggcagttgtgcagccttcctggctgggctctgagggggctggaagaatttagaacaacgaggaattctcttcttttttttctgcaggaAgtggtgcggaaggtggagagcaccaagacagacagccgggataaacccctgaaggatgtgatcatcgcagactgcggcaagatcgaggtggagaagccctttgccatcgccaaggagGACTACAAAGACGATGACGACAAGTAA |
| 34 | PPIB variation substitute G to T | atgtggcttctcagggacattgcgttcagctgcactctgtatacctcaggggtgggaccagcacgtcactgagtgaaggaggggagggaggctctggcagttgtgcagccttcctggctgggctctgagggggctggaagaatttagaacaacgaggaattctcttcttttttttctgcaggaTgtggtgcggaaggtggagagcaccaagacagacagccgggataaacccctgaaggatgtgatcatcgcagactgcggcaagatcgaggtggagaagccctttgccatcgccaaggagGACTACAAAGACGATGACGACAAGTAA |
| 35 | PPIB variation substitute G to C | atgtggcttctcagggacattgcgttcagctgcactctgtatacctcaggggtgggaccagcacgtcactgagtgaaggaggggagggaggctctggcagttgtgcagccttcctggctgggctctgagggggctggaagaatttagaacaacgaggaattctcttcttttttttctgcaggaCgtggtgcggaaggtggagagcaccaagacagacagccgggataaacccctgaaggatgtgatcatcgcagactgcggcaagatcgaggtggagaagccctttgccatcgccaaggagGACTACAAAGACGATGACGACAAGTAA |
| 36 | PPIB variation 1aa change | atgtggcttctcagggacattgcgttcagctgcactctgtatacctcaggggtgggaccagcacgtcactgagtgaaggaggggagggaggctctggcagttgtgcagccttcctggctgggctctgagggggctggaagaatttagaacaacgaggaattctcttcttttttttctgcagGCAgtggtgcggaaggtggagagcaccaagacagacagccgggataaacccctgaaggatgtgatcatcgcagactgcggcaagatcgaggtggagaagccctttgccatcgccaaggagGACTACAAAGACGATGACGACAAGTAA |
| 37 | PPIB variation 2aa change | atgtggcttctcagggacattgcgttcagctgcactctgtatacctcaggggtgggaccagcacgtcactgagtgaaggaggggagggaggctctggcagttgtgcagccttcctggctgggctctgagggggctggaagaatttagaacaacgaggaattctcttcttttttttctgcagGCACCGgtgcggaaggtggagagcaccaagacagacagccgggataaacccctgaaggatgtgatcatcgcagactgcggcaagatcgaggtggagaagccctttgccatcgccaaggagGACTACAAAGACGATGACGACAAGTAA |
| 38 | PPIB variation 3aa change | atgtggcttctcagggacattgcgttcagctgcactctgtatacctcaggggtgggaccagcacgtcactgagtgaaggaggggagggaggctctggcagttgtgcagccttcctggctgggctctgagggggctggaagaatttagaacaacgaggaattctcttcttttttttctgcagGCACCGACCcggaaggtggagagcaccaagacagacagccgggataaacccctgaaggatgtgatcatcgcagactgcggcaagatcgaggtggagaagccctttgccatcgccaaggagGACTACAAAGACGATGACGACAAGTAA |
| 39 | PPIB variation frameshift correction 1bp | atgtggcttctcagggacattgcgttcagctgcactctgtatacctcaggggtgggaccagcacgtcactgagtgaaggaggggagggaggctctggcagttgtgcagccttcctggctgggctctgagggggctggaagaatttagaacaacgaggaattctcttcttttttttctgcaggagAgtggtgcggaaggtggagagcaccaagacagacagccgggataaacccctgaaggatgtgatcatcgcagactgcggcaagatcgaggtggagaagccctttgccatcgccaaggagGACTACAAAGACGATGACGACAAGTAA |

| | | |
|---|---|---|
| 40 | PPIB variation frameshift correction 2bp | atgtggcttctcagggacattgcgttcagctgcactctgtatacctcaggggtgggaccagcacgtcactgagtgaaggaggggagggaggctctggcagttgtgcagccttcctggctgggctctgaggggggctggaagaatttagaacaacgaggaattctcttcttttttttctgcaggagAGgtggtgcggaaggtggagagcaccaagacagacagccgggataaacccctgaaggatgtgatcatcgcagactgcggcaagatcgaggtggagaagccctttgccatcgccaaggagGACTACAAAGACGATGACGACAAGTAA |
| 41 | PPIB variation 6 bp insertion | atgtggcttctcagggacattgcgttcagctgcactctgtatacctcaggggtgggaccagcacgtcactgagtgaaggaggggagggaggctctggcagttgtgcagccttcctggctgggctctgaggggggctggaagaatttagaacaacgaggaattctcttcttttttttctgcaggagTTGCACgtggtgcggaaggtggagagcaccaagacagacagccgggataaacccctgaaggatgtgatcatcgcagactgcggcaagatcgaggtggagaagccctttgccatcgccaaggagGACTACAAAGACGATGACGACAAGTAA |
| 42 | PPIB variation 12 bp insertion | atgtggcttctcagggacattgcgttcagctgcactctgtatacctcaggggtgggaccagcacgtcactgagtgaaggaggggagggaggctctggcagttgtgcagccttcctggctgggctctgaggggggctggaagaatttagaacaacgaggaattctcttcttttttttctgcaggagTTGCACATTGCGgtggtgcggaaggtggagagcaccaagacagacagccgggataaacccctgaaggatgtgatcatcgcagactgcggcaagatcgaggtggagaagccctttgccatcgccaaggagGACTACAAAGACGATGACGACAAGTAA |
| 43 | PPIB variation 24 bp insertion | atgtggcttctcagggacattgcgttcagctgcactctgtatacctcaggggtgggaccagcacgtcactgagtgaaggaggggagggaggctctggcagttgtgcagccttcctggctgggctctgaggggggctggaagaatttagaacaacgaggaattctcttcttttttttctgcaggagTTGCACATTGCGTCAACTCATAAGgtggtgcggaaggtggagagcaccaagacagacagccgggataaacccctgaaggatgtgatcatcgcagactgcggcaagatcgaggtggagaagccctttgccatcgccaaggagGACTACAAAGACGATGACGACAAGTAA |
| 44 | PPIB variation 48 bp insertion | atgtggcttctcagggacattgcgttcagctgcactctgtatacctcaggggtgggaccagcacgtcactgagtgaaggaggggagggaggctctggcagttgtgcagccttcctggctgggctctgaggggggctggaagaatttagaacaacgaggaattctcttcttttttttctgcaggagTTGCACATTGCGTCAACTCATAAGATGTCTCAACGGCATGCGCAACTTgtggtgcggaaggtggagagcaccaagacagacagccgggataaacccctgaaggatgtgatcatcgcagactgcggcaagatcgaggtggagaagccctttgccatcgccaaggagGACTACAAAGACGATGACGACAAGTAA |
| 45 | PPIB variation 96 bp insertion | atgtggcttctcagggacattgcgttcagctgcactctgtatacctcaggggtgggaccagcacgtcactgagtgaaggaggggagggaggctctggcagttgtgcagccttcctggctgggctctgaggggggctggaagaatttagaacaacgaggaattctcttcttttttttctgcaggagTTGCACATTGCGTCAACTCATAAGATGTCTCAACGGCATGCGCAACTTGTGAAGTGTCTACTATCCTTAAACGCATATCTCGCACAGTATCTCCCGgtggtgcggaaggtggagagcaccaagacagacagccgggataaacccctgaaggatgtgatcatcgcagactgcggcaagatcgaggtggagaagccctttgccatcgccaaggagGACTACAAAGACGATGACGACAAGTAA |
| 46 | PPIB variation 6 bp insertion | atgtggcttctcagggacattgcgttcagctgcactctgtatacctcaggggtgggaccagcacgtcactgagtgaaggaggggagggaggctctggcagttgtgcagccttcctggctgggctctgaggggggctggaagaatttagaacaacgaggaattctcttcttttttttctgcaggagcggaaggtggagagcaccaagacagacagccgggataaacccctgaaggatgtgatcatcgcagactgcggcaagatcgaggtggagaagccctttgccatcgccaaggagGACTACAAAGACGATGACGACAAGTAA |
| 47 | PPIB variation 12 bp deletion | atgtggcttctcagggacattgcgttcagctgcactctgtatacctcaggggtgggaccagcacgtcactgagtgaaggaggggagggaggctctggcagttgtgcagccttcctggctgggctctgaggggggctggaagaatttagaacaacgaggaattctcttcttttttttctgcaggaggtggagagcaccaagacagacagccgggataaacccctgaaggatgtgatcatcgcagactgcggcaagatcgaggtggagaagccctttgccatcgccaaggagGACTACAAAGACGATGACGACAAGTAA |
| 48 | HTT T cargo variation ISE BP | GCCACCATGGACTACAAAGACGATGACGACAAGggcccggctgtggctgaggagccCctCcaccgaccGTTGAAttgggcTGCATGacTGCATGgtTGCATGaacacaTATTAATttcctccacttagttctacacctcattcattcattcagtgagtgtttctcgactactatgaataaaccgttatactccatgttgcgggcagaatggggatctggacaggg |
| 49 | HTT T cargo variation BP | GCCACCATGGACTACAAAGACGATGACGACAAGggcccggctgtggctgaggagccCctCcaccgaccGTGAGTttgggcaacacaTATTAATttcctccacttagttctacacctcattcattcattcagtgagtgtttctcgactactatgaataaaccgttatactccatgttgcgggcagaatggggatctggacaggg |
| 50 | HTT T cargo variation ISE | GCCACCATGGACTACAAAGACGATGACGACAAGggcccggctgtggctgaggagccCctCcaccgaccGTTGAAttgggcTGCATGacTGCATGgtTGCATGaacacatccacttagttctacacctcattcattcattcagtgagtgtttctcgactactatgaataaaccgttatactccatgttgcgggcagaatggggatctggacaggg |
| 51 | HTT T cargo variation natGURAGU ISE BP | GCCACCATGGACTACAAAGACGATGACGACAAGggcccggctgtggctgaggagccCctCcaccgaccGTGAGTttgggcTGCATGacTGCATGgtTGCATGaacacaTATTAATttcctccacttagttctacacctcattcattcattcagtgagtgtttctcgactactatgaataaaccgttatactccatgttgcgggcagaatggggatctggacaggg |
| 52 | HTT T cargo variation GUAAGU BP | GCCACCATGGACTACAAAGACGATGACGACAAGggcccggctgtggctgaggagccCctCcaccgaccGTAAGTttgggcaacacaTATTAATttcctccacttagttctacacctcattcattcattcagtgagtgtttctcgactactatgaataaaccgttatactccatgttgcgggcagaatggggatctggacaggg |
| 53 | HTT T cargo variation GUAAGU | GCCACCATGGACTACAAAGACGATGACGACAAGggcccggctgtggctgaggagccCctCcaccgaccGTAAGTttgggcaacacatccacttagttctacacctcattcattcattcagtgagtgtttctcgactactatgaataaaccgttatactccatgttgcgggcagaatggggatctggacaggg |
| 54 | HTT T cargo variation 100 bp 3' shifted hybridization, GUAAGU ISE BP | GCCACCATGGACTACAAAGACGATGACGACAAGggcccggctgtggctgaggagccCctCcaccgaccGTAAGTttgggcTGCATGacTGCATGgtTGCATGaacacaTATTAATttcctccacttagttctacacctcattcattcattcagtgagtgtttctcgactactatgaataaaccgttatactccatgttgcgggcagaatggggatctggctggcggccgctcgagcatgcatctagagggccctattctatagtgtcacctaaatgctag |
| 55 | HTT T cargo variation 100 bp 3' shifted hybridization, GUAAGU ISE BP | GCCACCATGGACTACAAAGACGATGACGACAAGggcccggctgtggctgaggagccCctCcaccgaccGTAAGTttgggcTGCATGacTGCATGgtTGCATGaacacaTATTAATttccactatgaataaaccgttatactccatgttgcgggcagaatggggatctggacaggg |
| 56 | HTT T cargo variation U1 snRNA ISE BP | GCCACCATGGACTACAAAGACGATGACGACAAGggcccggctgtggctgaggagccCctCcaccgaccGTAAGTttgggcatacttacctggcaggggagataccatgatcacgaaggtggttttcccagggcgaggcttatccattgcactccggatgtgctgacccctgcgatttccccaaatgtgggaaactcgactgcataatttgtggtagtgggggactgcgttcgcgctttcccctgggtttcTGCATGacTGCATGgtTGCATGaacacaTATTAATttcctccacttagttctacacctcattcattcattcagtgagtgtttctcgactactatgaataaaccgttatactccatgttgcgggcagaatggggatctggacaggg |
| 57 | HTT T cargo variation U1 snRNA SL3 ISE BP | GCCACCATGGACTACAAAGACGATGACGACAAGggcccggctgtggctgaggagccCctCcaccgaccGTAAGTttgggctgcgatttccccaaatgtgggaaactcggggtttcTGCATGacTGCATGgtTGCATGaacacaTATTAATttcctccacttagttctacacctcattcattcattcagtgagtgtttctcgactactatgaataaaccgttatactccatgttgcgggcagaatggggatctggacaggg |
| 58 | HTT T cargo variation U1 snRNA smSL4 ISE BP | GCCACCATGGACTACAAAGACGATGACGACAAGggcccggctgtggctgaggagccCctCcaccgaccGTAAGTttgggcataatttgtggtagtgggggactgcgttcgcgctttcccctgggtttcTGCATGacTGCATGgtTGCATGaacacaTATTAATttcctccacttagttctacacctcattcattcattcagtgagtgtttctcgactactatgaataaaccgttatactccatgttgcgggcagaatggggatctggacaggg |
| 59 | HTT T cargo variation ISE U1 snRNA SL3 BP | GCCACCATGGACTACAAAGACGATGACGACAAGggcccggctgtggctgaggagccCctCcaccgaccGTAAGTttgggcTGCATGacTGCATGgtTGCATGtgcgatttccccaaatgtgggaaactcggggtttcaacacaTATTAATttcctccacttagttctacacctcattcattcattcagtgagtgtttctcgactactatgaataaaccgttatactccatgttgcgggcagaatggggatctggacaggg |
| 60 | HTT T cargo variation GUAAGU altISE BP | GCCACCATGGACTACAAAGACGATGACGACAAGggcccggctgtggctgaggagccCctCcaccgaccGTAAGTttgggcTTTGGGacTTTGGGgtTTTGGGaacacaTATTAATttcctccacttagttctacacctcattcattcattcagtgagtgtttctcgactactatgaataaaccgttatactccatgttgcgggcagaatggggatctggacaggg |

| | | |
|---|---|---|
| 61 | HTT T cargo variation GUAAGU mixISE BP | GCCACCATGGACTACAAAGACGATGACGACAAGggcccggctgtggctgaggagccCctCcaccgaccGTAAGTttgggcTTTGGGacGAGGGGgtTGCATGaacacaTATTAATttcctccacttagttctacacctcattcattcattcagtgagtgtttctcgactactatgaataaaccgttatactccatgttgcgggcagaatggggatctggacaggg |
| 62 | HTT T cargo variation GUAAGU mix-double ISE BP | GCCACCATGGACTACAAAGACGATGACGACAAGggcccggctgtggctgaggagccCctCcaccgaccGTAAGTttgggcTTTGGGcTTTGGGacGAGGGGcGAGGGGgtTGCATGcTGCATGaacacaTATTAATttcctccacttagttctacacctcattcattcattcagtgagtgtttctcgactactatgaataaaccgttatactccatgttgcgggcagaatggggatctggacaggg |
| 63 | HTT T cargo variation AX1 GUAAGU ISE BP | GCAAGGCGGAGGAAGGCCACCATGGACTACAAAGACGATGACGACAAGggcccggctgtggctgaggagccCctCcaccgaccGTAAGTttgggcTGCATGacTGCATGgtTGCATGaacacaTATTAATttcctccacttagttctacacctcattcattcattcagtgagtgtttctcgactactatgaataaaccgttatactccatgttgcgggcagaatggggatctggacaggg |
| 64 | HTT T cargo variation AX3 GUAAGU ISE BP | GCAAGGCGGAGGAAAGGCAAGGCGGAGGAAGGCAAGGCGGAGGAAGGCCACCATGGACTACAAAGACGATGACGACAAGggcccggctgtggctgaggagccCctCcaccgaccGTAAGTttgggcTGCATGacTGCATGgtTGCATGaacacaTATTAATttcctccacttagttctacacctcattcattcattcagtgagtgtttctcgactactatgaataaaccgttatactccatgttgcgggcagaatggggatctggacaggg |
| 65 | HTT T cargo variation BX4 GUAAGU ISE BP | GCACACAGGACCACACAGGACGCACACAGGACCACACAGGACGCCACCATGGACTACAAAGACGATGACGACAAGggcccggctgtggctgaggagccCctCcaccgaccGTAAGTttgggcTGCATGacTGCATGgtTGCATGaacacaTATTAATttcctccacttagttctacacctcattcattcattcagtgagtgtttctcgactactatgaataaaccgttatactccatgttgcgggcagaatggggatctggacaggg |
| 66 | HTT T cargo variation CX2 GUAAGU ISE BP | GAAAAAGAAAGAAAAAAAGAAAGAAGCCACCATGGACTACAAAGACGATGACGACAAGggcccggctgtggctgaggagccCctCcaccgaccGTAAGTttgggcTGCATGacTGCATGgtTGCATGaacacaTATTAATttcctccacttagttctacacctcattcattcattcagtgagtgtttctcgactactatgaataaaccgttatactccatgttgcgggcagaatggggatctggacaggg |
| 67 | HTT T cargo variation DX4 GUAAGU ISE BPHTT T cargo variation | GTCAGAGGATCAGAGGAGTCAGAGGATCAGAGGAGCCACCATGGACTACAAAGACGATGACGACAAGggcccggctgtggctgaggagccCctCcaccgaccGTAAGTttgggcTGCATGacTGCATGgtTGCATGaacacaTATTAATttcctccacttagttctacacctcattcattcattcagtgagtgtttctcgactactatgaataaaccgttatactccatgttgcgggcagaatggggatctggacaggg |
| 68 | HTT targeting cargo, original branchpoint | gcccggctgtggctgaggagccCctCcaccgaccGTAAGTttgggcTGCATGacTGCATGgtTGCATGaacacaTATTAATttcctccacttagttctacacctcattcattcattcagtgagtgtttctcgactactatgaataaaccgttatactccatgttgcgggcagaatggggatctggacagggaagcacagggcacgagttcaccaatggctgtcaagctacgctgc |
| 69 | USF1 NT cargo with XTEN 3x FLAG | TTGACCAAGTGGAGGGTGCTCTTCCAGCTCTTGAACAGGACCTAGAGAGTTGGATGTATTAGATGGGCGTACGCAgTATGTGCCCAGTTGTATGATTGTGCGTTTTCAAGGAAGGGAGTGTGCGTCGATTCGTTCAGTATCGACAgGGGGaacgacTtActctcttcttttttttctgcagagCaaGggAgggattctatccaaagcttgtgattatatccaggagcttcggcagagtaaccaccgcttgtctgaagaactgcagggacttgaccaactgcagctggacaatgacgtgcttcgacaacagGACTACAAGGACCACGACGGTGACTACAAGGACCACGACATCGACTACAAGGACGACGACGACAAGTAA |
| 70 | USF1 T cargo with XTEN 3x FLAG | agacccaagcttggtaccgagctcggatcctcacactttggacctcattttcatctaaggaaggtggtataatatctcccagggatacaggaacctcagggagagataagactactgtcatgtgtgcccctctctctaccatttctggaaacaataccaggaggcagaattcaggcatccaacgacTtActctcttcttttttttctgcagagCaaGggAgggattctatccaaagcttgtgattatatccaggagcttcggcagagtaaccaccgcttgtctgaagaactgcagggacttgaccaactgcagctggacaatgacgtgcttcgacaacagGACTACAAGGACCACGACGGTGACTACAAGGACCACGACATCGACTACAAGGACGACGACGACAAGTAA |
| 71 | RPL41 T cargo | taaatcaaacgtccacataaagaatgaggtggtaaaatgaacaagcactacggttctatcgttctctgttctgttaaatcctggctccagggagaaaacactcaaacgtttttctcctaaagatcttttaagatttccaaaccaaatgttaacgacTtActctcttcttttttttctgcaggctCaagcgcaaaagaagaaagatgaggcagaggtccaagGACTACAAAGACGATGACGACAAGTAA |
| 72 | STAT3 T cargo | ggaacagaaaatataaagtttctgaggagaattcaaatgaagccaaaacctcaaaaaagatacatgcaggacctgcaggcagtatccccaagagaaggctccctgttggccaggtgcagtggctcacgcctgtaatgccagcactttgagaacgaggaattctcttcttttttttctgcaggaCctCgaGcagaaaatgaaagtggtagagaatctccaggatgactttgatttcaactataaaccctcaagagtcaaggaGACTACAAAGACGATGACGACAAGTAA |
| 73 | PPIB T cargo | atgtggcttctcagggacattgcgttcagctgcactctgtatacctcaggggtgggaccagcacgtcactgagtgaaggaggggagggaggctctggcagttgtgcagccttcctggctgggctctgaggggggctggaagaatttagaacaacgaggaattctcttcttttttttctgcaggaAgtTgtTcggaaggtggagagcaccaagacagacagccgggataaacccctgaaggatgtgatcatcgcagactgcggcaagatcgaggtggagaagccctttgccatcgccaaggagGACTACAAAGACGATGACGACAAGTAA |
| 74 | SHANK3 T cargo | GAGGAGCCTAGAAGGCGCCGGGTTGGCAAGTGGGCAGGGAACAGAGACATGCTTTGTCGTGTTCTCAGCGGAGCTGGGGTACCTTGGTGGTCTCAGGCGTGAGACAGGGACTGCCTGAAGGCAGAGGCACCAAAAGCTACAAGAGCAAACaacgacTtActctcttcttttttttctgcagtcCCtgttCgaacgccagggcctcccaggcccagagaagctgccgggctccttgcggaaggggattccacggaccaagtctGACTACAAAGACGATGACGACAAGTAA |
| 75 | HTT targeting cargo, enhanced branchpoint | GCCACCATGGACTACAAAGACGATGACGACAAGggcccggctgtggctgaggagccCctCcaccgaccGTGAGTttgggcTGCATGacTGCATGgtTGCATGaacacaTATTAATttcctccacttagttctacacctcattcattcattcagtgagtgtttctcgactactatgaataaaccgttatactccatgttgcgggcagaatggggatctggacagggaagcacagggcacgagttcaccaatggctgtcaagctacgctgc |
| 76 | HTT targeting cargo, terminal hammerhead ribozyme | GCCACCATGGACTACAAAGACGATGACGACAAGggcccggctgtggctgaggagccCctCcaccgaccGTGAGTttgggcTGCATGacTGCATGgtTGCATGaacacaTATTAATttcctccacttagttctacacctcattcattcattcagtgagtgtttctcgactactatgaataaaccgttatactccatgttgcgggcagaatggggatctggacagggaagcacagggcacgagttcaccaatggctgtcaagctacgctgcctgatgagtccgtgaggacgaaacgagtaagctcgtcGCAGCG |
| 77 | HTT targeting cargo, terminal twister ribozyme | GCCACCATGGACTACAAAGACGATGACGACAAGggcccggctgtggctgaggagccCctCcaccgaccGTGAGTttgggcTGCATGacTGCATGgtTGCATGaacacaTATTAATttcctccacttagttctacacctcattcattcattcagtgagtgtttctcgactactatgaataaaccgttatactccatgttgcgggcagaatggggatctggacagggaagcacagggcacgagttcaccaatggctgtcaagctacgctgcCCGCCTAACACTGCCAATGCCGGTCCCAAGCCCGGATAAAAAGTGGAGGGGGCGG |
| 78 | HTT targeting cargo, terminal HDV ribozyme | GCCACCATGGACTACAAAGACGATGACGACAAGggcccggctgtggctgaggagccCctCcaccgaccGTGAGTttgggcTGCATGacTGCATGgtTGCATGaacacaTATTAATttcctccacttagttctacacctcattcattcattcagtgagtgtttctcgactactatgaataaaccgttatactccatgttgcgggcagaatggggatctggacagggaagcacagggcacgagttcaccaatggctgtcaagctacgctgcggccggcatggtcccagcctcctcgctggcgccggctgggcaacatgcttcggcatggcgaatgggacGCGGCCGC |
| 79 | PABPC1 5' targeting cargo 1 | GCCACCATGGACTACAAAGACGATGACGACAAGaaccccagtgcccccagctaccccatggcctcgctctacgtgggggacctccaccccgacgtgaccgaggcgatgctctacgagaagttcagcccggccgggcccatcctctccatccgggtctgcagggacatgatcacccgccgctccttgggctacgcgtatgtgaacttccagcagccAgcTgaTgGTGAGTttgggcTGCATGacTGCATGgtTGCATGaacacaTATTAATttccCAATATTACTTCAAAATTTTTGCTGGCTACTTAAGATTATATAAACTATGGTGACTGGAGTGGGAGGACACATGGTCTCACAGTTGAACGCTTCCTCTTTAAGCTTCAAGATGGCTAGACCTTTCAAGTATCACACACTAGTGTGGGACC |
| 80 | PABPC1 5' targeting cargo 2 | GCCACCATGGACTACAAAGACGATGACGACAAGaaccccagtgcccccagctaccccatggcctcgctctacgtgggggacctccaccccgacgtgaccgaggcgatgctctacgagaagttcagcccggccgggcccatcctctccatccgggtctgcagggacatgatcacccgccgctccttgggctacgcgtatgtgaacttccagcagccAgcTgaTgGTGAGTttgggcTGCATGacTGCATGgtTGCATGaacacaTATTAATttccGCTACTTAAGATTATATAAACTATGGTGACTGGAGTGGGAGGACACATGGTC |

| | | |
|---|---|---|
| | | TCACAGTTGAACGCTTCCTCTTTAAGCTTCAAGATGGCTAGACCTTTCAAGTATCACACACTAGTGTGGGACCTAAGTTGTATAAAGCAAAGACAAAT |
| 81 | PABPC1 5' targeting cargo 3 | GCCACCATGGACTACAAAGACGATGACGACAAGaaccccagtgcccccagctaccccatggcctcgctctacgtgggggacctccaccccgacgtgaccgaggcgatgctctacgagaagttcagcccggccgggcccatcctctccatccgggtctgcagggacatgatcacccgccgctccttgggctacgcgtatgtgaacttccagcagccAgcTgaTgGTGAGTttgggcTGCATGacTGCATGgtTGCATGaacacaTATTAATttccGTGACTGGAGTGGGAGGACACATGGTCTCACAGTTGAACGCTTCCTCTTTAAGCTTCAAGATGGCTAGACCTTTCAAGTATCACACACTAGTGTGGGACCTAAGTTGTATAAAGCAAAGACAAATACCAGAAGCCCCCGAAATTTCTCAC |
| 82 | PABPC1 5' targeting cargo 4 | GCCACCATGGACTACAAAGACGATGACGACAAGaaccccagtgcccccagctaccccatggcctcgctctacgtgggggacctccaccccgacgtgaccgaggcgatgctctacgagaagttcagcccggccgggcccatcctctccatccgggtctgcagggacatgatcacccgccgctccttgggctacgcgtatgtgaacttccagcagccAgcTgaTgGTGAGTttgggcTGCATGacTGCATGgtTGCATGaacacaTATTAATttccTCTCACAGTTGAACGCTTCCTCTTTAAGCTTCAAGATGGCTAGACCTTTCAAGTATCACACACTAGTGTGGGACCTAAGTTGTATAAAGCAAAGACAAATACCAGAAGCCCCCGAAATTTCTCACATATAAAAGAATTCCATATTGCTAA |
| 83 | PPIB 5' targeting cargo 1 | GCCACCATGGACTACAAAGACGATGACGACAAGctgcgcctctccgaacgcaacatgaaggtgctccttgccgccgccctcatcgcggggtccgtcttcttcctgctgctgccgggaccttctgcggccgatgagaagaagaaggggcccaaagtTacAgtGaagGTGAGTttgggcTGCATGacTGCATGgtTGCATGaacacaTATTAATttccCTTAGCTTCTTTAAAGGGGCGTTGCTAGGGGAGGGAAGGTACAAGAAGCTAACCTGAGGATGGGAGAGAGAATAGAGCCATATTTTTAGAGAAGTGGTTCTGAATCTGATTTTGGTGACGGTAAAAATCCTATGAGAATCTAATGAAAGC |
| 84 | PPIB 5' targeting cargo 2 | GCCACCATGGACTACAAAGACGATGACGACAAGctgcgcctctccgaacgcaacatgaaggtgctccttgccgccgccctcatcgcggggtccgtcttcttcctgctgctgccgggaccttctgcggccgatgagaagaagaaggggcccaaagtTacAgtGaagGTGAGTttgggcTGCATGacTGCATGgtTGCATGaacacaTATTAATttccTAGGGGAGGGAAGGTACAAGAAGCTAACCTGAGGATGGGAGAGAGAATAGAGCCATATTTTTAGAGAAGTGGTTCTGAATCTGATTTTGGTGACGGTAAAAATCCTATGAGAATCTAATGAAAGCTCCAGACCCTTTTCTTGGAAAACAT |
| 85 | PPIB 5' targeting cargo 3 | GCCACCATGGACTACAAAGACGATGACGACAAGctgcgcctctccgaacgcaacatgaaggtgctccttgccgccgccctcatcgcggggtccgtcttcttcctgctgctgccgggaccttctgcggccgatgagaagaagaaggggcccaaagtTacAgtGaagGTGAGTttgggcTGCATGacTGCATGgtTGCATGaacacaTATTAATttccAACCTGAGGATGGGAGAGAGAATAGAGCCATATTTTTAGAGAAGTGGTTCTGAATCTGATTTTGGTGACGGTAAAAATCCTATGAGAATCTAATGAAAGCTCCAGACCCTTTTCTTGGAAAACATGCTTATCTATACCCTCCTTGGACTA |
| 86 | PPIB 5' targeting cargo 4 | GCCACCATGGACTACAAAGACGATGACGACAAGctgcgcctctccgaacgcaacatgaaggtgctccttgccgccgccctcatcgcggggtccgtcttcttcctgctgctgccgggaccttctgcggccgatgagaagaagaaggggcccaaagtTacAgtGaagGTGAGTttgggcTGCATGacTGCATGgtTGCATGaacacaTATTAATttccAGCCATATTTTTAGAGAAGTGGTTCTGAATCTGATTTTGGTGACGGTAAAAATCCTATGAGAATCTAATGAAAGCTCCAGACCCTTTTCTTGGAAAACATGCTTATCTATACCCTCCTTGGACTACAGGATAACAATATTTGCTCTAAAC |
| 87 | RPL41 5' targeting cargo 1 | GCCACCATGGACTACAAAGACGATGACGACAAGagagccaagtggaggaagaagcgaatgcgGCgGTGAGTttgggcTGCATGacTGCATGgtTGCATGaacacaTATTAATttccCCAGAGTTGCCTTTCCCTCCCACATTAAATCAAACGTCCACATAAAGAATGAGGTGGTAAAATGAACAAGCACTACGGTTCTATCGTTCTCTGTTCTGTTAAATCCTGGCTCCAGGGAGAAAACACTCAAACGTTTTTCTCCTAAAGATC |
| 88 | RPL41 5' targeting cargo 2 | GCCACCATGGACTACAAAGACGATGACGACAAGagagccaagtggaggaagaagcgaatgcgGCgGTGAGTttgggcTGCATGacTGCATGgtTGCATGaacacaTATTAATttccTAAATCAAACGTCCACATAAAGAATGAGGTGGTAAAATGAACAAGCACTACGGTTCTATCGTTCTCTGTTCTGTTAAATCCTGGCTCCAGGGAGAAAACACTCAAACGTTTTTCTCCTAAAGATCTTTTAAGATTTCCAAACCAAATGTT |
| 89 | RPL41 5' targeting cargo 3 | GCCACCATGGACTACAAAGACGATGACGACAAGagagccaagtggaggaagaagcgaatgcgGCgGTGAGTttgggcTGCATGacTGCATGgtTGCATGaacacaTATTAATttccGAGGTGGTAAAATGAACAAGCACTACGGTTCTATCGTTCTCTGTTCTGTTAAATCCTGGCTCCAGGGAGAAAACACTCAAACGTTTTTCTCCTAAAGATCTTTTAAGATTTCCAAACCAAATGTTTCTCCTAAGTTTTGTCCAAGGAACT |
| 90 | RPL41 5' targeting cargo 4 | GCCACCATGGACTACAAAGACGATGACGACAAGagagccaagtggaggaagaagcgaatgcgGCgGTGAGTttgggcTGCATGacTGCATGgtTGCATGaacacaTATTAATttccCGGTTCTATCGTTCTCTGTTCTGTTAAATCCTGGCTCCAGGGAGAAAACACTCAAACGTTTTTCTCCTAAAGATCTTTTAAGATTTCCAAACCAAATGTTTCTCCTAAGTTTTGTCCAAGGAACTTCCCTCCTCCTGGGCTGGCAAAGTC |
| 91 | USF1 T cargo | tcacactttggacctcattttcatctaaggaaggtggtataatatctcccagggatacaggaacctcagggagagataagactactgtcatgtgtgcccctctctctaccatttctggaaacaataccaggaggcagaattcaggcatccaacgaggaattctcttcttttttttctgcagagCaaGggAgggattctatccaaagcttgtgattatatccaggagcttcggcagagtaaccaccgcttgtctgaagaactgcagggacttgaccaactgcagctggacaatgacgtgcttcgacaacagGACTACAAAGACGATGACGACAAGTAA |
| 92 | PPIB T cargo with 3x FLAG | atgtggcttctcagggacattgcgttcagctgcactctgtatacctcaggggtgggaccagcacgtcactgagtgaaggagggg agggaggctctggcagttgtgcagccttcctggctgggctctgaggggg ctggaagaatttagaacaacgacTtActctcttcttttttttctgcaggaAgtTgtTcggaaggtggagagcaccaagacagacagccgggataaacccctgaaggatgtgatcatcgcagactgcggcaagatcgaggtggagaagccctttgccatcgccaaggagGACTACAAGGACCACGACGGTGACTACAAGGACCACGACATCGACTACAAGGACGACGACGACAAG |
| 93 | PPIB T cargo with XTEN 3x FLAG | atgtggcttctcagggacattgcgttcagctgcactctgtatacctcaggggtgggaccagcacgtcactgagtgaaggaggggagggaggctctggcagttgtgcagccttcctggctgggctctgagggggctggaagaatttagaacaacgacTtActctcttcttttttttctgcaggaAgtTgtTcggaaggtggagagcaccaagacagacagccgggataaacccctgaaggatgtgatcatcgcagactgcggcaagatcgaggtggagaagccctttgccatcgccaaggagggagggccgagctctggcgcacccccaccaagtggagggtctcctgccgggtccccaacatctactgaagaaggcaccagcgaatccgcaacgcccgagtcaggccctggtacctccacagaaccatctgaaggtagtgcgcctggttccccagctggaagccctacttccaccgaagaaggcacgtcaaccgaaccaagtgaaggatctgcccctgggaccagcactgaaccatctgagGACTACAAGGACCACGACGGTGACTACAAGGACCACGACATCGACTACAAGGACGACGACGACAAGTAA |
| 94 | STAT3 3' targeting cargo, full length | ggaacagaaaatataaagtttctgaggagaattcaaatgaagccaaaacctcaaaaaagatacatgcaggacctgcaggcagtatccccaagagaaggctccctgttggccaggtgcagtggctcacgcctgtaatgccagcactttgagaacgatTaAttctcttcttttttttctgcaggatctagaacagaaaatgaaagtggtagagaatctccaggatgactttgatttcaactataaaaccctcaagagtcaaggagacatgcaagatctgaatggaaacaaccagtcagtgaccaggcagaagatgcagcagctggaacagatgctcactgcgctggaccagatgcggagaagcatcgtgagtgagctggcggggcttttgtcagcgatggagtacgtgcagaaaactctcacggacgaggagctggctgactggaagaggcggcaacagattgcctgcattggaggcccgcccaacatctgcctagatcggctagaaaactggataacgtcattagcagaatctcaacttcagacccgtcaacaaattaagaaactggaggagttgcagcaaaaagtttcctacaaaggggaccccattgtacagcaccggccgatgctggaggagagaatcgtggagctgtttagaaacttaatgaaaagtgcctttgtggtggagcggcagccctgcatgcccatgcatcctgaccggcccctcgtcatcaagaccggcgtccagttcactactaaagtcaggttgctggtcaaattccctgagttgaattatcagcttaaaattaaagtgtgcattgacaaagactctggggacgttgcagctctcagaggatcccggaaatttaacattctgggcacaaacacaaaagtgatgaacatggaagaatccaacaacggcagcctctctgcagaattcaaacacttgaccctgagggagcagagatgtgggaatgggggccgagccaattgtgatgcttccctgattgtgactgaggagctgcacctgatcacctttgagaccgaggtgtatcaccaaggcctcaagattgacctagagacccactccttgccagttgtggtgatctccaacatctgtcagatgccaaatgcctgggcgtccatcctgtggtacaacatgctgaccaacaatcccaagaatgtaaactttttttaccaagcccccaattggaacctgggatcaagtggccgaggtcctgagctggcagttctcctccaccaccaagcgaggactgagcatcgagcagctgactacactggcagagaaactcttgggacctggtgtgaattattcagggtgtcagatcacatgggctaaattttgcaaagaaaacatggctggcaagggcttctccttctgggtctggctggacaatatcattgaccttgtgaaaaagtacatcctggccctttggaacgaagggtacatcatgggctttatcagtaaggagcgggagcgggccatcttgagcactaagcctccaggcaccttcctgctaagattcagtgaaagcagcaaagaaggaggcgtcactttcacttgggtggagaaggacatcagcggtaagacccagatccagtccgtggaaccatacacaaagcagcagctgaacaacatgtcatttgctgaaatcatcatgggctataagatcatggatgctaccaatatcctggtgtctccactggtctatctctatcctgacattcccaaggaggaggcattcggaaagtattgtcggccagagagccaggagcatcctgaagctgacccaggcgctgccccatacctgaagaccaagtttatctgtgtgacaccaacgacctgcagcaataccattgacctgccgatgtcccccgcactttagattcattgatgcagtttggaaataatggtgaaggtgctgaaccctcagcaggagggcagtttgagtccctcacctttgacatggagttgacctcggagtgcgctacctcccccatgGACTACAAAGACGATGACGACAAGTAA |

| | | |
|---|---|---|
| 95 | PABPC1 T cargo | ttgagttctattacaccactattctagaattatgaatcgctcccctgcactactctttccttgtcctccccacactcgaaaaatatttctctttctccactagagaaagcagcagcagttgagagtatggctgttggagctgatgggattaacgaggaattctcttctttttttttctgcaggtCgaCgaGgctgtagctgtactacaagcccaccaagctaaagaggctgcccagaaagcagttaacagtgccaccggtgttccaactgttGACTACAAAGACGATGACGACAAGTAA |
| 96 | TOP2A T cargo | caatatttattgagcacttgctatgtgtcacgcacatggacataaagtctcaatcctcaaggagctcacagtccagtagaagtttgcaattaacacatattttgttaggtggtgggataaacaagagaagaaaaagtgggaaagtgactgaacgaggaattctcttctttttttttctgcagctcCttAgcAcgattgttatttccaccaaaagatgatcacacgttgaagtttttatatgatgacaaccagcgtgttgagcctgaatggtacattcctattattcccatggtgctgataaatggtgctgaaggaatcggtactgggtggtcctgcaaaGACTACAAAGACGATGACGACAAG |
| 97 | PPIB 3' T cargo, 50bp hybridization 1 | gaacgaggaattctcttctttttttttctgcaggaAgtTgtTcggaaggtggagagcaccaagacagacagccgggataaacccctgaaggatgtgatcatcgcagactgcggcaagatcgaggtggagaagccctttgccatcgccaaggagGACTACAAAGACGATGACGACAAGTAA |
| 98 | PPIB 3' T cargo, 50bp hybridization 2 | taatacgactcactataggggtgggaccagcacgtcactgagtgaaggaggggagggaggctctggcagaacgaggaattctcttctttttttttctgcaggaAgtTgtTcggaaggtggagagcaccaagacagacagccgggataaacccctgaaggatgtgatcatcgcagactgcggcaagatcgaggtggagaagccctttgccatcgccaaggagGACTACAAAGACGATGACGACAAGTAA |
| 99 | PPIB 3' T cargo, 50bp hybridization 3 | taatacgactcactataggttgtgcagccttcctggctgggctctgagggggctggaagaatttagaacaacgaggaattctcttctttttttttctgcaggaAgtTgtTcggaaggtggagagcaccaagacagacagccgggataaacccctgaaggatgtgatcatcgcagactgcggcaagatcgaggtggagaagccctttgccatcgccaaggagGACTACAAAGACGATGACGACAAGTAA |
| 100 | PPIB 3' T cargo, 100bp hybridization 1 | gggtgggaccagcacgtcactgagtgaaggaggggagggaggctctggcagaacgaggaattctcttctttttttttctgcaggaAgtTgtTcggaaggtggagagcaccaagacagacagccgggataaacccctgaaggatgtgatcatcgcagactgcggcaagatcgaggtggagaagccctttgccatcgccaaggagGACTACAAAGACGATGACGACAAGTAA |
| 101 | PPIB 3' T cargo, 100bp hybridization 2 | taatacgactcactataggggtgggaccagcacgtcactgagtgaaggaggggagggaggctctggcagttgtgcagccttcctggctgggctctgagggggctggaagaatttagaacaacgaggaattctcttctttttttttctgcaggaAgtTgtTcggaaggtggagagcaccaagacagacagccgggataaacccctgaaggatgtgatcatcgcagactgcggcaagatcgaggtggagaagccctttgccatcgccaaggagGACTACAAAGACGATGACGACAAGTAA |
| 102 | STAT3 3' T cargo, 200bp wider hybridization | acagaagtaaagaaagatttccttgggaacagaaaatataaagtttctgaggagaattcaaatgaagccaaaacctcaaaaaagatacatgcaggacctgcaggcagtatccccaagagaaggctccctgttggccaggtgcagtggctcacgcctgtaatgccagcactttgagaggctgagttgggaggatcacttgaaacgaggaattctcttctttttttttctgcaggaCctCgaGcagaaaatgaaagtggtagagaatctccaggatgactttgatttcaactataaaccctcaagagtcaaggaGACTACAAAGACGATGACGACAAGTAAGACTACAAAGACGATGACGACAAGTAA |
| 103 | STAT3 3' T cargo, 250bp wider hybridization | ctgtttaaaataagcaaacaaaaaaacagaagtaaagaaagatttccttgggaacagaaaatataaagtttctgaggagaattcaaatgaagccaaaacctcaaaaaagatacatgcaggacctgcaggcagtatccccaagagaaggctccctgttggccaggtgcagtggctcacgcctgtaatgccagcactttgagaggctgagttgggaggatcacttgagcccaggagttcatgatcagcctggaacgaggaattctcttctttttttttctgcaggaCctCgaGcagaaaatgaaagtggtagagaatctccaggatgactttgatttcaactataaaccctcaagagtcaaggaGACTACAAAGACGATGACGACAAGTAAGACTACAAAGACGATGACGACAAGTAA |
| 104 | STAT3 3' T cargo, 200bp additional 5' hybridization | ggaacagaaaatataaagtttctgaggagaattcaaatgaagccaaaacctcaaaaaagatacatgcaggacctgcaggcagtatccccaagagaaggctccctgttggccaggtgcagtggctcacgcctgtaatgccagcactttgagaggctgagttgggaggatcacttgagcccaggagttcatgatcagcctggaacgaggaattctcttctttttttttctgcaggaCctCgaGcagaaaatgaaagtggtagagaatctccaggatgactttgatttcaactataaaccctcaagagtcaaggaGACTACAAAGACGATGACGACAAGTAAGACTACAAAGACGATGACGACAAGTAA |
| 105 | STAT3 3' T cargo, 250bp additional 5' hybridization | ggaacagaaaatataaagtttctgaggagaattcaaatgaagccaaaacctcaaaaaagatacatgcaggacctgcaggcagtatccccaagagaaggctccctgttggccaggtgcagtggctcacgcctgtaatgccagcactttgagaggctgagttgggaggatcacttgagcccaggagttcatgatcagcctggacaacacagggagacccccatctctacaaattttttttttttaattagctaacgaggaattctcttctttttttttctgcaggaCctCgaGcagaaaatgaaagtggtagagaatctccaggatgactttgatttcaactataaaccctcaagagtcaaggaGACTACAAAGACGATGACGACAAGTAAGACTACAAAGACGATGACGACAAGTAA |
| 106 | STAT3 3' T cargo, 300bp additional 5' hybridization | ggaacagaaaatataaagtttctgaggagaattcaaatgaagccaaaacctcaaaaaagatacatgcaggacctgcaggcagtatccccaagagaaggctccctgttggccaggtgcagtggctcacgcctgtaatgccagcactttgagaggctgagttgggaggatcacttgagcccaggagttcatgatcagcctggacaacacagggagacccccatctctacaaattttttttttttaattagctgggcgtggtggtgcatgcctgtggtcccggctacttgggaggatgaggtaaacgaggaattctcttctttttttttctgcaggaCctCgaGcagaaaatgaaagtggtagagaatctccaggatgactttgatttcaactataaaccctcaagagtcaaggaGACTACAAAGACGATGACGACAAGTAAGACTACAAAGACGATGACGACAAGTAA |
| 107 | STAT3 3' T cargo, 200bp additional 3' hybridization | ctgtttaaaataagcaaacaaaaaaacagaagtaaagaaagatttccttgggaacagaaaatataaagtttctgaggagaattcaaatgaagccaaaacctcaaaaaagatacatgcaggacctgcaggcagtatccccaagagaaggctccctgttggccaggtgcagtggctcacgcctgtaatgccagcactttgagaacgaggaattctcttctttttttttctgcaggaCctCgaGcagaaaatgaaagtggtagagaatctccaggatgactttgatttcaactataaaccctcaagagtcaaggaGACTACAAAGACGATGACGACAAGTAAGACTACAAAGACGATGACGACAAGTAA |
| 108 | PPIB 3' T cargo, 300bp additional 5' hybridization | ATGTGGCTTCTCAGGGACATTGCGTTCAGCTGCACTCTGTATACCTCAGGGGTGGGACCAGCACGTCACTGAGTGAAGGAGGGGAGGGAGGCTCTGGCAGTTGTGCAGCCTTCCTGGCTGGGCTCTGAGGGGGCTGGAAGAATTTAGAACCTTGGAGGCATGGAGGTACAGGGTTTATTCTGGACAGGAGCACTGGGCTGCATCTGTGGGTTGGGTCCTTTTGGGAAAGGGATGGACACATGGAGCTCCTGCCCTGGGGTCTGTGTTGAATCCCCGGTGAGGATTGCCCAGTAGTAGCCCaacgaggaattctcttctttttttttctgcaggaAgtTgtTcggaaggtggagagcaccaagacagacagccgggataaacccctgaaggatgtgatcatcgcagactgcggcaagatcgaggtggagaagccctttgccatcgccaaggagGACTACAAAGACGATGACGACAAGTAA |
| 109 | PPIB 3' T cargo, 250bp additional 5' hybridization | ATGTGGCTTCTCAGGGACATTGCGTTCAGCTGCACTCTGTATACCTCAGGGGTGGGACCAGCACGTCACTGAGTGAAGGAGGGGAGGGAGGCTCTGGCAGTTGTGCAGCCTTCCTGGCTGGGCTCTGAGGGGGCTGGAAGAATTTAGAACCTTGGAGGCATGGAGGTACAGGGTTTATTCTGGACAGGAGCACTGGGCTGCATCTGTGGGTTGGGTCCTTTTGGGAAAGGGATGGACACATGGAGCTCCTaacgaggaattctcttctttttttttctgcaggaAgtTgtTcggaaggtggagagcaccaagacagacagccgggataaacccctgaaggatgtgatcatcgcagactgcggcaagatcgaggtggagaagccctttgccatcgccaaggagGACTACAAAGACGATGACGACAAGTAA |
| 110 | PPIB 3' T cargo, 200bp additional 5' hybridization | ATGTGGCTTCTCAGGGACATTGCGTTCAGCTGCACTCTGTATACCTCAGGGGTGGGACCAGCACGTCACTGAGTGAAGGAGGGGAGGGAGGCTCTGGCAGTTGTGCAGCCTTCCTGGCTGGGCTCTGAGGGGGCTGGAAGAATTTAGAACCTTGGAGGCATGGAGGTACAGGGTTTATTCTGGACAGGAGCACTGGGCTGaacgaggaattctcttctttttttttctgcaggaAgtTgtTcggaaggtggagagcaccaagacagacagccgggataaacccctgaaggatgtgatcatcgcagactgcggcaagatcgaggtggagaagccctttgccatcgccaaggagGACTACAAAGACGATGACGACAAGTAA |
| 111 | PPIB 3' T cargo, 300bp additional 3' hybridization | CACACAAAACTGGAGGCACCAAAATTCTAACAGACTCCTGGCCAGAGCAGGGAGAATGCAGATTTGACGAGGGGGTACAGGAATTTTGTTCCTTTGAAGTAAGACCCAGGTTGGGCCAAGGGTGAGGAGGAGGAAGAGGGTGACCAGGGCATGTGGCTTCTCAGGGACATTGCGTTCAGCTGCACTCTGTATACCTCAGGGGTGGGACCAGCACGTCACTGAGTGAAGGAGGGGAGGGAGGCTCTGGCAGTTGTGCAGCCTTCCTGGCTGGGCTCTGAGGGGGCTGGAAGAATTTAGAACaacgaggaattctcttctttttttttctgcaggaAgtTgtTcggaaggtggagagcaccaagacagacagccgggataaacccctgaaggatgtgatcatcgcagactgcggcaagatcgaggtggagaagccctttgccatcgccaaggagGACTACAAAGACGATGACGACAAGTAA |
| 112 | PPIB 3' T cargo, 250bp additional 3' hybridization | GGAGAATGCAGATTTGACGAGGGGGTACAGGAATTTTGTTCCTTTGAAGTAAGACCCAGGTTGGGCCAAGGGTGAGGAGGAGGAAGAGGGTGACCAGGGCATGTGGCTTCTCAGGGACATTGCGTTCAGCTGCACTCTGTATACCTCAGGGGTGGGACCAGCACGTCACTGAGTGAAGGAGGGGAGGGAGGCTCTGGCAGTTGTGCAGCCTTCCTGGCTGGGCTCTGAGGGGGCTGGAAGAATTTAGAACaacgaggaattctcttctttttttttctgcaggaAgtTgtTcggaaggtggagagcaccaagacagacagccgggataaacccctgaaggatgtgatcatcgcagactgcggcaagatcgaggtggagaagccctttgccatcgccaaggagGACTACAAAGACGATGACGACAAGTAA |

| | | |
|---|---|---|
| 113 | PPIB 3' T cargo, 250bp additional 3' hybridization | AAGACCCAGGTTGGGCCAAGGGTGAGGAGGAGGAAGAGGGTGACCAGGGCATGTGGCTTCTCAGGGACATTGCGTTCAGCTGCACTCTGTATACCTCAGGGGTGGGACCAGCACGTCACTGAGTGAAGGAGGGGAGGGAGGCTCTGGCAGTTGTGCAGCCTTCCTGGCTGGGCTCTGAGGGGGCTGGAAGAATTTAGAACaacgaggaattctcttcttttttttctgcaggaAgtTgtTcggaaggtggagagcaccaagacagacagccgggataaacccctgaaggatgtgatcatcgcagactgcggcaagatcgaggtggagaagccctttgccatcgccaaggagGACTACAAAGACGATGACGACAAGTAA |
| 114 | PPIB 3' T cargo, 300bp wider hybridization | TACAGGAATTTTGTTCCTTTGAAGTAAGACCCAGGTTGGGCCAAGGGTGAGGAGGAGGAAGAGGGTGACCAGGGCATGTGGCTTCTCAGGGACATTGCGTTCAGCTGCACTCTGTATACCTCAGGGGTGGGACCAGCACGTCACTGAGTGAAGGAGGGGAGGGAGGCTCTGGCAGTTGTGCAGCCTTCCTGGCTGGGCTCTGAGGGGGCTGGAAGAATTTAGAACCTTGGAGGCATGGAGGTACAGGGTTTATTCTGGACAGGAGCACTGGGCTGCATCTGTGGGTTGGGTCCTTTTGGGaacgaggaattctcttcttttttttctgcaggaAgtTgtTcggaaggtggagagcaccaagacagacagccgggataaacccctgaaggatgtgatcatcgcagactgcggcaagatcgaggtggagaagccctttgccatcgccaaggagGACTACAAAGACGATGACGACAAGTAA |
| 115 | PPIB 3' T cargo, 250bp wider hybridization | AAGACCCAGGTTGGGCCAAGGGTGAGGAGGAGGAAGAGGGTGACCAGGGCATGTGGCTTCTCAGGGACATTGCGTTCAGCTGCACTCTGTATACCTCAGGGGTGGGACCAGCACGTCACTGAGTGAAGGAGGGGAGGGAGGCTCTGGCAGTTGTGCAGCCTTCCTGGCTGGGCTCTGAGGGGGCTGGAAGAATTTAGAACCTTGGAGGCATGGAGGTACAGGGTTTATTCTGGACAGGAGCACTGGGCTGaacgaggaattctcttcttttttttctgcaggaAgtTgtTcggaaggtggagagcaccaagacagacagccgggataaacccctgaaggatgtgatcatcgcagactgcggcaagatcgaggtggagaagccctttgccatcgccaaggagGACTACAAAGACGATGACGACAAGTAA |
| 116 | PPIB 3' T cargo, 200bp wider hybridization | GGAGGAGGAAGAGGGTGACCAGGGCATGTGGCTTCTCAGGGACATTGCGTTCAGCTGCACTCTGTATACCTCAGGGGTGGGACCAGCACGTCACTGAGTGAAGGAGGGGAGGGAGGCTCTGGCAGTTGTGCAGCCTTCCTGGCTGGGCTCTGAGGGGGCTGGAAGAATTTAGAACCTTGGAGGCATGGAGGTACAGGGTTaacgaggaattctcttcttttttttctgcaggaAgtTgtTcggaaggtggagagcaccaagacagacagccgggataaacccctgaaggatgtgatcatcgcagactgcggcaagatcgaggtggagaagccctttgccatcgccaaggagGACTACAAAGACGATGACGACAAGTAA |
| 117 | STAT3 ESE AX1 | ggaacagaaaatataaagtttctgaggagaattcaaatgaagccaaaacctcaaaaaagatacatgcaggacctgcaggcagtatccccaagagaaggctccctgttggccaggtgcagtggctcacgcctgtaatgccagcactttgagaacgaggaattctcttcttttttttctgcaggaCctCgaGcagaaaatgaaagtggtagagaatctccaggatgactttgatttcaactataaaccctcaagagtcaaggaGACTACAAAGACGATGACGACAAGTAAGCAAGGCGGAGGAAG |
| 118 | STAT3 ESE AX2 | ggaacagaaaatataaagtttctgaggagaattcaaatgaagccaaaacctcaaaaaagatacatgcaggacctgcaggcagtatccccaagagaaggctccctgttggccaggtgcagtggctcacgcctgtaatgccagcactttgagaacgaggaattctcttcttttttttctgcaggaCctCgaGcagaaaatgaaagtggtagagaatctccaggatgactttgatttcaactataaaccctcaagagtcaaggaGACTACAAAGACGATGACGACAAGTAAGCAAGGCGGAGGAAGCAAGGCGGAGGAAG |
| 119 | STAT3 ESE AX3 | ggaacagaaaatataaagtttctgaggagaattcaaatgaagccaaaacctcaaaaaagatacatgcaggacctgcaggcagtatccccaagagaaggctccctgttggccaggtgcagtggctcacgcctgtaatgccagcactttgagaacgaggaattctcttcttttttttctgcaggaCctCgaGcagaaaatgaaagtggtagagaatctccaggatgactttgatttcaactataaaccctcaagagtcaaggaGACTACAAAGACGATGACGACAAGTAAGCAAGGCGGAGGAAAGCAAGGCGGAGGAAGGCAAGGCGGAGGAAG |
| 120 | STAT3 ESE AX4 | ggaacagaaaatataaagtttctgaggagaattcaaatgaagccaaaacctcaaaaaagatacatgcaggacctgcaggcagtatccccaagagaaggctccctgttggccaggtgcagtggctcacgcctgtaatgccagcactttgagaacgaggaattctcttcttttttttctgcaggaCctCgaGcagaaaatgaaagtggtagagaatctccaggatgactttgatttcaactataaaccctcaagagtcaaggaGACTACAAAGACGATGACGACAAGTAAGCAAGGCGGAGGAAAGCAAGGCGGAGGAAGAGCAAGGCGGAGGAAGGCAAGGCGGAGGAAG |
| 121 | STAT3 ESE BX2 | ggaacagaaaatataaagtttctgaggagaattcaaatgaagccaaaacctcaaaaaagatacatgcaggacctgcaggcagtatccccaagagaaggctccctgttggccaggtgcagtggctcacgcctgtaatgccagcactttgagaacgaggaattctcttcttttttttctgcaggaCctCgaGcagaaaatgaaagtggtagagaatctccaggatgactttgatttcaactataaaccctcaagagtcaaggaGACTACAAAGACGATGACGACAAGTAAGCACACAGGACCACACAGGAC |
| 122 | STAT3 ESE CX2 | ggaacagaaaatataaagtttctgaggagaattcaaatgaagccaaaacctcaaaaaagatacatgcaggacctgcaggcagtatccccaagagaaggctccctgttggccaggtgcagtggctcacgcctgtaatgccagcactttgagaacgaggaattctcttcttttttttctgcaggaCctCgaGcagaaaatgaaagtggtagagaatctccaggatgactttgatttcaactataaaccctcaagagtcaaggaGACTACAAAGACGATGACGACAAGTAAGAAAAAGAAAGAAAAAAAGAAAGAA |
| 123 | STAT3 ESE DX2 | ggaacagaaaatataaagtttctgaggagaattcaaatgaagccaaaacctcaaaaaagatacatgcaggacctgcaggcagtatccccaagagaaggctccctgttggccaggtgcagtggctcacgcctgtaatgccagcactttgagaacgaggaattctcttcttttttttctgcaggaCctCgaGcagaaaatgaaagtggtagagaatctccaggatgactttgatttcaactataaaccctcaagagtcaaggaGACTACAAAGACGATGACGACAAGTAAGTCAGAGGATCAGAGGA |
| 124 | SHANK3 branchpoint v0 | gaggagcctagaaggcgccgggttggcaagtgggcagggaacagagacatgctttgtcgtgttctcagcggagctggggtaccttggtggtctcaggcgtgagacagggactgcctgaaggcagaggcaccaaaagctacaagagcaaacaacgaggaAttctcttcttttttttctgcagtcCCtgttCgaacgccagggcctcccaggcccagagaagctgccgggctccttgcggaaggggattccacggaccaagtctGACTACAAAGACGATGACGACAAGTAA |
| 125 | SHANK3 branchpoint v1 | gaggagcctagaaggcgccgggttggcaagtgggcagggaacagagacatgctttgtcgtgttctcagcggagctggggtaccttggtggtctcaggcgtgagacagggactgcctgaaggcagaggcaccaaaagctacaagagcaaacaacgacTaActctcttcttttttttctgcagtcCCtgttCgaacgccagggcctcccaggcccagagaagctgccgggctccttgcggaaggggattccacggaccaagtctGACTACAAAGACGATGACGACAAGTAA |
| 126 | SHANK3 branchpoint v3 | gaggagcctagaaggcgccgggttggcaagtgggcagggaacagagacatgctttgtcgtgttctcagcggagctggggtaccttggtggtctcaggcgtgagacagggactgcctgaaggcagaggcaccaaaagctacaagagcaaacaacgatTaAttctcttcttttttttctgcagtcCCtgttCgaacgccagggcctcccaggcccagagaagctgccgggctccttgcggaaggggattccacggaccaagtctGACTACAAAGACGATGACGACAAGTAA |
| 127 | SHANK3 branchpoint v6 | gaggagcctagaaggcgccgggttggcaagtgggcagggaacagagacatgctttgtcgtgttctcagcggagctggggtaccttggtggtctcaggcgtgagacagggactgcctgaaggcagaggcaccaaaagctacaagagcaaacaacgatTgActctcttcttttttttctgcagtcCCtgttCgaacgccagggcctcccaggcccagagaagctgccgggctccttgcggaaggggattccacggaccaagtctGACTACAAAGACGATGACGACAAGTAA |
| 128 | SHANK3 branchpoint v12 | gaggagcctagaaggcgccgggttggcaagtgggcagggaacagagacatgctttgtcgtgttctcagcggagctggggtaccttggtggtctcaggcgtgagacagggactgcctgaaggcagaggcaccaaaagctacaagagcaaacaacgacTcAttctcttcttttttttctgcagtcCCtgttCgaacgccagggcctcccaggcccagagaagctgccgggctccttgcggaaggggattccacggaccaagtctGACTACAAAGACGATGACGACAAGTAA |
| 129 | SHANK3 branchpoint v13 | gaggagcctagaaggcgccgggttggcaagtgggcagggaacagagacatgctttgtcgtgttctcagcggagctggggtaccttggtggtctcaggcgtgagacagggactgcctgaaggcagaggcaccaaaagctacaagagcaaacaacgacTtActctcttcttttttttctgcagtcCCtgttCgaacgccagggcctcccaggcccagagaagctgccgggctccttgcggaaggggattccacggaccaagtctGACTACAAAGACGATGACGACAAGTAA |
| 130 | SHANK3 branchpoint v15 | gaggagcctagaaggcgccgggttggcaagtgggcagggaacagagacatgctttgtcgtgttctcagcggagctggggtaccttggtggtctcaggcgtgagacagggactgcctgaaggcagaggcaccaaaagctacaagagcaaacaacgatTtAttctcttcttttttttctgcagtcCCtgttCgaacgccagggcctcccaggcccagagaagctgccgggctccttgcggaaggggattccacggaccaagtctGACTACAAAGACGATGACGACAAGTAA |
| 131 | SHANK3 branchpoint yeast | gaggagcctagaaggcgccgggttggcaagtgggcagggaacagagacatgctttgtcgtgttctcagcggagctggggtaccttggtggtctcaggcgtgagacagggactgcctgaaggcagaggcac |

| | | |
|---|---|---|
| | | caaaagctacaagagcaaacaacTACTAACtctcttcttttttttctgcagtcCCtgttCgaacgccagggcctcccaggcccagagaagctgccgggctccttgcggaaggggattccacggaccaag<br>tctGACTACAAAGACGATGACGACAAGTAA |
| 132 | SHANK3 ESE AX1 | GAGGAGCCTAGAAGGCGCCGGGTTGGCAAGTGGGCAGGGAACAGAGACATGCTTTGTCGTGTTCTCAGCGGAGCTGGGGTAC<br>CTTGGTGGTCTCAGGCGTGAGACAGGGACTGCCTGAAGGCAGAGGCACCAAAAGCTACAAGAGCAAACaacgaggaattctcttcttttttttct<br>gcagtcCCtgttCgaacgccagggcctcccaggcccagagaagctgccgggctccttgcggaaggggattccacggaccaagtctGACTACAAAGACGATGACGACAAGTAA<br>GCAAGGCGGAGGAAG |
| 133 | SHANK3 ESE AX2 | GAGGAGCCTAGAAGGCGCCGGGTTGGCAAGTGGGCAGGGAACAGAGACATGCTTTGTCGTGTTCTCAGCGGAGCTGGGGTAC<br>CTTGGTGGTCTCAGGCGTGAGACAGGGACTGCCTGAAGGCAGAGGCACCAAAAGCTACAAGAGCAAACaacgaggaattctcttcttttttttct<br>gcagtcCCtgttCgaacgccagggcctcccaggcccagagaagctgccgggctccttgcggaaggggattccacggaccaagtctGACTACAAAGACGATGACGACAAGTAA<br>GCAAGGCGGAGGAAGCAAGGCGGAGGAAG |
| 134 | SHANK3 ESE AX3 | GAGGAGCCTAGAAGGCGCCGGGTTGGCAAGTGGGCAGGGAACAGAGACATGCTTTGTCGTGTTCTCAGCGGAGCTGGGGTAC<br>CTTGGTGGTCTCAGGCGTGAGACAGGGACTGCCTGAAGGCAGAGGCACCAAAAGCTACAAGAGCAAACaacgaggaattctcttcttttttttct<br>gcagtcCCtgttCgaacgccagggcctcccaggcccagagaagctgccgggctccttgcggaaggggattccacggaccaagtctGACTACAAAGACGATGACGACAAGTAA<br>GCAAGGCGGAGGAAAGCAAGGCGGAGGAAGGCAAGGCGGAGGAAG |
| 135 | SHANK3 ESE AX4 | GAGGAGCCTAGAAGGCGCCGGGTTGGCAAGTGGGCAGGGAACAGAGACATGCTTTGTCGTGTTCTCAGCGGAGCTGGGGTAC<br>CTTGGTGGTCTCAGGCGTGAGACAGGGACTGCCTGAAGGCAGAGGCACCAAAAGCTACAAGAGCAAACaacgaggaattctcttcttttttttct<br>gcagtcCCtgttCgaacgccagggcctcccaggcccagagaagctgccgggctccttgcggaaggggattccacggaccaagtctGACTACAAAGACGATGACGACAAGTAA<br>GCAAGGCGGAGGAAAGCAAGGCGGAGGAAGAGCAAGGCGGAGGAAGGCAAGGCGGAGGAAG |
| 136 | SHANK3 ESE BX2 | GAGGAGCCTAGAAGGCGCCGGGTTGGCAAGTGGGCAGGGAACAGAGACATGCTTTGTCGTGTTCTCAGCGGAGCTGGGGTAC<br>CTTGGTGGTCTCAGGCGTGAGACAGGGACTGCCTGAAGGCAGAGGCACCAAAAGCTACAAGAGCAAACaacgaggaattctcttcttttttttct<br>gcagtcCCtgttCgaacgccagggcctcccaggcccagagaagctgccgggctccttgcggaaggggattccacggaccaagtctGACTACAAAGACGATGACGACAAGTAA<br>GCACACAGGACCACACAGGAC |
| 137 | SHANK3 ESE BX4 | GAGGAGCCTAGAAGGCGCCGGGTTGGCAAGTGGGCAGGGAACAGAGACATGCTTTGTCGTGTTCTCAGCGGAGCTGGGGTAC<br>CTTGGTGGTCTCAGGCGTGAGACAGGGACTGCCTGAAGGCAGAGGCACCAAAAGCTACAAGAGCAAACaacgaggaattctcttcttttttttct<br>gcagtcCCtgttCgaacgccagggcctcccaggcccagagaagctgccgggctccttgcggaaggggattccacggaccaagtctGACTACAAAGACGATGACGACAAGTAA<br>GCACACAGGACCACACAGGACGCACACAGGACCACACAGGAC |
| 138 | SHANK3 ESE CX2 | GAGGAGCCTAGAAGGCGCCGGGTTGGCAAGTGGGCAGGGAACAGAGACATGCTTTGTCGTGTTCTCAGCGGAGCTGGGGTAC<br>CTTGGTGGTCTCAGGCGTGAGACAGGGACTGCCTGAAGGCAGAGGCACCAAAAGCTACAAGAGCAAACaacgaggaattctcttcttttttttct<br>gcagtcCCtgttCgaacgccagggcctcccaggcccagagaagctgccgggctccttgcggaaggggattccacggaccaagtctGACTACAAAGACGATGACGACAAGTAA<br>GAAAAAGAAAGAAAAAAAGAAAGAA |
| 139 | SHANK3 ESE CX4 | GAGGAGCCTAGAAGGCGCCGGGTTGGCAAGTGGGCAGGGAACAGAGACATGCTTTGTCGTGTTCTCAGCGGAGCTGGGGTAC<br>CTTGGTGGTCTCAGGCGTGAGACAGGGACTGCCTGAAGGCAGAGGCACCAAAAGCTACAAGAGCAAACaacgaggaattctcttcttttttttct<br>gcagtcCCtgttCgaacgccagggcctcccaggcccagagaagctgccgggctccttgcggaaggggattccacggaccaagtctGACTACAAAGACGATGACGACAAGTAA<br>GAAAAAGAAAGAAAAAAAGAAAGAA |
| 140 | SHANK3 ESE DX2 | GAGGAGCCTAGAAGGCGCCGGGTTGGCAAGTGGGCAGGGAACAGAGACATGCTTTGTCGTGTTCTCAGCGGAGCTGGGGTAC<br>CTTGGTGGTCTCAGGCGTGAGACAGGGACTGCCTGAAGGCAGAGGCACCAAAAGCTACAAGAGCAAACaacgaggaattctcttcttttttttct<br>gcagtcCCtgttCgaacgccagggcctcccaggcccagagaagctgccgggctccttgcggaaggggattccacggaccaagtctGACTACAAAGACGATGACGACAAGTAA<br>GTCAGAGGATCAGAGGA |
| 141 | SHANK3 ESE DX4 | GAGGAGCCTAGAAGGCGCCGGGTTGGCAAGTGGGCAGGGAACAGAGACATGCTTTGTCGTGTTCTCAGCGGAGCTGGGGTAC<br>CTTGGTGGTCTCAGGCGTGAGACAGGGACTGCCTGAAGGCAGAGGCACCAAAAGCTACAAGAGCAAACaacgaggaattctcttcttttttttct<br>gcagtcCCtgttCgaacgccagggcctcccaggcccagagaagctgccgggctccttgcggaaggggattccacggaccaagtctGACTACAAAGACGATGACGACAAGTAA<br>GTCAGAGGATCAGAGGAGTCAGAGGATCAGAGGA |

### Supplementary Table 3: Nuclease sequences.

| | | |
|---|---|---|
| 1 | wildtype *Dis*Cas7-11 | MKRTADGSEFESPKKKRKVTTTMKISIEFLEPFRMTKWQESTRRNKNNKEFVRGQAFARWHRNKKDNTKGRPYITGTLLRSAVIRS<br>AENLLTLSDGKISEKTCCPGKFDTEDKDRLLQLRQRSTLRWTDKNPCPDNAETYCPFCELLGRSGNDGKKAEKKDWRFRIHFGNLS<br>LPGKPDFDGPKAIGSQRVLNRVDFKSGKAHDFFKAYEVDHTRFPRFEGEITIDNKVSAEARKLLCDSLKFTDRLCGALCVIRFDEYTP<br>AADSGKQTENVQAEPNANLAEKTAEQIISILDDNKKTEYTRLLADAIRSLRRSSKLVAGLPKDHDGKDDHYLWDIGKKKKDENSVT<br>IRQILTTSADTKELKNAGKWREFCEKLGEALYLKSKDMSGGLKITRRILGDAEFHGKPDRLEKSRSVSIGSVLKETVVCGELVAKTPF<br>FFGAIDEDAKQTDLQVLLTPDNKYRLPRSAVRGILRRDLQTYFDSPCNAELGGRPCMCKTCRIMRGITVMDARSEYNAPPEIRHRTRI<br>NPFTGTVAEGALFNMEVAPEGIVFPFQLRYRGSEDGLPDALKTVLKWWAEGQAFMSGAASTGKGRFRMENAKYETLDLSDENQR<br>NDYLKNWGWRDEKGLEELKKRLNSGLPEPGNYRDPKWHEINVSIEMASPFINGDPIRAAVDKRGTDVVTFVKYKAEGEEAKPVCA<br>YKAESFRGVIRSAVARIHMEDGVPLTELTHSDCECLLCQIFGSEYEAGKIRFEDLVFESDPEPVTFDHVAIDRFTGGAADKKKFDDSP<br>LPGSPARPLMLKGSFWIRRDVLEDEEYCKALGKALADVNNGLYPLGGKSAIGYGQVKSLGIKGDDKRISRLMNPAFDETDVAVPEK<br>PKTDAEVRIEAEKVYYPHYFVEPHKKVEREEKPCGHQKFHEGRLTGKIRCKLITKTPLIVPDTSNDDFFRPADKEARKEKDEYHKSY<br>AFFRLHKQIMIPGSELRGMVSSVYETVTNSCFRIFDETKRLSWRMDADHQNVLQDFLPGRVTADGKHIQKFSETARVPFYDKTQKH<br>FDILDEQEIAGEKPVRMWVKRFIKRLSLVDPAKHPQKKQDNKWKRRKEGIATFIEQKNGSYYFNVVTNNGCTSFHLWHKPDNFDQ<br>EKLEGIQNGEKLDCWVRDSRYQKAFQEIPENDPDGWECKEGYLHVVGPSKVEFSDKKGDVINNFQGTLPSVPNDWKTIRTNDFKN<br>RKRKNEPVFCCEDDKGNYYTMAKYCETFFFDLKENEEYEIPEKARIKYKELLRVYNNNPQAVPESVFQSRVARENVEKLKSGDLVY<br>FKHNEKYVEDIVPVRISRTVDDRMIGKRMSADLRPCHGDWVEDGDLSALNAYPEKRLLLRHPKGLCPACRLFGTGSYKGRVRFGF<br>ASLENDPEWLIPGKNPGDPFHGGPVMLSLLERPRPTWSIPGSDNKFKVPGRKFYVHHHAWKTIKDGNHPTTGKAIEQSPNNRTVEAL<br>AGGNSFSFEIAFENLKEWELGLLIHSLQLEKGLAHKLGMAKSMGFGSVEIDVESVRLRKDWKQWRNGNSEIPNWLGKGFAKLKEW<br>FRDELDFIENLKKLLWFPEGDQAPRVCYPMLRKKDDPNGNSGYEELKDGEFKKEDRQKKLTTPWTPWAKRTADGSEFESPKKKRK<br>V |
| 2 | Dead *Dis*Cas7-11 | MKRTADGSEFESPKKKRKVTTTMKISIEFLEPFRMTKWQESTRRNKNNKEFVRGQAFARWHRNKKDNTKGRPYITGTLLRSAVIRS<br>AENLLTLSDGKISEKTCCPGKFDTEDKDRLLQLRQRSTLRWTDKNPCPDNAETYCPFCELLGRSGNDGKKAEKKDWRFRIHFGNLS<br>LPGKPDFDGPKAIGSQRVLNRVDFKSGKAHDFFKAYEVDHTRFPRFEGEITIDNKVSAEARKLLCDSLKFTDRLCGALCVIRFDEYTP<br>AADSGKQTENVQAEPNANLAEKTAEQIISILDDNKKTEYTRLLADAIRSLRRSSKLVAGLPKDHDGKDDHYLWDIGKKKKDENSVT<br>IRQILTTSADTKELKNAGKWREFCEKLGEALYLKSKDMSGGLKITRRILGDAEFHGKPDRLEKSRSVSIGSVLKETVVCGELVAKTPF<br>FFGAIDEDAKQTALQVLLTPDNKYRLPRSAVRGILRRDLQTYFDSPCNAELGGRPCMCKTCRIMRGITVMDARSEYNAPPEIRHRTRI<br>NPFTGTVAEGALFNMEVAPEGIVFPFQLRYRGSEDGLPDALKTVLKWWAEGQAFMSGAASTGKGRFRMENAKYETLDLSDENQR<br>NDYLKNWGWRDEKGLEELKKRLNSGLPEPGNYRDPKWHEINVSIEMASPFINGDPIRAAVDKRGTAVVTFVKYKAEGEEAKPVCA<br>YKAESFRGVIRSAVARIHMEDGVPLTELTHSDCECLLCQIFGSEYEAGKIRFEDLVFESDPEPVTFDHVAIDRFTGGAADKKKFDDSP<br>LPGSPARPLMLKGSFWIRRDVLEDEEYCKALGKALADVNNGLYPLGGKSAIGYGQVKSLGIKGDDKRISRLMNPAFDETDVAVPEK<br>PKTDAEVRIEAEKVYYPHYFVEPHKKVEREEKPCGHQKFHEGRLTGKIRCKLITKTPLIVPDTSNDDFFRPADKEARKEKDEYHKSY<br>AFFRLHKQIMIPGSELRGMVSSVYETVTNSCFRIFDETKRLSWRMDADHQNVLQDFLPGRVTADGKHIQKFSETARVPFYDKTQKH<br>FDILDEQEIAGEKPVRMWVKRFIKRLSLVDPAKHPQKKQDNKWKRRKEGIATFIEQKNGSYYFNVVTNNGCTSFHLWHKPDNFDQ<br>EKLEGIQNGEKLDCWVRDSRYQKAFQEIPENDPDGWECKEGYLHVVGPSKVEFSDKKGDVINNFQGTLPSVPNDWKTIRTNDFKN<br>RKRKNEPVFCCEDDKGNYYTMAKYCETFFFDLKENEEYEIPEKARIKYKELLRVYNNNPQAVPESVFQSRVARENVEKLKSGDLVY<br>FKHNEKYVEDIVPVRISRTVDDRMIGKRMSADLRPCHGDWVEDGDLSALNAYPEKRLLLRHPKGLCPACRLFGTGSYKGRVRFGF<br>ASLENDPEWLIPGKNPGDPFHGGPVMLSLLERPRPTWSIPGSDNKFKVPGRKFYVHHHAWKTIKDGNHPTTGKAIEQSPNNRTVEAL<br>AGGNSFSFEIAFENLKEWELGLLIHSLQLEKGLAHKLGMAKSMGFGSVEIDVESVRLRKDWKQWRNGNSEIPNWLGKGFAKLKEW<br>FRDELDFIENLKKLLWFPEGDQAPRVCYPMLRKKDDPNGNSGYEELKDGEFKKEDRQKKLTTPWTPWAKRTADGSEFESPKKKRK<br>V |
| 3 | *Lwa*Cas13 | MKVTKVDGISHKKYIEEGKLVKSTSEENRTSERLSELLSIRLDIYIKNPDNASEEENRIRRENLKKFFSNKVLHLKDSVLYLKNRKEK<br>NAVQDKNYSEEDISEYDLKNKNSFSVLKKILLNEDVNSEELEIFRKDVEAKLNKINSLKYSFEENKANYQKINENNVEKVGGKSKRN<br>IIYDYYRESAKRNDYINNVQEAFDKLYKKEDIEKLFFLIENSKKHEKYKIREYYHKIIGRKNDKENFAKIIYEEIQNVNNIKELIEKIPD<br>MSELKKSQVFYKYYLDKEELNDKNIKYAFCHFVEIEMSQLLKNYVYKRLSNISNDKIKRIFEYQNLKKLIENKLLNKLDTYVRNCGK<br>YNYYLQVGEIATSDFIARNRQNEAFLRNIIGVSSVAYFSLRNILETENENGITGRMRGKTVKNNKGEEKYVSGEVDKIYNENKQNEV<br>KENLKMFYSYDFNMDNKNEIEDFFANIDEAISSIRHGIVHFNLELEGKDIFAFKNIAPSEISKKMFQNEINEKKLKLKIFKQLNSANVF<br>NYYEKDVIIKYLKNTKFNFVNKNIPFVPSFTKLYNKIEDLRNTLKFFWSVPKDKEEKDAQIYLLKNIYYGEFLNKFVKNSKVFFKITN<br>EVIKINKQRNQKTGHYKYQKFENIEKTVPVEYLAIIQSREMINNQDKEEKNTYIDFIQQIFLKGFIDYLNKNNLKYIESNNNNDNNDIF<br>SKIKIKKDNKEKYDKILKNYEKHNRNKEIPHEINEFVREIKLGKILKYTENLNMFYLILKLLNHKELTNLKGSLEKYQSANKEETFSDE<br>LELINLLNLDNNRVTEDFELEANEIGKFLDFNENKIKDRKELKKFDTNKIYFDGENIIKHRAFYNIKKYGMLNLLEKIADKAKYKISL<br>KELKEYSNKKNEIEKNYTMQQNLHRKYARPKKDEKFNDEDYKEYEKAIGNIQKYTHLKNKVEFNELNLLQGLLLKILHRLVGYTSI<br>WERDLRFRLKGEFPENHYIEEIFNFDNSKNVKYKSGQIVEKYINFYKELYKDNVEKRSIYSDKKVKKLKQEKKDLYIRNYIAHFNYIP<br>HAEISLLEVLENLRKLLSYDRKLKNAIMKSIVDILKEYGFVATFKIGADKKIEIQTLESEKIVHLKNLKKKKLMTDRNSEELCELVKV<br>MFEYKALEGGGGSGGGGSGGGGSVSKGEELFTGVVPILVELDGDVNGHKFSVRGEGEGDATNGKLTLKFICTTGKLPVPWPTLVTT<br>LTYGVQCFSRYPDHMKQHDFFKSAMPEGYVQERTISFKDDGTYKTRAEVKFEGDTLVNRIELKGIDFKEDGNILGHKLEYNFNSHN<br>VYITADKQKNGIKANFKIRHNVEDGSVQLADHYQQNTPIGDGPVLLPDNHYLSTQSKLSKDPNEKRDHMVLLEFVTAAGITLGMDE<br>LYKGSEGAPKKKRKVGSSYPYDVPDYAYPYDVPDYAYPYDVPDYAKRTADGSEFES |
| 4 | *Psp*Cas13 | MNIPALVENQKKYFGTYSVMAMLNAQTVLDHIQKVADIEGEQNENNENLWFHPVMSHLYNAKNGYDKQPEKTMFIIERLQSYFPF<br>LKIMAENQREYSNGKYKQNRVEVNSNDIFEVLKRAFGVLKMYRDLTNHYKTYEEKLNDGCEFLTSTEQPLSGMINNYYTVALRNM<br>NERYGYKTEDLAFIQDKRFKFVKDAYGKKKSQVNTGFFLSLQDYNGDTQKKLHLSGVGIALLICLFLDKQYINIFLSRLPIFSSYNAQ<br>SEERRIIIRSFGINSIKLPKDRIHSEKSNKSVAMDMLNEVKRCPDELFTTLSAEKQSRFRIISDDHNEVLMKRSSDRFVPLLLQYIDYGK<br>LFDHIRFHVNMGKLRYLLKADKTCIDGQTRVRVIEQPLNGFGRLEEAETMRKQENGTFGNSGIRIRDFENMKRDDANPANYPYIVD<br>TYTHYILENNKVEMFINDKEDSAPLLPVIEDDRYVVKTIPSCRMSTLEIPAMAFHMFLFGSKKTEKLIVDVHNRYKRLFQAMQKEEV<br>TAENIASFGIAESDLPQKILDLISGNAHGKDVDAFIRLTVDDMLTDTERRIKRFKDDRKSIRSADNKMGKRGFKQISTGKLADFLAKD<br>IVLFQPSVNDGENKITGLNYRIMQSAIAVYDSGDDYEAKQQFKLMFEKARLIGKGTTEPHPFLYKVFARSIPANAVEFYERYLIERKF<br>YLTGLSNEIKKGNRVDVPFIRRDQNKWKTPAMKTLGRIYSEDLPVELPRQMFDNEIKSHLKSLPQMEGIDFNNANVTYLIAEYMKR<br>VLDDDFQTFYQWNRNYRYMDMLKGEYDRKGSLQHCFTSVEEREGLWKERASRTERYRKQASNKIRSNRQMRNASSEEIETILDKR<br>LSNSRNEYQKSEKVIRRYRVQDALLFLLAKKTLTELADFDGERFKLKEIMPDAEKGILSEIMPMSFTFEKGGKKYTITSEGMKLKNY<br>GDFFVLASDKRIGNLLELVGSDIVSKEDIMEEFNKYDQCRPEISSIVFNLEKWAFDTYPELSARVDREEKVDFKSILKILLNNKNINKE<br>QSDILRKIRNAFDHNNYPDKGVVEIKALPEIAMSIKKAFGEYAIMKGSLQLPPLERLTLGSSYPYDVPDYAYPYDVPDYAYPYDVPD<br>YA |
| 5 | *Rfx*Cas13 | MSPKKKRKVEASIEKKKSFAKGMGVKSTLVSGSKVYMTTFAEGSDARLEKIVEGDSIRSVNEGEAFSAEMADKNAGYKIGNAKFSH<br>PKGYAVVANNPLYTGPVQQDMLGLKETLEKRYFGESADGNDNICIQVIHNILDIEKILAEYITNAAYAVNNISGLDKDIIGFGKFSTV<br>YTYDEFKDPEHHRAAFNNNDKLINAIKAQYDEFDNFLDNPRLGYFGQAFFSKEGRNYIINYGNECYDILALLSGLRHWVVHNNEEE<br>SRISRTWLYNLDKNLDNEYISTLNYLYDRITNELTNSFSKNSAANVNYIAETLGINPAEFAEQYFRFSIMKEQKNLGFNITKLREVML<br>DRKDMSEIRKNHKVFDSIRTKVYTMMDFVIYRYYIEEDAKVAAANKSLPDNEKSLSEKDIFVINLRGSFNDDQKDALYYDEANRIW<br>RKLENIMHNIKEFRGNKTREYKKKDAPRLPRILPAGRDVSAFSKLMYALTMFLDGKEINDLLTTLINKFDNIQSFLKVMPLIGVNAKF<br>VEEYAFFKDSAKIADELRLIKSFARMGEPIADARRAMYIDAIRILGTNLSYDELKALADTFSLDENGNKLKKGKHGMRNFIINNVISN<br>KRFHYLIRYGDPAHLHEIAKNEAVVKFVLGRIADIQKKQGQNGKNQIDRYYETCIGKDKGKSVSEKVDALTKIITGMNYDQFDKKR |

| | | |
|---|---|---|
| | | SVIEDTGRENAEREKFKKIISLYLTVIYHILKNIVNINARYVIGFHCVERDAQLYKEKGYDINLKKLEEKGFSSVTKLCAGIDETAPDK<br>RKDVEKEMAERAKESIDSLESANPKLYANYIKYSDEKKAEEFTRQINREKAKTALNAYLRNTKWNVIIREDLLRIDNKTCTLFRNKA<br>VHLEVARYVHAYINDIAEVNSYFQLYHYIMQRIIMNERYEKSSGKVSEYFDAVNDEKKYNDRLLKLLCVPFGYCIPRFKNLSIEALF<br>DRNEAAKFDKEKKKVSGNSGSGPKKKRKVAAAYPYDVPDYAASGSGKRTADGSEFES |
| 6 | *Lwa*Cas13 with NLS | MKRTADGSEFESPKKKRKVKVTKVDGISHKKYIEEGKLVKSTSEENRTSERLSELLSIRLDIYIKNPDNASEEENRIRRENLKKFFSNK<br>VLHLKDSVLYLKNRKEKNAVQDKNYSEEDISEYDLKNKNSFSVLKKILLNEDVNSEELEIFRKDVEAKLNKINSLKYSFEENKANYQ<br>KINENNVEKVGGKSKRNIIYDYYRESAKRNDYINNVQEAFDKLYKKEDIEKLFFLIENSKKHEKYKIREYYHKIIGRKNDKENFAKII<br>YEEIQNVNNIKELIEKIPDMSELKKSQVFYKYYLDKEELNDKNIKYAFCHFVEIEMSQLLKNYVYKRLSNISNDKIKRIFEYQNLKKLI<br>ENKLLNKLDTYVRNCGKYNYYLQVGEIATSDFIARNRQNEAFLRNIIGVSSVAYFSLRNILETENENGITGRMRGKTVKNNKGEEKY<br>VSGEVDKIYNENKQNEVKENLKMFYSYDFNMDNKNEIEDFFANIDEAISSIRHGIVHFNLELEGKDIFAFKNIAPSEISKKMFQNEINE<br>KKLKLKIFKQLNSANVFNYYEKDVIIKYLKNTKFNFVNKNIPFVPSFTKLYNKIEDLRNTLKFFWSVPKDKEEKDAQIYLLKNIYYGE<br>FLNKFVKNSKVFFKITNEVIKINKQRNQKTGHYKYQKFENIEKTVPVEYLAIIQSREMINNQDKEEKNTYIDFIQQIFLKGFIDYLNKN<br>NLKYIESNNNNDNNDIFSKIKIKKDNKEKYDKILKNYEKHNRNKEIPHEINEFVREIKLGKILKYTENLNMFYLILKLLNHKELTNLKG<br>SLEKYQSANKEETFSDELELINLLNLDNNRVTEDFELEANEIGKFLDFNENKIKDRKELKKFDTNKIYFDGENIIKHRAFYNIKKYGM<br>LNLLEKIADKAKYKISLKELKEYSNKKNEIEKNYTMQQNLHRKYARPKKDEKFNDEDYKEYEKAIGNIQKYTHLKNKVEFNELNLL<br>QGLLLKILHRLVGYTSIWERDLRFRLKGEFPENHYIEEIFNFDNSKNVKYKSGQIVEKYINFYKELYKDNVEKRSIYSDKKVKKLKQE<br>KKDLYIRNYIAHFNYIPHAEISLLEVLENLRKLLSYDRKLKNAIMKSIVDILKEYGFVATFKIGADKKIEIQTLESEKIVHLKNLKKKKL<br>MTDRNSEELCELVKVMFEYKALEGGGGSGGGGSGGGGSVSKGEELFTGVVPILVELDGDVNGHKFSVRGEGEGDATNGKLTLKFI<br>CTTGKLPVPWPTLVTTLTYGVQCFSRYPDHMKQHDFFKSAMPEGYVQERTISFKDDGTYKTRAEVKFEGDTLVNRIELKGIDFKED<br>GNILGHKLEYNFNSHNVYITADKQKNGIKANFKIRHNVEDGSVQLADHYQQNTPIGDGPVLLPDNHYLSTQSKLSKDPNEKRDHMV<br>LLEFVTAAGITLGMDELYKGSEGAPKKKRKVGSSYPYDVPDYAYPYDVPDYAYPYDVPDYAKRTADGSEFESPKKKRKV |
| 7 | PspCas13 with NLS | MKRTADGSEFESPKKKRKVNIPALVENQKKYFGTYSVMAMLNAQTVLDHIQKVADIEGEQNENNENLWFHPVMSHLYNAKNGYD<br>KQPEKTMFIIERLQSYFPFLKIMAENQREYSNGKYKQNRVEVNSNDIFEVLKRAFGVLKMYRDLTNHYKTYEEKLNDGCEFLTSTEQ<br>PLSGMINNYYTVALRNMNERYGYKTEDLAFIQDKRFKFVKDAYGKKKSQVNTGFFLSLQDYNGDTQKKLHLSGVGIALLICLFLDK<br>QYINIFLSRLPIFSSYNAQSEERRIIIRSFGINSIKLPKDRIHSEKSNKSVAMDMLNEVKRCPDELFTTLSAEKQSRFRIISDDHNEVLMKR<br>SSDRFVPLLLQYIDYGKLFDHIRFHVNMGKLRYLLKADKTCIDGQTRVRVIEQPLNGFGRLEEAETMRKQENGTFGNSGIRIRDFEN<br>MKRDDANPANYPYIVDTYTHYILENNKVEMFINDKEDSAPLLPVIEDDRYVVKTIPSCRMSTLEIPAMAFHMFLFGSKKTEKLIVDV<br>HNRYKRLFQAMQKEEVTAENIASFGIAESDLPQKILDLISGNAHGKDVDAFIRLTVDDMLTDTERRIKRFKDDRKSIRSADNKMGKR<br>GFKQISTGKLADFLAKDIVLFQPSVNDGENKITGLNYRIMQSAIAVYDSGDDYEAKQQFKLMFEKARLIGKGTTEPHPFLYKVFARSI<br>PANAVEFYERYLIERKFYLTGLSNEIKKGNRVDVPFIRRDQNKWKTPAMKTLGRIYSEDLPVELPRQMFDNEIKSHLKSLPQMEGIDF<br>NNANVTYLIAEYMKRVLDDDFQTFYQWNRNYRYMDMLKGEYDRKGSLQHCFTSVEEREGLWKERASRTERYRKQASNKIRSNR<br>QMRNASSEEIETILDKRLSNSRNEYQKSEKVIRRYRVQDALLFLLAKKTLTELADFDGERFKLKEIMPDAEKGILSEIMPMSFTFEKG<br>GKKYTITSEGMKLKNYGDFFVLASDKRIGNLLELVGSDIVSKEDIMEEFNKYDQCRPEISSIVFNLEKWAFDTYPELSARVDREEKVD<br>FKSILKILLNNKNINKEQSDILRKIRNAFDHNNYPDKGVVEIKALPEIAMSIKKAFGEYAIMKGSLQLPPLERLTLGSSYPYDVPDYAY<br>PYDVPDYAYPYDVPDYAKRTADGSEFESPKKKRKV |
| 8 | *Rfx*Cas13 with NLS | MKRTADGSEFESPKKKRKVSPKKKRKVEASIEKKKSFAKGMGVKSTLVSGSKVYMTTFAEGSDARLEKIVEGDSIRSVNEGEAFSA<br>EMADKNAGYKIGNAKFSHPKGYAVVANNPLYTGPVQQDMLGLKETLEKRYFGESADGNDNICIQVIHNILDIEKILAEYITNAAYA<br>VNNISGLDKDIIGFGKFSTVYTYDEFKDPEHHRAAFNNNDKLINAIKAQYDEFDNFLDNPRLGYFGQAFFSKEGRNYIINYGNECYDI<br>LALLSGLRHWVVHNNEEESRISRTWLYNLDKNLDNEYISTLNYLYDRITNELTNSFSKNSAANVNYIAETLGINPAEFAEQYFRFSIM<br>KEQKNLGFNITKLREVMLDRKDMSEIRKNHKVFDSIRTKVYTMMDFVIYRYYIEEDAKVAAANKSLPDNEKSLSEKDIFVINLRGSF<br>NDDQKDALYYDEANRIWRKLENIMHNIKEFRGNKTREYKKKDAPRLPRILPAGRDVSAFSKLMYALTMFLDGKEINDLLTTLINKF<br>DNIQSFLKVMPLIGVNAKFVEEYAFFKDSAKIADELRLIKSFARMGEPIADARRAMYIDAIRILGTNLSYDELKALADTFSLDENGNK<br>LKKGKHGMRNFIINNVISNKRFHYLIRYGDPAHLHEIAKNEAVVKFVLGRIADIQKKQGQNGKNQIDRYYETCIGKDKGKSVSEKVD<br>ALTKIITGMNYDQFDKKRSVIEDTGRENAEREKFKKIISLYLTVIYHILKNIVNINARYVIGFHCVERDAQLYKEKGYDINLKKLEEKG<br>FSSVTKLCAGIDETAPDKRKDVEKEMAERAKESIDSLESANPKLYANYIKYSDEKKAEEFTRQINREKAKTALNAYLRNTKWNVIIR<br>EDLLRIDNKTCTLFRNKAVHLEVARYVHAYINDIAEVNSYFQLYHYIMQRIIMNERYEKSSGKVSEYFDAVNDEKKYNDRLLKLLC<br>VPFGYCIPRFKNLSIEALFDRNEAAKFDKEKKKVSGNSGSGPKKKRKVAAAYPYDVPDYAASGSGKRTADGSEFESPKKKRKV |
| 9 | *Dis*Cas7-11 D1580R | MKRTADGSEFESPKKKRKVTTTMKISIEFLEPFRMTKWQESTRRNKNNKEFVRGQAFARWHRNKKDNTKGRPYITGTLLRSAVIRS<br>AENLLTLSDGKISEKTCCPGKFDTEDKDRLLQLRQRSTLRWTDKNPCPDNAETYCPFCELLGRSGNDGKKAEKKDWRFRIHFGNLS<br>LPGKPDFDGPKAIGSQRVLNRVDFKSGKAHDFFKAYEVDHTRFPRFEGEITIDNKVSAEARKLLCDSLKFTDRLCGALCVIRFDEYTP<br>AADSGKQTENVQAEPNANLAEKTAEQIISILDDNKKTEYTRLLADAIRSLRRSSKLVAGLPKDHDGKDDHYLWDIGKKKKDENSVT<br>IRQILTTSADTKELKNAGKWREFCEKLGEALYLKSKDMSGGLKITRRILGDAEFHGKPDRLEKSRSVSIGSVLKETVVCGELVAKTPF<br>FFGAIDEDAKQTDLQVLLTPDNKYRLPRSAVRGILRRDLQTYFDSPCNAELGGRPCMCKTCRIMRGITVMDARSEYNAPPEIRHRTRI<br>NPFTGTVAEGALFNMEVAPEGIVFPFQLRYRGSEDGLPDALKTVLKWWAEGQAFMSGAASTGKGRFRMENAKYETLDLSDENQR<br>NDYLKNWGWRDEKGLEELKKRLNSGLPEPGNYRDPKWHEINVSIEMASPFINGDPIRAAVDKRGTDVVTFVKYKAEGEEAKPVCA<br>YKAESFRGVIRSAVARIHMEDGVPLTELTHSDCECLLCQIFGSEYEAGKIRFEDLVFESDPEPVTFDHVAIDRFTGGAADKKKFDDSP<br>LPGSPARPLMLKGSFWIRRDVLEDEEYCKALGKALADVNNGLYPLGGKSAIGYGQVKSLGIKGDDKRISRLMNPAFDETDVAVPEK<br>PKTDAEVRIEAEKVYYPHYFVEPHKKVEREEKPCGHQKFHEGRLTGKIRCKLITKTPLIVPDTSNDDFFRPADKEARKEKDEYHKSY<br>AFFRLHKQIMIPGSELRGMVSSVYETVTNSCFRIFDETKRLSWRMDADHQNVLQDFLPGRVTADGKHIQKFSETARVPFYDKTQKH<br>FDILDEQEIAGEKPVRMWVKRFIKRLSLVDPAKHPQKKQDNKWKRRKEGIATFIEQKNGSYYFNVVTNNGCTSFHLWHKPDNFDQ<br>EKLEGIQNGEKLDCWVRDSRYQKAFQEIPENDPDGWECKEGYLHVVGPSKVEFSDKKGDVINNFQGTLPSVPNDWKTIRTNDFKN<br>RKRKNEPVFCCEDDKGNYYTMAKYCETFFFDLKENEEYEIPEKARIKYKELLRVYNNNPQAVPESVFQSRVARENVEKLKSGDLVY<br>FKHNEKYVEDIVPVRISRTVDDRMIGKRMSADLRPCHGDWVEDGDLSALNAYPEKRLLLRHPKGLCPACRLFGTGSYKGRVRFGF<br>ASLENDPEWLIPGKNPGDPFHGGPVMLSLLERPRPTWSIPGSDNKFKVPGRKFYVHHHAWKTIKDGNHPTTGKAIEQSPNNRTVEAL<br>AGGNSFSFEIAFENLKEWELGLLIHSLQLEKGLAHKLGMAKSMGFGSVEIDVESVRLRKDWKQWRNGNSEIPNWLGKGFAKLKEW<br>FRDELDFIENLKKLLWFPEGDQAPRVCYPMLRKKDDPNGNSGYEELKRGEFKKEDRQKKLTTPWTPWAKRTADGSEFESPKKKRK<br>V |
| 10 | *Dis*Cas7-11 D1580R D988K Y312K | MKRTADGSEFESPKKKRKVTTTMKISIEFLEPFRMTKWQESTRRNKNNKEFVRGQAFARWHRNKKDNTKGRPYITGTLLRSAVIRS<br>AENLLTLSDGKISEKTCCPGKFDTEDKDRLLQLRQRSTLRWTDKNPCPDNAETYCPFCELLGRSGNDGKKAEKKDWRFRIHFGNLS<br>LPGKPDFDGPKAIGSQRVLNRVDFKSGKAHDFFKAYEVDHTRFPRFEGEITIDNKVSAEARKLLCDSLKFTDRLCGALCVIRFDEYTP<br>AADSGKQTENVQAEPNANLAEKTAEQIISILDDNKKTEYTRLLADAIRSLRRSSKLVAGLPKDHDGKDDHKLWDIGKKKKDENSVT<br>IRQILTTSADTKELKNAGKWREFCEKLGEALYLKSKDMSGGLKITRRILGDAEFHGKPDRLEKSRSVSIGSVLKETVVCGELVAKTPF<br>FFGAIDEDAKQTDLQVLLTPDNKYRLPRSAVRGILRRDLQTYFDSPCNAELGGRPCMCKTCRIMRGITVMDARSEYNAPPEIRHRTRI<br>NPFTGTVAEGALFNMEVAPEGIVFPFQLRYRGSEDGLPDALKTVLKWWAEGQAFMSGAASTGKGRFRMENAKYETLDLSDENQR<br>NDYLKNWGWRDEKGLEELKKRLNSGLPEPGNYRDPKWHEINVSIEMASPFINGDPIRAAVDKRGTDVVTFVKYKAEGEEAKPVCA<br>YKAESFRGVIRSAVARIHMEDGVPLTELTHSDCECLLCQIFGSEYEAGKIRFEDLVFESDPEPVTFDHVAIDRFTGGAADKKKFDDSP<br>LPGSPARPLMLKGSFWIRRDVLEDEEYCKALGKALADVNNGLYPLGGKSAIGYGQVKSLGIKGDDKRISRLMNPAFDETDVAVPEK<br>PKTDAEVRIEAEKVYYPHYFVEPHKKVEREEKPCGHQKFHEGRLTGKIRCKLITKTPLIVPDTSNDDFFRPADKEARKEKDEYHKSY<br>AFFRLHKQIMIPGSELRGMVSSVYETVTNSCFRIFDETKRLSWRMDADHQNVLQKFLPGRVTADGKHIQKFSETARVPFYDKTQKH<br>FDILDEQEIAGEKPVRMWVKRFIKRLSLVDPAKHPQKKQDNKWKRRKEGIATFIEQKNGSYYFNVVTNNGCTSFHLWHKPDNFDQ<br>EKLEGIQNGEKLDCWVRDSRYQKAFQEIPENDPDGWECKEGYLHVVGPSKVEFSDKKGDVINNFQGTLPSVPNDWKTIRTNDFKN<br>RKRKNEPVFCCEDDKGNYYTMAKYCETFFFDLKENEEYEIPEKARIKYKELLRVYNNNPQAVPESVFQSRVARENVEKLKSGDLVY<br>FKHNEKYVEDIVPVRISRTVDDRMIGKRMSADLRPCHGDWVEDGDLSALNAYPEKRLLLRHPKGLCPACRLFGTGSYKGRVRFGF<br>ASLENDPEWLIPGKNPGDPFHGGPVMLSLLERPRPTWSIPGSDNKFKVPGRKFYVHHHAWKTIKDGNHPTTGKAIEQSPNNRTVEAL |

| | | |
|---|---|---|
| | | AGGNSFSFEIAFENLKEWELGLLIHSLQLEKGLAHKLGMAKSMGFGSVEIDVESVRLRKDWKQWRNGNSEIPNWLGKGFAKLKEW<br>FRDELDFIENLKKLLWFPEGDQAPRVCYPMLRKKDDPNGNSGYEELKRGEFKKEDRQKKLTTPWTPWAKRTADGSEFESPKKKRK<br>V |
| 11 | *Dis*Cas7-11 SF3B6 C-terminal fusion | MKRTADGSEFESPKKKRKVTTTMKISIEFLEPFRMTKWQESTRRNKNNKEFVRGQAFARWHRNKKDNTKGRPYITGTLLRSAVIRS<br>AENLLTLSDGKISEKTCCPGKFDTEDKDRLLQLRQRSTLRWTDKNPCPDNAETYCPFCELLGRSGNDGKKAEKKDWRFRIHFGNLS<br>LPGKPDFDGPKAIGSQRVLNRVDFKSGKAHDFFKAYEVDHTRFPRFEGEITIDNKVSAEARKLLCDSLKFTDRLCGALCVIRFDEYTP<br>AADSGKQTENVQAEPNANLAEKTAEQIISILDDNKKTEYTRLLADAIRSLRRSSKLVAGLPKDHDGKDDHYLWDIGKKKKDENSVT<br>IRQILTTSADTKELKNAGKWREFCEKLGEALYLKSKDMSGGLKITRRILGDAEFHGKPDRLEKSRSVSIGSVLKETVVCGELVAKTPF<br>FFGAIDEDAKQTDLQVLLTPDNKYRLPRSAVRGILRRDLQTYFDSPCNAELGGRPCMCKTCRIMRGITVMDARSEYNAPPEIRHRTRI<br>NPFTGTVAEGALFNMEVAPEGIVFPFQLRYRGSEDGLPDALKTVLKWWAEGQAFMSGAASTGKGRFRMENAKYETLDLSDENQR<br>NDYLKNWGWRDEKGLEELKKRLNSGLPEPGNYRDPKWHEINVSIEMASPFINGDPIRAAVDKRGTDVVTFVKYKAEGEEAKPVCA<br>YKAESFRGVIRSAVARIHMEDGVPLTELTHSDCECLLCQIFGSEYEAGKIRFEDLVFESDPEPVTFDHVAIDRFTGGAADKKKFDDSP<br>LPGSPARPLMLKGSFWIRRDVLEDEEYCKALGKALADVNNGLYPLGGKSAIGYGQVKSLGIKGDDKRISRLMNPAFDETDVAVPEK<br>PKTDAEVRIEAEKVYYPHYFVEPHKKVEREEKPCGHQKFHEGRLTGKIRCKLITKTPLIVPDTSNDDFFRPADKEARKEKDEYHKSY<br>AFFRLHKQIMIPGSELRGMVSSVYETVTNSCFRIFDETKRLSWRMDADHQNVLQDFLPGRVTADGKHIQKFSETARVPFYDKTQKH<br>FDILDEQEIAGEKPVRMWVKRFIKRLSLVDPAKHPQKKQDNKWKRRKEGIATFIEQKNGSYYFNVVTNNGCTSFHLWHKPDNFDQ<br>EKLEGIQNGEKLDCWVRDSRYQKAFQEIPENDPDGWECKEGYLHVVGPSKVEFSDKKGDVINNFQGTLPSVPNDWKTIRTNDFKN<br>RKRKNEPVFCCEDDKGNYYTMAKYCETFFFDLKENEEYEIPEKARIKYKELLRVYNNNPQAVPESVFQSRVARENVEKLKSGDLVY<br>FKHNEKYVEDIVPVRISRTVDDRMIGKRMSADLRPCHGDWVEDGDLSALNAYPEKRLLLRHPKGLCPACRLFGTGSYKGRVRFGF<br>ASLENDPEWLIPGKNPGDPFHGGPVMLSLLERPRPTWSIPGSDNKFKVPGRKFYVHHHAWKTIKDGNHPTTGKAIEQSPNNRTVEAL<br>AGGNSFSFEIAFENLKEWELGLLIHSLQLEKGLAHKLGMAKSMGFGSVEIDVESVRLRKDWKQWRNGNSEIPNWLGKGFAKLKEW<br>FRDELDFIENLKKLLWFPEGDQAPRVCYPMLRKKDDPNGNSGYEELKDGEFKKEDRQKKLTTPWTPWAMAMQAAKRANIRLPPE<br>VNRILYIRNLPYKITAEEMYDIFGKYGPIRQIRVGNTPETRGTAYVVYEDIFDAKNACDHLSGFNVCNRYLVVLYYNANRAFQKMD<br>TKKKEEQLKLLKEKYGINTDPPKKRTADGSEFESPKKKRKV |
| 12 | *Dis*Cas7-11 U2AF2 C-terminal fusion | MKRTADGSEFESPKKKRKVTTTMKISIEFLEPFRMTKWQESTRRNKNNKEFVRGQAFARWHRNKKDNTKGRPYITGTLLRSAVIRS<br>AENLLTLSDGKISEKTCCPGKFDTEDKDRLLQLRQRSTLRWTDKNPCPDNAETYCPFCELLGRSGNDGKKAEKKDWRFRIHFGNLS<br>LPGKPDFDGPKAIGSQRVLNRVDFKSGKAHDFFKAYEVDHTRFPRFEGEITIDNKVSAEARKLLCDSLKFTDRLCGALCVIRFDEYTP<br>AADSGKQTENVQAEPNANLAEKTAEQIISILDDNKKTEYTRLLADAIRSLRRSSKLVAGLPKDHDGKDDHYLWDIGKKKKDENSVT<br>IRQILTTSADTKELKNAGKWREFCEKLGEALYLKSKDMSGGLKITRRILGDAEFHGKPDRLEKSRSVSIGSVLKETVVCGELVAKTPF<br>FFGAIDEDAKQTDLQVLLTPDNKYRLPRSAVRGILRRDLQTYFDSPCNAELGGRPCMCKTCRIMRGITVMDARSEYNAPPEIRHRTRI<br>NPFTGTVAEGALFNMEVAPEGIVFPFQLRYRGSEDGLPDALKTVLKWWAEGQAFMSGAASTGKGRFRMENAKYETLDLSDENQR<br>NDYLKNWGWRDEKGLEELKKRLNSGLPEPGNYRDPKWHEINVSIEMASPFINGDPIRAAVDKRGTDVVTFVKYKAEGEEAKPVCA<br>YKAESFRGVIRSAVARIHMEDGVPLTELTHSDCECLLCQIFGSEYEAGKIRFEDLVFESDPEPVTFDHVAIDRFTGGAADKKKFDDSP<br>LPGSPARPLMLKGSFWIRRDVLEDEEYCKALGKALADVNNGLYPLGGKSAIGYGQVKSLGIKGDDKRISRLMNPAFDETDVAVPEK<br>PKTDAEVRIEAEKVYYPHYFVEPHKKVEREEKPCGHQKFHEGRLTGKIRCKLITKTPLIVPDTSNDDFFRPADKEARKEKDEYHKSY<br>AFFRLHKQIMIPGSELRGMVSSVYETVTNSCFRIFDETKRLSWRMDADHQNVLQDFLPGRVTADGKHIQKFSETARVPFYDKTQKH<br>FDILDEQEIAGEKPVRMWVKRFIKRLSLVDPAKHPQKKQDNKWKRRKEGIATFIEQKNGSYYFNVVTNNGCTSFHLWHKPDNFDQ<br>EKLEGIQNGEKLDCWVRDSRYQKAFQEIPENDPDGWECKEGYLHVVGPSKVEFSDKKGDVINNFQGTLPSVPNDWKTIRTNDFKN<br>RKRKNEPVFCCEDDKGNYYTMAKYCETFFFDLKENEEYEIPEKARIKYKELLRVYNNNPQAVPESVFQSRVARENVEKLKSGDLVY<br>FKHNEKYVEDIVPVRISRTVDDRMIGKRMSADLRPCHGDWVEDGDLSALNAYPEKRLLLRHPKGLCPACRLFGTGSYKGRVRFGF<br>ASLENDPEWLIPGKNPGDPFHGGPVMLSLLERPRPTWSIPGSDNKFKVPGRKFYVHHHAWKTIKDGNHPTTGKAIEQSPNNRTVEAL<br>AGGNSFSFEIAFENLKEWELGLLIHSLQLEKGLAHKLGMAKSMGFGSVEIDVESVRLRKDWKQWRNGNSEIPNWLGKGFAKLKEW<br>FRDELDFIENLKKLLWFPEGDQAPRVCYPMLRKKDDPNGNSGYEELKDGEFKKEDRQKKLTTPWTPWAMSDFDEFERQLNENKQE<br>RDKENRHRKRSHSRSRSRDRKRRSRSRDRRNRDQRSASRDRRRRSKPLTRGAKEEHGGLIRSPRHEKKKKVRKYWDVPPPGFEHIT<br>PMQYKAMQAAGQIPATALLPTMTPDGLAVTPTPVPVVGSQMTRQARRLYVGNIPFGITEEAMMDFFNAQMRLGGLTQAPGNPVLA<br>VQINQDKNFAFLEFRSVDETTQAMAFDGIIFQGQSLKIRRPHDYQPLPGMSENPSVYVPGVVSTVVPDSAHKLFIGGLPNYLNDDQV<br>KELLTSFGPLKAFNLVKDSATGLSKGYAFCEYVDINVTDQAIAGLNGMQLGDKKLLVQRASVGAKNATLVSPPSTINQTPVTLQVP<br>GLMSSQVQMGGHPTEVLCLMNMVLPEELLDDEEYEEIVEDVRDECSKYGLVKSIEIPRPVDGVEVPGCGKIFVEFTSVFDCQKAMQ<br>GLTGRKFANRVVVTKYCDPDSYHRRDFWKRTADGSEFESPKKKRKV |
| 13 | *Dis*Cas7-11 RBM17 N-terminal fusion | MKRTADGSEFESPKKKRKVMSLYDDLGVETSDSKTEGWSKNFKLLQSQLQVKKAALTQAKSQRTKQSTVLAPVIDLKRGGSSDDR<br>QIVDTPPHVAAGLKDPVPSGFSAGEVLIPLADEYDPMFPNDYEKVVKRQREERQRQRELERQKEIEEREKRRKDRHEASGFARRPDP<br>DSDEDEDYERERRKRSMGGAAIAPPTSLVEKDKELPRDFPYEEDSRPRSQSSKAAIPPPVYEEQDRPRSPTGPSNSFLANMGGTVAH<br>KIMQKYGFREGQGLGKHEQGLSTALSVEKTSKRGGKIIVGDATEKDASKKSDSNPLTEILKCPTKVVLLRNMVGAGEVDEDLEVET<br>KEECEKYGKVGKCVIFEIPGAPDDEAVRIFLEFERVESAIKAVVDLNGRYFGGRVVKACFYNLDKFRVLDLAEQVTTTMKISIEFLEP<br>FRMTKWQESTRRNKNNKEFVRGQAFARWHRNKKDNTKGRPYITGTLLRSAVIRSAENLLTLSDGKISEKTCCPGKFDTEDKDRLLQ<br>LRQRSTLRWTDKNPCPDNAETYCPFCELLGRSGNDGKKAEKKDWRFRIHFGNLSLPGKPDFDGPKAIGSQRVLNRVDFKSGKAHDF<br>FKAYEVDHTRFPRFEGEITIDNKVSAEARKLLCDSLKFTDRLCGALCVIRFDEYTPAADSGKQTENVQAEPNANLAEKTAEQIISILD<br>DNKKTEYTRLLADAIRSLRRSSKLVAGLPKDHDGKDDHYLWDIGKKKKDENSVTIRQILTTSADTKELKNAGKWREFCEKLGEALY<br>LKSKDMSGGLKITRRILGDAEFHGKPDRLEKSRSVSIGSVLKETVVCGELVAKTPFFFGAIDEDAKQTDLQVLLTPDNKYRLPRSAVR<br>GILRRDLQTYFDSPCNAELGGRPCMCKTCRIMRGITVMDARSEYNAPPEIRHRTRINPFTGTVAEGALFNMEVAPEGIVFPFQLRYRG<br>SEDGLPDALKTVLKWWAEGQAFMSGAASTGKGRFRMENAKYETLDLSDENQRNDYLKNWGWRDEKGLEELKKRLNSGLPEPGN<br>YRDPKWHEINVSIEMASPFINGDPIRAAVDKRGTDVVTFVKYKAEGEEAKPVCAYKAESFRGVIRSAVARIHMEDGVPLTELTHSDC<br>ECLLCQIFGSEYEAGKIRFEDLVFESDPEPVTFDHVAIDRFTGGAADKKKFDDSPLPGSPARPLMLKGSFWIRRDVLEDEEYCKALGK<br>ALADVNNGLYPLGGKSAIGYGQVKSLGIKGDDKRISRLMNPAFDETDVAVPEKPKTDAEVRIEAEKVYYPHYFVEPHKKVEREEKP<br>CGHQKFHEGRLTGKIRCKLITKTPLIVPDTSNDDFFRPADKEARKEKDEYHKSYAFFRLHKQIMIPGSELRGMVSSVYETVTNSCFRIF<br>DETKRLSWRMDADHQNVLQDFLPGRVTADGKHIQKFSETARVPFYDKTQKHFDILDEQEIAGEKPVRMWVKRFIKRLSLVDPAKH<br>PQKKQDNKWKRRKEGIATFIEQKNGSYYFNVVTNNGCTSFHLWHKPDNFDQEKLEGIQNGEKLDCWVRDSRYQKAFQEIPENDPD<br>GWECKEGYLHVVGPSKVEFSDKKGDVINNFQGTLPSVPNDWKTIRTNDFKNRKRKNEPVFCCEDDKGNYYTMAKYCETFFFDLKE<br>NEEYEIPEKARIKYKELLRVYNNNPQAVPESVFQSRVARENVEKLKSGDLVYFKHNEKYVEDIVPVRISRTVDDRMIGKRMSADLRP<br>CHGDWVEDGDLSALNAYPEKRLLLRHPKGLCPACRLFGTGSYKGRVRFGFASLENDPEWLIPGKNPGDPFHGGPVMLSLLERPRPT<br>WSIPGSDNKFKVPGRKFYVHHHAWKTIKDGNHPTTGKAIEQSPNNRTVEALAGGNSFSFEIAFENLKEWELGLLIHSLQLEKGLAHK<br>LGMAKSMGFGSVEIDVESVRLRKDWKQWRNGNSEIPNWLGKGFAKLKEWFRDELDFIENLKKLLWFPEGDQAPRVCYPMLRKKD<br>DPNGNSGYEELKDGEFKKEDRQKKLTTPWTPWAKRTADGSEFESPKKKRKV |
| 14 | *Dis*Cas7-11 RBM17 C-terminal fusion | MKRTADGSEFESPKKKRKVTTTMKISIEFLEPFRMTKWQESTRRNKNNKEFVRGQAFARWHRNKKDNTKGRPYITGTLLRSAVIRS<br>AENLLTLSDGKISEKTCCPGKFDTEDKDRLLQLRQRSTLRWTDKNPCPDNAETYCPFCELLGRSGNDGKKAEKKDWRFRIHFGNLS<br>LPGKPDFDGPKAIGSQRVLNRVDFKSGKAHDFFKAYEVDHTRFPRFEGEITIDNKVSAEARKLLCDSLKFTDRLCGALCVIRFDEYTP<br>AADSGKQTENVQAEPNANLAEKTAEQIISILDDNKKTEYTRLLADAIRSLRRSSKLVAGLPKDHDGKDDHYLWDIGKKKKDENSVT<br>IRQILTTSADTKELKNAGKWREFCEKLGEALYLKSKDMSGGLKITRRILGDAEFHGKPDRLEKSRSVSIGSVLKETVVCGELVAKTPF<br>FFGAIDEDAKQTDLQVLLTPDNKYRLPRSAVRGILRRDLQTYFDSPCNAELGGRPCMCKTCRIMRGITVMDARSEYNAPPEIRHRTRI<br>NPFTGTVAEGALFNMEVAPEGIVFPFQLRYRGSEDGLPDALKTVLKWWAEGQAFMSGAASTGKGRFRMENAKYETLDLSDENQR<br>NDYLKNWGWRDEKGLEELKKRLNSGLPEPGNYRDPKWHEINVSIEMASPFINGDPIRAAVDKRGTDVVTFVKYKAEGEEAKPVCA<br>YKAESFRGVIRSAVARIHMEDGVPLTELTHSDCECLLCQIFGSEYEAGKIRFEDLVFESDPEPVTFDHVAIDRFTGGAADKKKFDDSP<br>LPGSPARPLMLKGSFWIRRDVLEDEEYCKALGKALADVNNGLYPLGGKSAIGYGQVKSLGIKGDDKRISRLMNPAFDETDVAVPEK<br>PKTDAEVRIEAEKVYYPHYFVEPHKKVEREEKPCGHQKFHEGRLTGKIRCKLITKTPLIVPDTSNDDFFRPADKEARKEKDEYHKSY<br>AFFRLHKQIMIPGSELRGMVSSVYETVTNSCFRIFDETKRLSWRMDADHQNVLQDFLPGRVTADGKHIQKFSETARVPFYDKTQKH |

| | | |
|---|---|---|
| | | FDILDEQEIAGEKPVRMWVKRFIKRLSLVDPAKHPQKKQDNKWKRRKEGIATFIEQKNGSYYFNVVTNNGCTSFHLWHKPDNFDQ<br>EKLEGIQNGEKLDCWVRDSRYQKAFQEIPENDPDGWECKEGYLHVVGPSKVEFSDKKGDVINNFQGTLPSVPNDWKTIRTNDFKN<br>RKRKNEPVFCCEDDKGNYYTMAKYCETFFFDLKENEEYEIPEKARIKYKELLRVYNNNPQAVPESVFQSRVARENVEKLKSGDLVY<br>FKHNEKYVEDIVPVRISRTVDDRMIGKRMSADLRPCHGDWVEDGDLSALNAYPEKRLLLRHPKGLCPACRLFGTGSYKGRVRFGF<br>ASLENDPEWLIPGKNPGDPFHGGPVMLSLLERPRPTWSIPGSDNKFKVPGRKFYVHHHAWKTIKDGNHPTTGKAIEQSPNNRTVEAL<br>AGGNSFSFEIAFENLKEWELGLLIHSLQLEKGLAHKLGMAKSMGFGSVEIDVESVRLRKDWKQWRNGNSEIPNWLGKGFAKLKEW<br>FRDELDFIENLKKLLWFPEGDQAPRVCYPMLRKKDDPNGNSGYEELKDGEFKKEDRQKKLTTPWTPWAMSLYDDLGVETSDSKTE<br>GWSKNFKLLQSQLQVKKAALTQAKSQRTKQSTVLAPVIDLKRGGSSDDRQIVDTPPHVAAGLKDPVPSGFSAGEVLIPLADEYDPM<br>FPNDYEKVVKRQREERQRQRELERQKEIEEREKRRKDRHEASGFARRPDPDSDEDEDYERERRKRSMGGAAIAPPTSLVEKDKELP<br>RDFPYEEDSRPRSQSSKAAIPPPVYEEQDRPRSPTGPSNSFLANMGGTVAHKIMQKYGFREGQGLGKHEQGLSTALSVEKTSKRGGK<br>IIVGDATEKDASKKSDSNPLTEILKCPTKVVLLRNMVGAGEVDEDLEVETKEECEKYGKVGKCVIFEIPGAPDDEAVRIFLEFERVES<br>AIKAVVDLNGRYFGGRVVKACFYNLDKFRVLDLAEQVKRTADGSEFESPKKKRKV |
| 15 | *Dis*Cas7-11 RBM17 C-terminal fusion, XTEN linker | MKRTADGSEFESPKKKRKVTTTMKISIEFLEPFRMTKWQESTRRNKNNKEFVRGQAFARWHRNKKDNTKGRPYITGTLLRSAVIRS<br>AENLLTLSDGKISEKTCCPGKFDTEDKDRLLQLRQRSTLRWTDKNPCPDNAETYCPFCELLGRSGNDGKKAEKKDWRFRIHFGNLS<br>LPGKPDFDGPKAIGSQRVLNRVDFKSGKAHDFFKAYEVDHTRFPRFEGEITIDNKVSAEARKLLCDSLKFTDRLCGALCVIRFDEYTP<br>AADSGKQTENVQAEPNANLAEKTAEQIISILDDNKKTEYTRLLADAIRSLRRSSKLVAGLPKDHDGKDDHYLWDIGKKKKDENSVT<br>IRQILTTSADTKELKNAGKWREFCEKLGEALYLKSKDMSGGLKITRRILGDAEFHGKPDRLEKSRSVSIGSVLKETVVCGELVAKTPF<br>FFGAIDEDAKQTDLQVLLTPDNKYRLPRSAVRGILRRDLQTYFDSPCNAELGGRPCMCKTCRIMRGITVMDARSEYNAPPEIRHRTRI<br>NPFTGTVAEGALFNMEVAPEGIVFPFQLRYRGSEDGLPDALKTVLKWWAEGQAFMSGAASTGKGRFRMENAKYETLDLSDENQR<br>NDYLKNWGWRDEKGLEELKKRLNSGLPEPGNYRDPKWHEINVSIEMASPFINGDPIRAAVDKRGTDVVTFVKYKAEGEEAKPVCA<br>YKAESFRGVIRSAVARIHMEDGVPLTELTHSDCECLLCQIFGSEYEAGKIRFEDLVFESDPEPVTFDHVAIDRFTGGAADKKKFDDSP<br>LPGSPARPLMLKGSFWIRRDVLEDEEYCKALGKALADVNNGLYPLGGKSAIGYGQVKSLGIKGDDKRISRLMNPAFDETDVAVPEK<br>PKTDAEVRIEAEKVYYPHYFVEPHKKVEREEKPCGHQKFHEGRLTGKIRCKLITKTPLIVPDTSNDDFFRPADKEARKEKDEYHKSY<br>AFFRLHKQIMIPGSELRGMVSSVYETVTNSCFRIFDETKRLSWRMDADHQNVLQDFLPGRVTADGKHIQKFSETARVPFYDKTQKH<br>FDILDEQEIAGEKPVRMWVKRFIKRLSLVDPAKHPQKKQDNKWKRRKEGIATFIEQKNGSYYFNVVTNNGCTSFHLWHKPDNFDQ<br>EKLEGIQNGEKLDCWVRDSRYQKAFQEIPENDPDGWECKEGYLHVVGPSKVEFSDKKGDVINNFQGTLPSVPNDWKTIRTNDFKN<br>RKRKNEPVFCCEDDKGNYYTMAKYCETFFFDLKENEEYEIPEKARIKYKELLRVYNNNPQAVPESVFQSRVARENVEKLKSGDLVY<br>FKHNEKYVEDIVPVRISRTVDDRMIGKRMSADLRPCHGDWVEDGDLSALNAYPEKRLLLRHPKGLCPACRLFGTGSYKGRVRFGF<br>ASLENDPEWLIPGKNPGDPFHGGPVMLSLLERPRPTWSIPGSDNKFKVPGRKFYVHHHAWKTIKDGNHPTTGKAIEQSPNNRTVEAL<br>AGGNSFSFEIAFENLKEWELGLLIHSLQLEKGLAHKLGMAKSMGFGSVEIDVESVRLRKDWKQWRNGNSEIPNWLGKGFAKLKEW<br>FRDELDFIENLKKLLWFPEGDQAPRVCYPMLRKKDDPNGNSGYEELKDGEFKKEDRQKKLTTPWTPWASGSETPGTSESATPESSL<br>YDDLGVETSDSKTEGWSKNFKLLQSQLQVKKAALTQAKSQRTKQSTVLAPVIDLKRGGSSDDRQIVDTPPHVAAGLKDPVPSGFSA<br>GEVLIPLADEYDPMFPNDYEKVVKRQREERQRQRELERQKEIEEREKRRKDRHEASGFARRPDPDSDEDEDYERERRKRSMGGAAI<br>APPTSLVEKDKELPRDFPYEEDSRPRSQSSKAAIPPPVYEEQDRPRSPTGPSNSFLANMGGTVAHKIMQKYGFREGQGLGKHEQGLS<br>TALSVEKTSKRGGKIIVGDATEKDASKKSDSNPLTEILKCPTKVVLLRNMVGAGEVDEDLEVETKEECEKYGKVGKCVIFEIPGAPD<br>DEAVRIFLEFERVESAIKAVVDLNGRYFGGRVVKACFYNLDKFRVLDLAEQVKRTADGSEFESPKKKRKV |
| 16 | *Dis*Cas7-11 SF3B6 N-terminal fusion | MKRTADGSEFESPKKKRKVMAMQAAKRANIRLPPEVNRILYIRNLPYKITAEEMYDIFGKYGPIRQIRVGNTPETRGTAYVVYEDIF<br>DAKNACDHLSGFNVCNRYLVVLYYNANRAFQKMDTKKKEEQLKLLKEKYGINTDPPKTTTMKISIEFLEPFRMTKWQESTRRNKN<br>NKEFVRGQAFARWHRNKKDNTKGRPYITGTLLRSAVIRSAENLLTLSDGKISEKTCCPGKFDTEDKDRLLQLRQRSTLRWTDKNPC<br>PDNAETYCPFCELLGRSGNDGKKAEKKDWRFRIHFGNLSLPGKPDFDGPKAIGSQRVLNRVDFKSGKAHDFFKAYEVDHTRFPRFE<br>GEITIDNKVSAEARKLLCDSLKFTDRLCGALCVIRFDEYTPAADSGKQTENVQAEPNANLAEKTAEQIISILDDNKKTEYTRLLADAI<br>RSLRRSSKLVAGLPKDHDGKDDHYLWDIGKKKKDENSVTIRQILTTSADTKELKNAGKWREFCEKLGEALYLKSKDMSGGLKITRR<br>ILGDAEFHGKPDRLEKSRSVSIGSVLKETVVCGELVAKTPFFFGAIDEDAKQTDLQVLLTPDNKYRLPRSAVRGILRRDLQTYFDSPC<br>NAELGGRPCMCKTCRIMRGITVMDARSEYNAPPEIRHRTRINPFTGTVAEGALFNMEVAPEGIVFPFQLRYRGSEDGLPDALKTVLK<br>WWAEGQAFMSGAASTGKGRFRMENAKYETLDLSDENQRNDYLKNWGWRDEKGLEELKKRLNSGLPEPGNYRDPKWHEINVSIE<br>MASPFINGDPIRAAVDKRGTDVVTFVKYKAEGEEAKPVCAYKAESFRGVIRSAVARIHMEDGVPLTELTHSDCECLLCQIFGSEYEA<br>GKIRFEDLVFESDPEPVTFDHVAIDRFTGGAADKKKFDDSPLPGSPARPLMLKGSFWIRRDVLEDEEYCKALGKALADVNNGLYPLG<br>GKSAIGYGQVKSLGIKGDDKRISRLMNPAFDETDVAVPEKPKTDAEVRIEAEKVYYPHYFVEPHKKVEREEKPCGHQKFHEGRLTG<br>KIRCKLITKTPLIVPDTSNDDFFRPADKEARKEKDEYHKSYAFFRLHKQIMIPGSELRGMVSSVYETVTNSCFRIFDETKRLSWRMDA<br>DHQNVLQDFLPGRVTADGKHIQKFSETARVPFYDKTQKHFDILDEQEIAGEKPVRMWVKRFIKRLSLVDPAKHPQKKQDNKWKRR<br>KEGIATFIEQKNGSYYFNVVTNNGCTSFHLWHKPDNFDQEKLEGIQNGEKLDCWVRDSRYQKAFQEIPENDPDGWECKEGYLHVV<br>GPSKVEFSDKKGDVINNFQGTLPSVPNDWKTIRTNDFKNRKRKNEPVFCCEDDKGNYYTMAKYCETFFFDLKENEEYEIPEKARIKY<br>KELLRVYNNNPQAVPESVFQSRVARENVEKLKSGDLVYFKHNEKYVEDIVPVRISRTVDDRMIGKRMSADLRPCHGDWVEDGDLS<br>ALNAYPEKRLLLRHPKGLCPACRLFGTGSYKGRVRFGFASLENDPEWLIPGKNPGDPFHGGPVMLSLLERPRPTWSIPGSDNKFKVP<br>GRKFYVHHHAWKTIKDGNHPTTGKAIEQSPNNRTVEALAGGNSFSFEIAFENLKEWELGLLIHSLQLEKGLAHKLGMAKSMGFGSV<br>EIDVESVRLRKDWKQWRNGNSEIPNWLGKGFAKLKEWFRDELDFIENLKKLLWFPEGDQAPRVCYPMLRKKDDPNGNSGYEELK<br>DGEFKKEDRQKKLTTPWTPWAKRTADGSEFESPKKKRKV |
| 17 | *Dis*Cas7-11 SF3B6 C-terminal fusion | MKRTADGSEFESPKKKRKVTTTMKISIEFLEPFRMTKWQESTRRNKNNKEFVRGQAFARWHRNKKDNTKGRPYITGTLLRSAVIRS<br>AENLLTLSDGKISEKTCCPGKFDTEDKDRLLQLRQRSTLRWTDKNPCPDNAETYCPFCELLGRSGNDGKKAEKKDWRFRIHFGNLS<br>LPGKPDFDGPKAIGSQRVLNRVDFKSGKAHDFFKAYEVDHTRFPRFEGEITIDNKVSAEARKLLCDSLKFTDRLCGALCVIRFDEYTP<br>AADSGKQTENVQAEPNANLAEKTAEQIISILDDNKKTEYTRLLADAIRSLRRSSKLVAGLPKDHDGKDDHYLWDIGKKKKDENSVT<br>IRQILTTSADTKELKNAGKWREFCEKLGEALYLKSKDMSGGLKITRRILGDAEFHGKPDRLEKSRSVSIGSVLKETVVCGELVAKTPF<br>FFGAIDEDAKQTDLQVLLTPDNKYRLPRSAVRGILRRDLQTYFDSPCNAELGGRPCMCKTCRIMRGITVMDARSEYNAPPEIRHRTRI<br>NPFTGTVAEGALFNMEVAPEGIVFPFQLRYRGSEDGLPDALKTVLKWWAEGQAFMSGAASTGKGRFRMENAKYETLDLSDENQR<br>NDYLKNWGWRDEKGLEELKKRLNSGLPEPGNYRDPKWHEINVSIEMASPFINGDPIRAAVDKRGTDVVTFVKYKAEGEEAKPVCA<br>YKAESFRGVIRSAVARIHMEDGVPLTELTHSDCECLLCQIFGSEYEAGKIRFEDLVFESDPEPVTFDHVAIDRFTGGAADKKKFDDSP<br>LPGSPARPLMLKGSFWIRRDVLEDEEYCKALGKALADVNNGLYPLGGKSAIGYGQVKSLGIKGDDKRISRLMNPAFDETDVAVPEK<br>PKTDAEVRIEAEKVYYPHYFVEPHKKVEREEKPCGHQKFHEGRLTGKIRCKLITKTPLIVPDTSNDDFFRPADKEARKEKDEYHKSY<br>AFFRLHKQIMIPGSELRGMVSSVYETVTNSCFRIFDETKRLSWRMDADHQNVLQDFLPGRVTADGKHIQKFSETARVPFYDKTQKH<br>FDILDEQEIAGEKPVRMWVKRFIKRLSLVDPAKHPQKKQDNKWKRRKEGIATFIEQKNGSYYFNVVTNNGCTSFHLWHKPDNFDQ<br>EKLEGIQNGEKLDCWVRDSRYQKAFQEIPENDPDGWECKEGYLHVVGPSKVEFSDKKGDVINNFQGTLPSVPNDWKTIRTNDFKN<br>RKRKNEPVFCCEDDKGNYYTMAKYCETFFFDLKENEEYEIPEKARIKYKELLRVYNNNPQAVPESVFQSRVARENVEKLKSGDLVY<br>FKHNEKYVEDIVPVRISRTVDDRMIGKRMSADLRPCHGDWVEDGDLSALNAYPEKRLLLRHPKGLCPACRLFGTGSYKGRVRFGF<br>ASLENDPEWLIPGKNPGDPFHGGPVMLSLLERPRPTWSIPGSDNKFKVPGRKFYVHHHAWKTIKDGNHPTTGKAIEQSPNNRTVEAL<br>AGGNSFSFEIAFENLKEWELGLLIHSLQLEKGLAHKLGMAKSMGFGSVEIDVESVRLRKDWKQWRNGNSEIPNWLGKGFAKLKEW<br>FRDELDFIENLKKLLWFPEGDQAPRVCYPMLRKKDDPNGNSGYEELKDGEFKKEDRQKKLTTPWTPWAMAMQAAKRANIRLPPE<br>VNRILYIRNLPYKITAEEMYDIFGKYGPIRQIRVGNTPETRGTAYVVYEDIFDAKNACDHLSGFNVCNRYLVVLYYNANRAFQKMD<br>TKKKEEQLKLLKEKYGINTDPPKKRTADGSEFESPKKKRKV |
| 18 | *Dis*Cas7-11 SF3B6 C-terminal fusion, XTEN linker | MKRTADGSEFESPKKKRKVTTTMKISIEFLEPFRMTKWQESTRRNKNNKEFVRGQAFARWHRNKKDNTKGRPYITGTLLRSAVIRS<br>AENLLTLSDGKISEKTCCPGKFDTEDKDRLLQLRQRSTLRWTDKNPCPDNAETYCPFCELLGRSGNDGKKAEKKDWRFRIHFGNLS<br>LPGKPDFDGPKAIGSQRVLNRVDFKSGKAHDFFKAYEVDHTRFPRFEGEITIDNKVSAEARKLLCDSLKFTDRLCGALCVIRFDEYTP<br>AADSGKQTENVQAEPNANLAEKTAEQIISILDDNKKTEYTRLLADAIRSLRRSSKLVAGLPKDHDGKDDHYLWDIGKKKKDENSVT<br>IRQILTTSADTKELKNAGKWREFCEKLGEALYLKSKDMSGGLKITRRILGDAEFHGKPDRLEKSRSVSIGSVLKETVVCGELVAKTPF<br>FFGAIDEDAKQTDLQVLLTPDNKYRLPRSAVRGILRRDLQTYFDSPCNAELGGRPCMCKTCRIMRGITVMDARSEYNAPPEIRHRTRI<br>NPFTGTVAEGALFNMEVAPEGIVFPFQLRYRGSEDGLPDALKTVLKWWAEGQAFMSGAASTGKGRFRMENAKYETLDLSDENQR |

| | | |
|---|---|---|
| | | NDYLKNWGWRDEKGLEELKKRLNSGLPEPGNYRDPKWHEINVSIEMASPFINGDPIRAAVDKRGTDVVTFVKYKAEGEEAKPVCA<br>YKAESFRGVIRSAVARIHMEDGVPLTELTHSDCECLLCQIFGSEYEAGKIRFEDLVFESDPEPVTFDHVAIDRFTGGAADKKKFDDSP<br>LPGSPARPLMLKGSFWIRRDVLEDEEYCKALGKALADVNNGLYPLGGKSAIGYGQVKSLGIKGDDKRISRLMNPAFDETDVAVPEK<br>PKTDAEVRIEAEKVYYPHYFVEPHKKVEREEKPCGHQKFHEGRLTGKIRCKLITKTPLIVPDTSNDDFFRPADKEARKEKDEYHKSY<br>AFFRLHKQIMIPGSELRGMVSSVYETVTNSCFRIFDETKRLSWRMDADHQNVLQDFLPGRVTADGKHIQKFSETARVPFYDKTQKH<br>FDILDEQEIAGEKPVRMWVKRFIKRLSLVDPAKHPQKKQDNKWKRRKEGIATFIEQKNGSYYFNVVTNNGCTSFHLWHKPDNFDQ<br>EKLEGIQNGEKLDCWVRDSRYQKAFQEIPENDPDGWECKEGYLHVVGPSKVEFSDKKGDVINNFQGTLPSVPNDWKTIRTNDFKN<br>RKRKNEPVFCCEDDKGNYYTMAKYCETFFFDLKENEEYEIPEKARIKYKELLRVYNNNPQAVPESVFQSRVARENVEKLKSGDLVY<br>FKHNEKYVEDIVPVRISRTVDDRMIGKRMSADLRPCHGDWVEDGDLSALNAYPEKRLLLRHPKGLCPACRLFGTGSYKGRVRFGF<br>ASLENDPEWLIPGKNPGDPFHGGPVMLSLLERPRPTWSIPGSDNKFKVPGRKFYVHHHAWKTIKDGNHPTTGKAIEQSPNNRTVEAL<br>AGGNSFSFEIAFENLKEWELGLLIHSLQLEKGLAHKLGMAKSMGFGSVEIDVESVRLRKDWKQWRNGNSEIPNWLGKGFAKLKEW<br>FRDELDFIENLKKLLWFPEGDQAPRVCYPMLRKKDDPNGNSGYEELKDGEFKKEDRQKKLTTPWTPWASGSETPGTSESATPESAM<br>QAAKRANIRLPPEVNRILYIRNLPYKITAEEMYDIFGKYGPIRQIRVGNTPETRGTAYVVYEDIFDAKNACDHLSGFNVCNRYLVVLY<br>YNANRAFQKMDTKKKEEQLKLLKEKYGINTDPPKKRTADGSEFESPKKKRKV |
| 19 | *Dis*Cas7-11 U2AF1 N-terminal fusion | MKRTADGSEFESPKKKRKVMAEYLASIFGTEKDKVNCSFYFKIGACRHGDRCSRLHNKPTFSQTIALLNIYRNPQNSSQSADGLRCA<br>VSDVEMQEHYDEFFEEVFTEMEEKYGEVEEMNVCDNLGDHLVGNVYVKFRREEDAEKAVIDLNNRWFNGQPIHAELSPVTDFREA<br>CCRQYEMGECTRGGFCNFMHLKPISRELRRELYGRRRKKHRSRSRSRERRSRSRDRGRGGGGGGGGGGGRERDRRRSRDRERSG<br>RFTTTMKISIEFLEPFRMTKWQESTRRNKNNKEFVRGQAFARWHRNKKDNTKGRPYITGTLLRSAVIRSAENLLTLSDGKISEKTCCP<br>GKFDTEDKDRLLQLRQRSTLRWTDKNPCPDNAETYCPFCELLGRSGNDGKKAEKKDWRFRIHFGNLSLPGKPDFDGPKAIGSQRVL<br>NRVDFKSGKAHDFFKAYEVDHTRFPRFEGEITIDNKVSAEARKLLCDSLKFTDRLCGALCVIRFDEYTPAADSGKQTENVQAEPNAN<br>LAEKTAEQIISILDDNKKTEYTRLLADAIRSLRRSSKLVAGLPKDHDGKDDHYLWDIGKKKKDENSVTIRQILTTSADTKELKNAGK<br>WREFCEKLGEALYLKSKDMSGGLKITRRILGDAEFHGKPDRLEKSRSVSIGSVLKETVVCGELVAKTPFFFGAIDEDAKQTDLQVLL<br>TPDNKYRLPRSAVRGILRRDLQTYFDSPCNAELGGRPCMCKTCRIMRGITVMDARSEYNAPPEIRHRTRINPFTGTVAEGALFNMEV<br>APEGIVFPFQLRYRGSEDGLPDALKTVLKWWAEGQAFMSGAASTGKGRFRMENAKYETLDLSDENQRNDYLKNWGWRDEKGLE<br>ELKKRLNSGLPEPGNYRDPKWHEINVSIEMASPFINGDPIRAAVDKRGTDVVTFVKYKAEGEEAKPVCAYKAESFRGVIRSAVARIH<br>MEDGVPLTELTHSDCECLLCQIFGSEYEAGKIRFEDLVFESDPEPVTFDHVAIDRFTGGAADKKKFDDSPLPGSPARPLMLKGSFWIR<br>RDVLEDEEYCKALGKALADVNNGLYPLGGKSAIGYGQVKSLGIKGDDKRISRLMNPAFDETDVAVPEKPKTDAEVRIEAEKVYYP<br>HYFVEPHKKVEREEKPCGHQKFHEGRLTGKIRCKLITKTPLIVPDTSNDDFFRPADKEARKEKDEYHKSYAFFRLHKQIMIPGSELRG<br>MVSSVYETVTNSCFRIFDETKRLSWRMDADHQNVLQDFLPGRVTADGKHIQKFSETARVPFYDKTQKHFDILDEQEIAGEKPVRMW<br>VKRFIKRLSLVDPAKHPQKKQDNKWKRRKEGIATFIEQKNGSYYFNVVTNNGCTSFHLWHKPDNFDQEKLEGIQNGEKLDCWVRD<br>SRYQKAFQEIPENDPDGWECKEGYLHVVGPSKVEFSDKKGDVINNFQGTLPSVPNDWKTIRTNDFKNRKRKNEPVFCCEDDKGNY<br>YTMAKYCETFFFDLKENEEYEIPEKARIKYKELLRVYNNNPQAVPESVFQSRVARENVEKLKSGDLVYFKHNEKYVEDIVPVRISRT<br>VDDRMIGKRMSADLRPCHGDWVEDGDLSALNAYPEKRLLLRHPKGLCPACRLFGTGSYKGRVRFGFASLENDPEWLIPGKNPGDP<br>FHGGPVMLSLLERPRPTWSIPGSDNKFKVPGRKFYVHHHAWKTIKDGNHPTTGKAIEQSPNNRTVEALAGGNSFSFEIAFENLKEWE<br>LGLLIHSLQLEKGLAHKLGMAKSMGFGSVEIDVESVRLRKDWKQWRNGNSEIPNWLGKGFAKLKEWFRDELDFIENLKKLLWFPE<br>GDQAPRVCYPMLRKKDDPNGNSGYEELKDGEFKKEDRQKKLTTPWTPWAKRTADGSEFESPKKKRKV |
| 20 | *Dis*Cas7-11 U2AF1 C-terminal fusion | MKRTADGSEFESPKKKRKVTTTMKISIEFLEPFRMTKWQESTRRNKNNKEFVRGQAFARWHRNKKDNTKGRPYITGTLLRSAVIRS<br>AENLLTLSDGKISEKTCCPGKFDTEDKDRLLQLRQRSTLRWTDKNPCPDNAETYCPFCELLGRSGNDGKKAEKKDWRFRIHFGNLS<br>LPGKPDFDGPKAIGSQRVLNRVDFKSGKAHDFFKAYEVDHTRFPRFEGEITIDNKVSAEARKLLCDSLKFTDRLCGALCVIRFDEYTP<br>AADSGKQTENVQAEPNANLAEKTAEQIISILDDNKKTEYTRLLADAIRSLRRSSKLVAGLPKDHDGKDDHYLWDIGKKKKDENSVT<br>IRQILTTSADTKELKNAGKWREFCEKLGEALYLKSKDMSGGLKITRRILGDAEFHGKPDRLEKSRSVSIGSVLKETVVCGELVAKTPF<br>FFGAIDEDAKQTDLQVLLTPDNKYRLPRSAVRGILRRDLQTYFDSPCNAELGGRPCMCKTCRIMRGITVMDARSEYNAPPEIRHRTRI<br>NPFTGTVAEGALFNMEVAPEGIVFPFQLRYRGSEDGLPDALKTVLKWWAEGQAFMSGAASTGKGRFRMENAKYETLDLSDENQR<br>NDYLKNWGWRDEKGLEELKKRLNSGLPEPGNYRDPKWHEINVSIEMASPFINGDPIRAAVDKRGTDVVTFVKYKAEGEEAKPVCA<br>YKAESFRGVIRSAVARIHMEDGVPLTELTHSDCECLLCQIFGSEYEAGKIRFEDLVFESDPEPVTFDHVAIDRFTGGAADKKKFDDSP<br>LPGSPARPLMLKGSFWIRRDVLEDEEYCKALGKALADVNNGLYPLGGKSAIGYGQVKSLGIKGDDKRISRLMNPAFDETDVAVPEK<br>PKTDAEVRIEAEKVYYPHYFVEPHKKVEREEKPCGHQKFHEGRLTGKIRCKLITKTPLIVPDTSNDDFFRPADKEARKEKDEYHKSY<br>AFFRLHKQIMIPGSELRGMVSSVYETVTNSCFRIFDETKRLSWRMDADHQNVLQDFLPGRVTADGKHIQKFSETARVPFYDKTQKH<br>FDILDEQEIAGEKPVRMWVKRFIKRLSLVDPAKHPQKKQDNKWKRRKEGIATFIEQKNGSYYFNVVTNNGCTSFHLWHKPDNFDQ<br>EKLEGIQNGEKLDCWVRDSRYQKAFQEIPENDPDGWECKEGYLHVVGPSKVEFSDKKGDVINNFQGTLPSVPNDWKTIRTNDFKN<br>RKRKNEPVFCCEDDKGNYYTMAKYCETFFFDLKENEEYEIPEKARIKYKELLRVYNNNPQAVPESVFQSRVARENVEKLKSGDLVY<br>FKHNEKYVEDIVPVRISRTVDDRMIGKRMSADLRPCHGDWVEDGDLSALNAYPEKRLLLRHPKGLCPACRLFGTGSYKGRVRFGF<br>ASLENDPEWLIPGKNPGDPFHGGPVMLSLLERPRPTWSIPGSDNKFKVPGRKFYVHHHAWKTIKDGNHPTTGKAIEQSPNNRTVEAL<br>AGGNSFSFEIAFENLKEWELGLLIHSLQLEKGLAHKLGMAKSMGFGSVEIDVESVRLRKDWKQWRNGNSEIPNWLGKGFAKLKEW<br>FRDELDFIENLKKLLWFPEGDQAPRVCYPMLRKKDDPNGNSGYEELKDGEFKKEDRQKKLTTPWTPWAMAEYLASIFGTEKDKVN<br>CSFYFKIGACRHGDRCSRLHNKPTFSQTIALLNIYRNPQNSSQSADGLRCAVSDVEMQEHYDEFFEEVFTEMEEKYGEVEEMNVCD<br>NLGDHLVGNVYVKFRREEDAEKAVIDLNNRWFNGQPIHAELSPVTDFREACCRQYEMGECTRGGFCNFMHLKPISRELRRELYGRR<br>RKKHRSRSRSRERRSRSRDRGRGGGGGGGGGGGRERDRRRSRDRERSGRFKRTADGSEFESPKKKRKV |
| 21 | *Dis*Cas7-11 U2AF1 C-terminal fusion, XTEN linker | MKRTADGSEFESPKKKRKVTTTMKISIEFLEPFRMTKWQESTRRNKNNKEFVRGQAFARWHRNKKDNTKGRPYITGTLLRSAVIRS<br>AENLLTLSDGKISEKTCCPGKFDTEDKDRLLQLRQRSTLRWTDKNPCPDNAETYCPFCELLGRSGNDGKKAEKKDWRFRIHFGNLS<br>LPGKPDFDGPKAIGSQRVLNRVDFKSGKAHDFFKAYEVDHTRFPRFEGEITIDNKVSAEARKLLCDSLKFTDRLCGALCVIRFDEYTP<br>AADSGKQTENVQAEPNANLAEKTAEQIISILDDNKKTEYTRLLADAIRSLRRSSKLVAGLPKDHDGKDDHYLWDIGKKKKDENSVT<br>IRQILTTSADTKELKNAGKWREFCEKLGEALYLKSKDMSGGLKITRRILGDAEFHGKPDRLEKSRSVSIGSVLKETVVCGELVAKTPF<br>FFGAIDEDAKQTDLQVLLTPDNKYRLPRSAVRGILRRDLQTYFDSPCNAELGGRPCMCKTCRIMRGITVMDARSEYNAPPEIRHRTRI<br>NPFTGTVAEGALFNMEVAPEGIVFPFQLRYRGSEDGLPDALKTVLKWWAEGQAFMSGAASTGKGRFRMENAKYETLDLSDENQR<br>NDYLKNWGWRDEKGLEELKKRLNSGLPEPGNYRDPKWHEINVSIEMASPFINGDPIRAAVDKRGTDVVTFVKYKAEGEEAKPVCA<br>YKAESFRGVIRSAVARIHMEDGVPLTELTHSDCECLLCQIFGSEYEAGKIRFEDLVFESDPEPVTFDHVAIDRFTGGAADKKKFDDSP<br>LPGSPARPLMLKGSFWIRRDVLEDEEYCKALGKALADVNNGLYPLGGKSAIGYGQVKSLGIKGDDKRISRLMNPAFDETDVAVPEK<br>PKTDAEVRIEAEKVYYPHYFVEPHKKVEREEKPCGHQKFHEGRLTGKIRCKLITKTPLIVPDTSNDDFFRPADKEARKEKDEYHKSY<br>AFFRLHKQIMIPGSELRGMVSSVYETVTNSCFRIFDETKRLSWRMDADHQNVLQDFLPGRVTADGKHIQKFSETARVPFYDKTQKH<br>FDILDEQEIAGEKPVRMWVKRFIKRLSLVDPAKHPQKKQDNKWKRRKEGIATFIEQKNGSYYFNVVTNNGCTSFHLWHKPDNFDQ<br>EKLEGIQNGEKLDCWVRDSRYQKAFQEIPENDPDGWECKEGYLHVVGPSKVEFSDKKGDVINNFQGTLPSVPNDWKTIRTNDFKN<br>RKRKNEPVFCCEDDKGNYYTMAKYCETFFFDLKENEEYEIPEKARIKYKELLRVYNNNPQAVPESVFQSRVARENVEKLKSGDLVY<br>FKHNEKYVEDIVPVRISRTVDDRMIGKRMSADLRPCHGDWVEDGDLSALNAYPEKRLLLRHPKGLCPACRLFGTGSYKGRVRFGF<br>ASLENDPEWLIPGKNPGDPFHGGPVMLSLLERPRPTWSIPGSDNKFKVPGRKFYVHHHAWKTIKDGNHPTTGKAIEQSPNNRTVEAL<br>AGGNSFSFEIAFENLKEWELGLLIHSLQLEKGLAHKLGMAKSMGFGSVEIDVESVRLRKDWKQWRNGNSEIPNWLGKGFAKLKEW<br>FRDELDFIENLKKLLWFPEGDQAPRVCYPMLRKKDDPNGNSGYEELKDGEFKKEDRQKKLTTPWTPWASGSETPGTSESATPESAE<br>YLASIFGTEKDKVNCSFYFKIGACRHGDRCSRLHNKPTFSQTIALLNIYRNPQNSSQSADGLRCAVSDVEMQEHYDEFFEEVFTEMEE<br>KYGEVEEMNVCDNLGDHLVGNVYVKFRREEDAEKAVIDLNNRWFNGQPIHAELSPVTDFREACCRQYEMGECTRGGFCNFMHLK<br>PISRELRRELYGRRRKKHRSRSRSRERRSRSRDRGRGGGGGGGGGGGRERDRRRSRDRERSGRFKRTADGSEFESPKKKRKV |
| 22 | *Dis*Cas7-11 U2AF2 C-terminal fusion | MKRTADGSEFESPKKKRKVTTTMKISIEFLEPFRMTKWQESTRRNKNNKEFVRGQAFARWHRNKKDNTKGRPYITGTLLRSAVIRS<br>AENLLTLSDGKISEKTCCPGKFDTEDKDRLLQLRQRSTLRWTDKNPCPDNAETYCPFCELLGRSGNDGKKAEKKDWRFRIHFGNLS<br>LPGKPDFDGPKAIGSQRVLNRVDFKSGKAHDFFKAYEVDHTRFPRFEGEITIDNKVSAEARKLLCDSLKFTDRLCGALCVIRFDEYTP<br>AADSGKQTENVQAEPNANLAEKTAEQIISILDDNKKTEYTRLLADAIRSLRRSSKLVAGLPKDHDGKDDHYLWDIGKKKKDENSVT<br>IRQILTTSADTKELKNAGKWREFCEKLGEALYLKSKDMSGGLKITRRILGDAEFHGKPDRLEKSRSVSIGSVLKETVVCGELVAKTPF |

| | | |
|---|---|---|
| | | FFGAIDEDAKQTDLQVLLTPDNKYRLPRSAVRGILRRDLQTYFDSPCNAELGGRPCMCKTCRIMRGITVMDARSEYNAPPEIRHRTRI<br>NPFTGTVAEGALFNMEVAPEGIVFPFQLRYRGSEDGLPDALKTVLKWWAEGQAFMSGAASTGKGRFRMENAKYETLDLSDENQR<br>NDYLKNWGWRDEKGLEELKKRLNSGLPEPGNYRDPKWHEINVSIEMASPFINGDPIRAAVDKRGTDVVTFVKYKAEGEEAKPVCA<br>YKAESFRGVIRSAVARIHMEDGVPLTELTHSDCECLLCQIFGSEYEAGKIRFEDLVFESDPEPVTFDHVAIDRFTGGAADKKKFDDSP<br>LPGSPARPLMLKGSFWIRRDVLEDEEYCKALGKALADVNNGLYPLGGKSAIGYGQVKSLGIKGDDKRISRLMNPAFDETDVAVPEK<br>PKTDAEVRIEAEKVYYPHYFVEPHKKVEREEKPCGHQKFHEGRLTGKIRCKLITKTPLIVPDTSNDDFFRPADKEARKEKDEYHKSY<br>AFFRLHKQIMIPGSELRGMVSSVYETVTNSCFRIFDETKRLSWRMDADHQNVLQDFLPGRVTADGKHIQKFSETARVPFYDKTQKH<br>FDILDEQEIAGEKPVRMWVKRFIKRLSLVDPAKHPQKKQDNKWKRRKEGIATFIEQKNGSYYFNVVTNNGCTSFHLWHKPDNFDQ<br>EKLEGIQNGEKLDCWVRDSRYQKAFQEIPENDPDGWECKEGYLHVVGPSKVEFSDKKGDVINNFQGTLPSVPNDWKTIRTNDFKN<br>RKRKNEPVFCCEDDKGNYYTMAKYCETFFFDLKENEEYEIPEKARIKYKELLRVYNNNPQAVPESVFQSRVARENVEKLKSGDLVY<br>FKHNEKYVEDIVPVRISRTVDDRMIGKRMSADLRPCHGDWVEDGDLSALNAYPEKRLLLRHPKGLCPACRLFGTGSYKGRVRFGF<br>ASLENDPEWLIPGKNPGDPFHGGPVMLSLLERPRPTWSIPGSDNKFKVPGRKFYVHHHAWKTIKDGNHPTTGKAIEQSPNNRTVEAL<br>AGGNSFSFEIAFENLKEWELGLLIHSLQLEKGLAHKLGMAKSMGFGSVEIDVESVRLRKDWKQWRNGNSEIPNWLGKGFAKLKEW<br>FRDELDFIENLKKLLWFPEGDQAPRVCYPMLRKKDDPNGNSGYEELKDGEFKKEDRQKKLTTPWTPWAMSDFDEFERQLNENKQE<br>RDKENRHRKRSHSRSRSRDRKRRSRSRDRRNRDQRSASRDRRRRSKPLTRGAKEEHGGLIRSPRHEKKKKVRKYWDVPPPGFEHIT<br>PMQYKAMQAAGQIPATALLPTMTPDGLAVTPTPVPVVGSQMTRQARRLYVGNIPFGITEEAMMDFFNAQMRLGGLTQAPGNPVLA<br>VQINQDKNFAFLEFRSVDETTQAMAFDGIIFQGQSLKIRRPHDYQPLPGMSENPSVYVPGVVSTVVPDSAHKLFIGGLPNYLNDDQV<br>KELLTSFGPLKAFNLVKDSATGLSKGYAFCEYVDINVTDQAIAGLNGMQLGDKKLLVQRASVGAKNATLVSPPSTINQTPVTLQVP<br>GLMSSQVQMGGHPTEVLCLMNMVLPEELLDDEEYEEIVEDVRDECSKYGLVKSIEIPRPVDGVEVPGCGKIFVEFTSVFDCQKAMQ<br>GLTGRKFANRVVVTKYCDPDSYHRRDFWKRTADGSEFESPKKKRKV |
| 23 | *Dis*Cas7-11 U2AF2 C-terminal fusion, XTEN linker | MKRTADGSEFESPKKKRKVTTTMKISIEFLEPFRMTKWQESTRRNKNNKEFVRGQAFARWHRNKKDNTKGRPYITGTLLRSAVIRS<br>AENLLTLSDGKISEKTCCPGKFDTEDKDRLLQLRQRSTLRWTDKNPCPDNAETYCPFCELLGRSGNDGKKAEKKDWRFRIHFGNLS<br>LPGKPDFDGPKAIGSQRVLNRVDFKSGKAHDFFKAYEVDHTRFPRFEGEITIDNKVSAEARKLLCDSLKFTDRLCGALCVIRFDEYTP<br>AADSGKQTENVQAEPNANLAEKTAEQIISILDDNKKTEYTRLLADAIRSLRRSSKLVAGLPKDHDGKDDHYLWDIGKKKKDENSVT<br>IRQILTTSADTKELKNAGKWREFCEKLGEALYLKSKDMSGGLKITRRILGDAEFHGKPDRLEKSRSVSIGSVLKETVVCGELVAKTPF<br>FFGAIDEDAKQTDLQVLLTPDNKYRLPRSAVRGILRRDLQTYFDSPCNAELGGRPCMCKTCRIMRGITVMDARSEYNAPPEIRHRTRI<br>NPFTGTVAEGALFNMEVAPEGIVFPFQLRYRGSEDGLPDALKTVLKWWAEGQAFMSGAASTGKGRFRMENAKYETLDLSDENQR<br>NDYLKNWGWRDEKGLEELKKRLNSGLPEPGNYRDPKWHEINVSIEMASPFINGDPIRAAVDKRGTDVVTFVKYKAEGEEAKPVCA<br>YKAESFRGVIRSAVARIHMEDGVPLTELTHSDCECLLCQIFGSEYEAGKIRFEDLVFESDPEPVTFDHVAIDRFTGGAADKKKFDDSP<br>LPGSPARPLMLKGSFWIRRDVLEDEEYCKALGKALADVNNGLYPLGGKSAIGYGQVKSLGIKGDDKRISRLMNPAFDETDVAVPEK<br>PKTDAEVRIEAEKVYYPHYFVEPHKKVEREEKPCGHQKFHEGRLTGKIRCKLITKTPLIVPDTSNDDFFRPADKEARKEKDEYHKSY<br>AFFRLHKQIMIPGSELRGMVSSVYETVTNSCFRIFDETKRLSWRMDADHQNVLQDFLPGRVTADGKHIQKFSETARVPFYDKTQKH<br>FDILDEQEIAGEKPVRMWVKRFIKRLSLVDPAKHPQKKQDNKWKRRKEGIATFIEQKNGSYYFNVVTNNGCTSFHLWHKPDNFDQ<br>EKLEGIQNGEKLDCWVRDSRYQKAFQEIPENDPDGWECKEGYLHVVGPSKVEFSDKKGDVINNFQGTLPSVPNDWKTIRTNDFKN<br>RKRKNEPVFCCEDDKGNYYTMAKYCETFFFDLKENEEYEIPEKARIKYKELLRVYNNNPQAVPESVFQSRVARENVEKLKSGDLVY<br>FKHNEKYVEDIVPVRISRTVDDRMIGKRMSADLRPCHGDWVEDGDLSALNAYPEKRLLLRHPKGLCPACRLFGTGSYKGRVRFGF<br>ASLENDPEWLIPGKNPGDPFHGGPVMLSLLERPRPTWSIPGSDNKFKVPGRKFYVHHHAWKTIKDGNHPTTGKAIEQSPNNRTVEAL<br>AGGNSFSFEIAFENLKEWELGLLIHSLQLEKGLAHKLGMAKSMGFGSVEIDVESVRLRKDWKQWRNGNSEIPNWLGKGFAKLKEW<br>FRDELDFIENLKKLLWFPEGDQAPRVCYPMLRKKDDPNGNSGYEELKDGEFKKEDRQKKLTTPWTPWASGSETPGTSESATPESSD<br>FDEFERQLNENKQERDKENRHRKRSHSRSRSRDRKRRSRSRDRRNRDQRSASRDRRRRSKPLTRGAKEEHGGLIRSPRHEKKKKVR<br>KYWDVPPPGFEHITPMQYKAMQAAGQIPATALLPTMTPDGLAVTPTPVPVVGSQMTRQARRLYVGNIPFGITEEAMMDFFNAQMR<br>LGGLTQAPGNPVLAVQINQDKNFAFLEFRSVDETTQAMAFDGIIFQGQSLKIRRPHDYQPLPGMSENPSVYVPGVVSTVVPDSAHKL<br>FIGGLPNYLNDDQVKELLTSFGPLKAFNLVKDSATGLSKGYAFCEYVDINVTDQAIAGLNGMQLGDKKLLVQRASVGAKNATLVSP<br>PSTINQTPVTLQVPGLMSSQVQMGGHPTEVLCLMNMVLPEELLDDEEYEEIVEDVRDECSKYGLVKSIEIPRPVDGVEVPGCGKIFVE<br>FTSVFDCQKAMQGLTGRKFANRVVVTKYCDPDSYHRRDFWKRTADGSEFESPKKKRKV |

<u>**Supplementary Table 4: RT, NGS and qRT-PCR gene-specific primers.**</u>

| | | |
|---|---|---|
| 1 | ACTB qPCR Primer Forward | catgtacgttgctatccaggc |
| 2 | ACTB qPCR Primer Rev | ctccttaatgtcacgcacgat |
| 3 | PABPC1 RT primer | aacagttggaacaccgg |
| 4 | PABPC1 NGS primer 1 Forward | ACACTCTTTCCCTACACGACGCTCTTCCGATCTcccctcctcaagagcaaaagcaaatgttggg |
| 5 | PABPC1 NGS primer 2 Forward | ACACTCTTTCCCTACACGACGCTCTTCCGATCTAcccctcctcaagagcaaaagcaaatgttggg |
| 6 | PABPC1 NGS primer 3 Forward | ACACTCTTTCCCTACACGACGCTCTTCCGATCTGAcccctcctcaagagcaaaagcaaatgttggg |
| 7 | PABPC1 NGS primer 4 Forward | ACACTCTTTCCCTACACGACGCTCTTCCGATCTTGAcccctcctcaagagcaaaagcaaatgttggg |
| 8 | PABPC1 NGS primer Reverse | GTGACTGGAGTTCAGACGTGTGCTCTTCCGATCcttcggtgaagcacaagtttcttttcatggtccc |
| 9 | PPIB RT primer | cttctccacctcgatcttg |
| 10 | PPIB NGS primer 1 Forward | ACACTCTTTCCCTACACGACGCTCTTCCGATCTctgaagcactacgggcctggctg |
| 11 | PPIB NGS primer 2 Forward | ACACTCTTTCCCTACACGACGCTCTTCCGATCTActgaagcactacgggcctggctg |
| 12 | PPIB NGS primer 3 Forward | ACACTCTTTCCCTACACGACGCTCTTCCGATCTGActgaagcactacgggcctggctg |
| 13 | PPIB NGS primer 4 Forward | ACACTCTTTCCCTACACGACGCTCTTCCGATCTTGActgaagcactacgggcctggctg |
| 14 | PPIB NGS primer Reverse | GTGACTGGAGTTCAGACGTGTGCTCTTCCGATCcagtctgcgatgatcacatccttcagggg |
| 15 | PPIB qPCR Primer Fw | ggcctggctgggtgagcatg |
| 16 | PPIB qPCR Primer Rev | agggcttctccacctcgatcttgc |
| 17 | RPL41 RT primer Forward | cttggacctctgcctcat |
| 18 | RPL41 NGS primer 1 Forward | ACACTCTTTCCCTACACGACGCTCTTCCGATCTctctcggccttagcgccatttttttgg |
| 19 | RPL41 NGS primer 2 Forward | ACACTCTTTCCCTACACGACGCTCTTCCGATCTActctcggccttagcgccatttttttgg |
| 20 | RPL41 NGS primer 3 Forward | ACACTCTTTCCCTACACGACGCTCTTCCGATCTGActctcggccttagcgccatttttttgg |
| 21 | RPL41 NGS primer 4 Forward | ACACTCTTTCCCTACACGACGCTCTTCCGATCTTGActctcggccttagcgccatttttttgg |
| 22 | RPL41 NGS primer Reverse | GTGACTGGAGTTCAGACGTGTGCTCTTCCGATCtatgagcaaggtgggtctcagaggtgatcg |

| | | |
|---|---|---|
| 23 | SHANK3 RT primer Forward | agacttggtccgtggaatc |
| 24 | SHANK3 NGS primer 1 Forward | ACACTCTTTCCCTACACGACGCTCTTCCGATCTgctgcggccagacatcgc |
| 25 | SHANK3 NGS primer 2 Forward | ACACTCTTTCCCTACACGACGCTCTTCCGATCTAgctgcggccagacatcgc |
| 26 | SHANK3 NGS primer 3 Forward | ACACTCTTTCCCTACACGACGCTCTTCCGATCTGAgctgcggccagacatcgc |
| 27 | SHANK3 NGS primer 4 Forward | ACACTCTTTCCCTACACGACGCTCTTCCGATCTTGAgctgcggccagacatcgc |
| 28 | SHANK3 NGS primer Reverse | GTGACTGGAGTTCAGACGTGTGCTCTTCCGATCgagcccggcagcttctctgg |
| 29 | SHANK3 qPCR Primer Fw | agaagctggacgagatgctggc |
| 30 | SHANK3 qPCR Primer Rev | ggagcccggcagcttctct |
| 31 | STAT3 RT primer | atagttgaaatcaaagtcatcctg |
| 32 | STAT3 NGS primer 1 Forward | ACACTCTTTCCCTACACGACGCTCTTCCGATCTccaggccaaccaccccacag |
| 33 | STAT3 NGS primer 2 Forward | ACACTCTTTCCCTACACGACGCTCTTCCGATCTAccaggccaaccaccccacag |
| 34 | STAT3 NGS primer 3 Forward | ACACTCTTTCCCTACACGACGCTCTTCCGATCTGAccaggccaaccaccccacag |
| 35 | STAT3 NGS primer 4 Forward | ACACTCTTTCCCTACACGACGCTCTTCCGATCTTGAccaggccaaccaccccacag |
| 36 | STAT3 NGS primer Reverse | GTGACTGGAGTTCAGACGTGTGCTCTTCCGATCgttgaaatcaaagtcatcctggagattctctaccact |
| 37 | TOP2A RT primer | gattccttcagcaccatttatc |
| 38 | TOP2A NGS primer 1 Forward | ACACTCTTTCCCTACACGACGCTCTTCCGATCTggtcagtttggtaccaggctacatggtgg |
| 39 | TOP2A NGS primer 2 Forward | ACACTCTTTCCCTACACGACGCTCTTCCGATCTAggtcagtttggtaccaggctacatggtgg |
| 40 | TOP2A NGS primer 3 Forward | ACACTCTTTCCCTACACGACGCTCTTCCGATCTGAggtcagtttggtaccaggctacatggtgg |
| 41 | TOP2A NGS primer 4 Forward | ACACTCTTTCCCTACACGACGCTCTTCCGATCTTGAggtcagtttggtaccaggctacatggtgg |
| 42 | TOP2A NGS primer Reverse | GTGACTGGAGTTCAGACGTGTGCTCTTCCGATCccattcaggctcaacacgctggttgtc |

| | | |
|---|---|---|
| 43 | TOP2A qPCR Primer Fw | tcaatttggctcagaattttgtgggtagcaa |
| 44 | TOP2A qPCR Primer Rev | cattcaggctcaacacgctggttgtc |
| 45 | USF1 RT primer | tcgaagcacgtcattgtc |
| 46 | USF1 NGS primer 1 Forward | ACACTCTTTCCCTACACGACGCTCTTCCGATCTgccgccgagacaagatcaacaactgg |
| 47 | USF1 NGS primer 2 Forward | ACACTCTTTCCCTACACGACGCTCTTCCGATCTAgccgccgagacaagatcaacaactgg |
| 48 | USF1 NGS primer 3 Forward | ACACTCTTTCCCTACACGACGCTCTTCCGATCTGAgccgccgagacaagatcaacaactgg |
| 49 | USF1 NGS primer 4 Forward | ACACTCTTTCCCTACACGACGCTCTTCCGATCTTGAgccgccgagacaagatcaacaactgg |
| 50 | USF1 NGS primer Reverse | GTGACTGGAGTTCAGACGTGTGCTCTTCCGATCgctgcagttggtcaagtccctgcag |
| 51 | HTT 5' RT primer | tgacagactgtgccact |
| 52 | HTT 5' NGS primer 1 Forward | ACACTCTTTCCCTACACGACGCTCTTCCGATCTccggctgtggctgaggagc |
| 53 | HTT 5' NGS primer 2 Forward | ACACTCTTTCCCTACACGACGCTCTTCCGATCTAccggctgtggctgaggagc |
| 54 | HTT 5' NGS primer 3 Forward | ACACTCTTTCCCTACACGACGCTCTTCCGATCTGAccggctgtggctgaggagc |
| 55 | HTT 5' NGS primer Reverse | GTGACTGGAGTTCAGACGTGTGCTCTTCCGATCgactgtgccactatgttttcacatattgtcagacaatga |
| 56 | RPL41 5' RT primer | tctagacagtagcatgcagt |
| 57 | RPL41 5' NGS primer 1 Forward | ACACTCTTTCCCTACACGACGCTCTTCCGATCTagagccaagtggaggaagaagcgaat |
| 58 | RPL41 5' NGS primer 2 Forward | ACACTCTTTCCCTACACGACGCTCTTCCGATCTAagagccaagtggaggaagaagcgaat |
| 59 | RPL41 5' NGS primer 3 Forward | ACACTCTTTCCCTACACGACGCTCTTCCGATCTGAagagccaagtggaggaagaagcgaat |
| 60 | RPL41 5' NGS primer 4 Forward | ACACTCTTTCCCTACACGACGCTCTTCCGATCTTGAagagccaagtggaggaagaagcgaat |
| 61 | RPL41 5' NGS primer Reverse | GTGACTGGAGTTCAGACGTGTGCTCTTCCGATCcaacttgtaccagcatccccagcgtc |
| 62 | PABPC1 5' RT primer | gaatatgttgcctactccactt |

| | | |
|---|---|---|
| 63 | PABPC1 5' NGS primer 1 Forward | ACACTCTTTCCCTACACGACGCTCTTCCGATCTgacatgatcacccgccgctccttg |
| 64 | PABPC1 5' NGS primer 2 Forward | ACACTCTTTCCCTACACGACGCTCTTCCGATCTAgacatgatcacccgccgctccttg |
| 65 | PABPC1 5' NGS primer 3 Forward | ACACTCTTTCCCTACACGACGCTCTTCCGATCTGAgacatgatcacccgccgctccttg |
| 66 | PABPC1 5' NGS primer 4 Forward | ACACTCTTTCCCTACACGACGCTCTTCCGATCTTGAgacatgatcacccgccgctccttg |
| 67 | PABPC1 5' NGS primer Reverse | GTGACTGGAGTTCAGACGTGTGCTCTTCCGATCgcgaagtgatggatcacgctgagacc |
| 68 | PPIB 5' RT primer | ccaaatcctttctctcctgt |
| 69 | PPIB 5' NGS primer 1 Forward | ACACTCTTTCCCTACACGACGCTCTTCCGATCTatcgcggggtccgtcttcttcc |
| 70 | PPIB 5' NGS primer 2 Forward | ACACTCTTTCCCTACACGACGCTCTTCCGATCTAatcgcggggtccgtcttcttcc |
| 71 | PPIB 5' NGS primer 3 Forward | ACACTCTTTCCCTACACGACGCTCTTCCGATCTGAatcgcggggtccgtcttcttcc |
| 72 | PPIB 5' NGS primer 4 Forward | ACACTCTTTCCCTACACGACGCTCTTCCGATCTTGAatcgcggggtccgtcttcttcc |
| 73 | PPIB 5' NGS primer Reverse | GTGACTGGAGTTCAGACGTGTGCTCTTCCGATCggccacaaaattatccactgtttttggaacagtctttcc |
